# HyperCas12a enables multiplexed CRISPRi screens

**DOI:** 10.1101/2024.07.08.602263

**Authors:** Schuyler M. Melore, Christian McRoberts Amador, Marisa C. Hamilton, Charles A. Gersbach, Timothy E. Reddy

**Affiliations:** University Program in Genetics & Genomics, Duke University, Durham, NC, USA; Department of Biostatistics & Bioinformatics, Duke University School of Medicine, Durham, NC, USA; Center for Advanced Genomic Technologies, Duke University, Durham, NC, USA; Center for Combinatorial Gene Regulation, Duke University, Durham, NC, USA; Department of Pharmacology & Cancer Biology, Duke University School of Medicine, Durham, NC, USA; Department of Biomedical Engineering, Duke University, Durham, NC, USA

## Abstract

Interactions between multiple genes or cis-regulatory elements (CREs) underlie a wide range of biological processes in both health and disease. High-throughput screens using dCas9 fused to epigenome editing domains have allowed researchers to assess the impact of activation or repression of both coding and non-coding genomic regions on a phenotype of interest, but assessment of genetic interactions between those elements has been limited to pairs. Here, we combine a hyper-efficient version of *Lachnospiraceae bacterium* dCas12a (dHyperLbCas12a) with RNA Polymerase II expression of long CRISPR RNA (crRNA) arrays to enable efficient highly-multiplexed epigenome editing. We demonstrate that this system is compatible with several activation and repression domains, including the P300 histone acetyltransferase domain and SIN3A interacting domain (SID). We further show that the system can be used in cultured primary immune cells and to drive differentiation of induced pluripotent stem cells. We also developed new approaches to use the dCas12a platform for simultaneous activation and repression from a single crRNA array via co-expression of multiple dCas12a orthologues. Lastly, we demonstrate that the dHyperLbCas12a effectors are highly effective for multiple modalities of high-throughput screens, namely proliferation screens and screens to dissect the independent and combinatorial contributions of CREs on gene expression. The tools and methods introduced here create new possibilities for highly multiplexed control of gene expression in a wide variety of biological systems.

## Introduction

Many biological systems depend on coordinated activity between genetic elements. In mammalian systems, it is common that several cis-regulatory elements (CREs) control the expression of the same gene both at steady state and in response to environmental conditions^1–10^. Recent work to experimentally test and validate thousands of CRE-gene pairs found between two and three CREs per gene, while predictive modeling of millions of CRE-promoter interactions estimates about four CREs per gene^11,12^. It is also common that coordinated expression of several genes controls aspects of cell differentiation and disease^13–28^.

One way to evaluate the function of combinations of gene regulatory elements is to perturb their function *in vitro* or *in vivo*. Several technologies for genetic or epigenetic perturbation of regulatory elements have been developed based on bacterial clustered regularly interspaced short palindromic repeats (CRISPR) systems. Briefly, bacterial CRISPR systems typically use short CRISPR RNAs (crRNAs) to target a Cas nuclease to specific complementary ∼20 bp DNA sequences^29–33^. Those systems have been adapted for gene activation and repression by fusing an epigenome-modifying domain to nuclease-inactivated “dead” dCas proteins^34,35^. Moreover, by delivering multiple crRNAs at once, it is possible to study interactions between CREs. For example, dCas9-based studies of pairwise epigenetic perturbations have revealed combinations of transcription factors that enhance neuronal differentiation^13^; long-distance synergistic interactions between enhancers controlling MYC expression^2^; synthetic lethality in the human genome^27^; and hierarchies in CRE interactions^19^.

Scaling beyond pairwise or three-way combinations of sgRNAs is challenging, however, for several reasons. Cas9 requires an RNA molecule known as a trans-activating crRNA (tracrRNA) to bind the crRNA. To simplify studies using Cas9, the tracrRNA and crRNA are commonly expressed as a single ∼100 bp molecule known as a single guide RNA or sgRNA^30,31^. However, repetitive sequences in multi-sgRNA plasmids, including the tracrRNA, can also lead to extensive plasmid recombination^36–39^. Creating further design constraints, each sgRNA must either be expressed from its own promoter, or expressed as part of a longer sgRNA array that is cleaved via separately expressed enzymes^40^. Together, those complexities severely limit highly multiplexed Cas9 genome- or epigenome-editing screens.

In contrast, several features of the Cas12a nuclease make it ideally suited for highly multiplexed manipulation of gene regulatory elements. Like Cas9, deoxyribonuclease-inactivated dCas12a can be fused with epigenetic effectors to activate or repress gene expression^25,41–44^. Unlike Cas9, however, a ∼40 bp crRNA is sufficient to target Cas12a or dCas12a to sites in the human genome^32,33^. Cas12a also has an endoribonuclease activity that allows it to cleave individual crRNAs from a single pre-crRNA molecule containing a tandem array of many crRNAs^32^. Using those features, previous studies have targeted 20 genomic sites at once via a single pre-crRNA^44^. Several studies have also used that multiplexing capability in high-throughput genome-editing screens to identify synthetic lethal combinations of human genes; and in an epigenome-editing screen to identify combinations of gene regulatory elements contributing to cell fitness^10,21,23–25,45^.

Here, we introduce a suite of novel dCas12a epigenome editors for effective, highly-multiplexed gene activation and repression; and demonstrate that those effectors together with additional advances enable multiplexed high-throughput screens of combinatorial regulatory element activity. To do so, we systematically compared epigenome editing activity across a wide range of existing and novel dCas12a epigenome effectors. We identify dCas12a effectors sensitive enough to alter regulatory element activity when integrated into the genome by lentiviral transduction and show that these effectors can be used to modulate gene expression in cultured primary cells. We also demonstrate expression of complex pre-crRNAs that include selectable markers and up to 14 crRNAs from an RNA polymerase II promoter. Further, by combining crRNAs for different dCas12a proteins in a single pre-crRNA, we can simultaneously activate and repress gene expression. We further show that dCas12a repressors can be successfully used in low MOI fitness screens. Together, those advances enable highly-multiplexed high-throughput epigenome editing screens, which we demonstrate by dissecting the cis-regulatory architecture for a dose-dependent gene expression response to glucocorticoids.

## Results

### dHyperLbCas12a and dEnAsCas12a outperform other dCas12a variants for epigenome editing

To optimize the dCas12a epigenome editing system for high-throughput screening, we first evaluated which Cas12a variants are most effective for epigenome editing. In addition to the wild-type Cas12a from *Acidaminococcus* sp. BV3L6 (As) and *Lachnospiraceae bacterium* (Lb), several engineered Cas12a versions with improved activity have been recently published^46–48^. However, not all these variants have been used for epigenome editing, nor has their activation and repression strength been compared head-to-head in a single study. We systematically compared the strength of gene activation and gene repression across five dCas12a variants: the wild type, enhanced, and ultra versions of dAsCas12a; and the wild type and hyper versions of dLbCas12a. For activation, we fused each dCas12a variant to the tripartite synthetic activator VPR (VP64-p65-Rta), while for repression we fused each dCas12a variant to a Krüppel-associated box (KRAB) domain from KOX1. We cloned each of the resulting dCas12a effector proteins into the same plasmid backbone where they are expressed co-transcriptionally with a green fluorescent protein (EGFP). We then evaluated the effects on gene expression by co-transfecting each dCas12a effector into HEK293T cells along with a pre-crRNA array encoding three crRNAs targeting a gene of interest, or three non-targeting control crRNAs. Because Lb and AsCas12a have the same PAM sequence (TTTV), we used the same crRNA target sequences throughout, enabling well-controlled comparisons between As and Lb-derived Cas12a proteins. We then assessed the impact of each Cas12a variant on target gene expression 48 hours later using RT-qPCR **(Fig. 1a-b)**.

**Figure 1.**
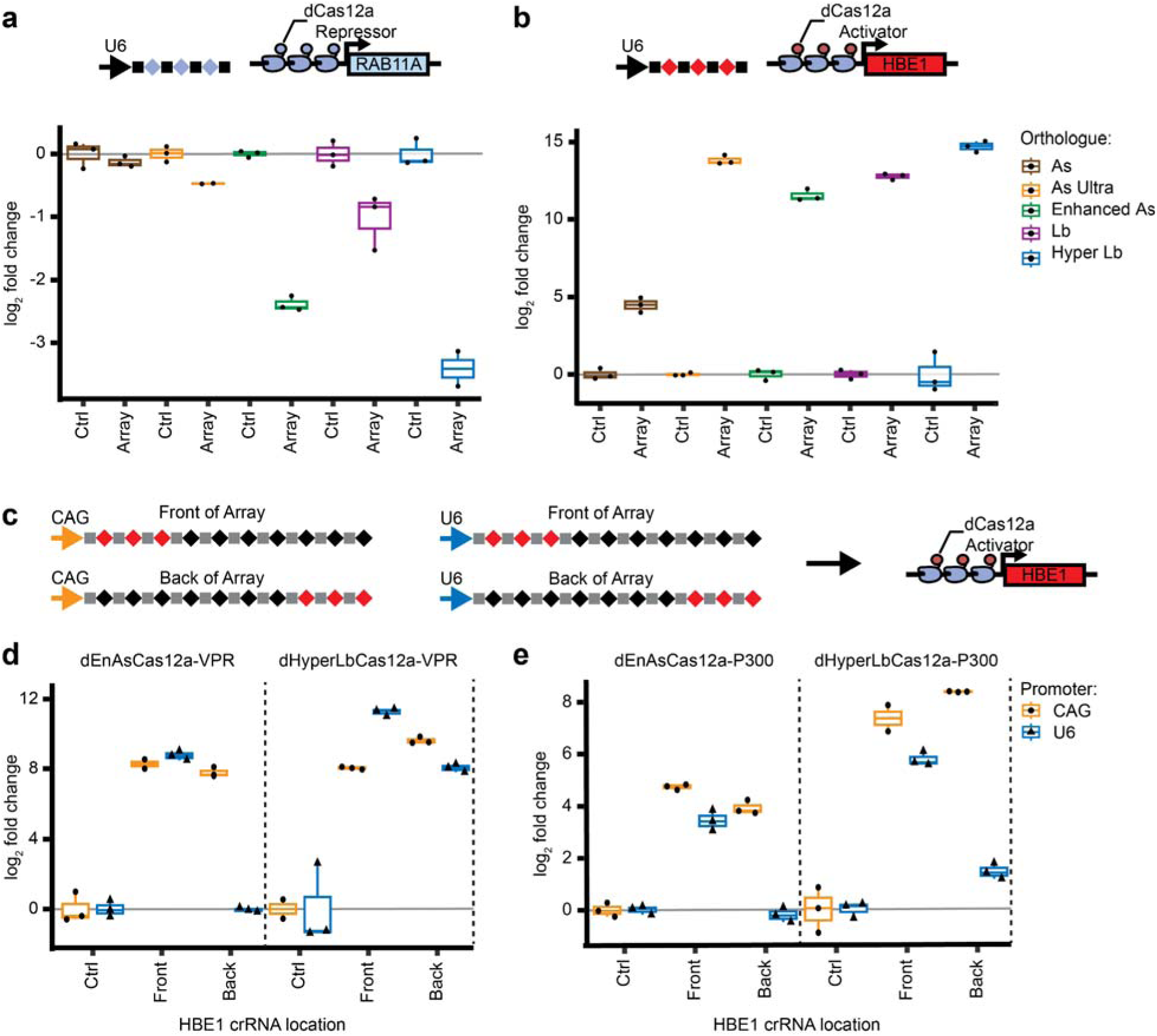
Optimization of the dCas12a system for multiplexed epigenome editing. **(a)** Comparison of dCas12a repressors. Five different Cas12a variants were fused to the transcriptional repressor KRAB. Each dCas12a-KRAB construct was co-transfected into HEK293T cells with either a pre-crRNA of three crRNAs targeting the *RAB11A* promoter (nTPM = 295.3)^50^ or a control pre-crRNA containing three non-targeting crRNAs. After two days, cells were sorted on Cas12a expression using a GFP marker, mRNA was harvested and relative *RAB11A* expression levels were quantified using RT-qPCR (two or more independent experiments per transfection condition) **(b)** Comparison of dCas12a activators. Five different Cas12a variants were fused to the transcriptional activator VPR. Each dCas12a-VPR construct was co-transfected into HEK293T cells with either a pre-crRNA of three crRNAs targetin the *HBE1* promoter (nTPM = 0.1) or a control pre-crRNA containing three non-targeting crRNAs. After 2 days, mRNA was harvested and relative *HBE1* expression levels were quantified using RT-qPCR (n = 3 independent experiments per transfection condition) **(c)** Constructs used to assess different promoters for pre-crRNAs. Pre-crRNAs of 10 crRNAs, driven by CAG or U6, were designed to contain three crRNA targeting the *HBE1* promoter in either position 1-3 (Front of array) or 8-10 (Back of array). These constructs were then co-transfected into HEK293T cells with either **(d)** dCas12a-VPR or **(e)** dCas12a-P300. After 2 days, mRNA was harvested and relative *HBE1* expression levels were quantified using RT-qPCR (two or more independent experiments per transfection condition)

The dHyperLbCas12a had the strongest effects for both activation and repression, though dEnAsCas12a showed consistently strong activity as well. EnAsCas12a has the additional benefit of recognizing several non-canonical PAM sequences, with an estimated 7-fold increase in target sites in the human genome over wild type Lb and AsCas12a^46^. Our results also replicated previous studies showing that wild type AsCas12a has lower epigenome editing activity than wild type LbCas12a, but that the engineered dAsCas12a variants have comparable activity^41,46^. Due to claims that HyperLbCas12a most strongly outperforms other Cas12a variants under low crRNA conditions^48^, we also tested the activators in a “low crRNA’’ condition where we transfected 1/10th the amount of pre-crRNA array as in **Fig. 1a**. Under those conditions, dHyperLbCas12a had the strongest gene activation as well **(Supp. Fig. 1a)**. The dCas12 constructs we evaluated all have a C terminal nuclear localization signal (NLS). We also evaluated adding an additional NLS to the N-terminus of the dCas12a protein, but found no increase in gene expression **(Supp. Fig. 1b)**. Taken together, we focused on developing multiplexed high-throughput screening using dHyperLbCas12a due to its high overall activity, and dEnAsCas12a due to its expanded targeting range.

### Improved dCas12a pre-crRNA expression for long arrays via an RNA Pol II promoter

Next, we focused on optimizing the Cas12a array expression system. In bacteria, RNA polymerase expresses Cas12a pre-crRNA arrays without a 5’ cap or poly-A tail; and synthetic mammalian systems recreate that structure using RNA Pol III. Expression with RNA Pol III limits the length of the pre-crRNA, however. For example, a recent study showed that U6-driven pre-crRNA expression drops off after ∼4 crRNAs^23^. Because our goal is to enable highly-multiplexed screening involving arrays of >4 crRNAs, we expected that driving array expression with RNA Pol III would be insufficient and we would need to use a different expression system.

We hypothesized that an RNA pol II promoter would effectively drive longer arrays due to the higher processivity of RNA pol II compared to RNA pol III. To test this, we created pre-crRNA arrays containing 10 crRNAs wherein we placed three crRNAs targeting the *HBE1* promoter either at the front or back of the array. We then expressed the pre-crRNAs via the RNA pol III promoter U6 or the RNA pol II promoter CAG, which was chosen due to its consistently high transcription across many cell types and DNA delivery mechanisms^49^ (**Fig. 1c)**. We then co-transfected the pre-crRNA arrays along with plasmids expressing variants of dCas12a-VPR or dCas12a-P300 (described below) into HEK293T cells. As negative controls, we used pre-crRNA arrays containing ten non-targeting crRNAs **(Fig. 1d-e)**. When we used the U6 promoter to express the pre-crRNA arrays, all dCas12a variants robustly activated *HBE1* expression when *HBE1*-targeting crRNAs were at the front of the array; but there was little or no *HBE1* activation for three of the four dCas12a epigenome activators when we moved the *HBE1*-targeting crRNAs to the back of the array. Expressing the same pre-crRNA arrays using the CAG promoter rescued *HBE1* activation from the back of the pre-crRNA array for all dCas12a variants tested. There was also no substantial difference in *HBE1* activation from the front or back of the pre-crRNA array. Together, those results indicate that RNA Pol II improves expression of long pre-crRNA arrays.

### dCas12a-SID fusions enable multiplexed gene repression

To date, dCas12a-based repression studies have largely used the KRAB repressor domain^10,42,44,48^. Though KRAB is a potent repressor of transcription, there are potential advantages for repressing via other effector domains. First, KRAB is a potent repressor that recruits KAP1 and induces heterochromatin^51^. In some cases, more subtle repression may be more relevant to model disease. Second, there are also reports that heterochromatin formation can spread over long distances in some cases, causing potential off target effects. For example, targeting KRAB to an enhancer element can repress multiple enhancers or gene promoters nearby in some cases^52^, but not always^53,54^. Third, there are cases when dCas12a-KRAB fusions have little to no effect on gene expression, and alternative effectors may be able to repress those genes^10,44,55^.

To develop additional dCas12a-based repressors, we created and tested dCas12a fusions to four copies of the SIN3A interacting domain (SID) derived from MAD1. The SID domain mediates gene repression through recruitment of SIN3A and the histone deacetylases HDAC1 and HDAC2^56^. Previous studies have also shown that SID fusion proteins can repress enhancers and promoters^4,57^, and we sought here to extend those findings to dCas12a fusions.

We first compared fusions of KRAB, 3xKRAB, 4xSID, and both 3xKRAB and 4xSID to dLbCas12a or dEnAsCas12a. We co-transfected plasmids encoding those fusion proteins and a pre-crRNA containing three crRNAs targeting the *VEGFA* or *RAB5A* (one spacer sequence originally reported in Ref. 44) promoter into HEK293T cells. We then evaluated changes in *VEGFA* and *RAB5A* expression relative to cells expressing the same fusion protein but a control pre-crRNA array targeting an unrelated gene (**Supp. Fig 2a-b**). The C-terminal fusions of KRAB and 4xSID had the strongest overall repressive effect for both dLbCas12a and EnAsCas12a, demonstrating the dCas12a-4xSID fusions can repress gene expression.

We further evaluated dCas12a-4xSID repression using multiple crRNAs targeted to the promoter of individual genes. We cloned dEnAsCas12a and dHyperLbCas12a repressor fusions into a plasmid backbone with a stronger promoter and a strong NLS^48^ **(Fig. 2a)**. We then tested the efficacy of these repressors in HEK293T cells using a pre-crRNA containing three crRNAs targeting *VEGFA* or *RAB11A* **(Fig. 2b)**. All repressors tested were able to induce significant and robust gene repression, with dHyperLbCas12a repressor fusions achieving >50% repression for both gene targets (p < 0.005, FDR-adjusted Student’s T-test) **(Fig. 2c-d, and Fig. 1)**.

**Figure 2.**
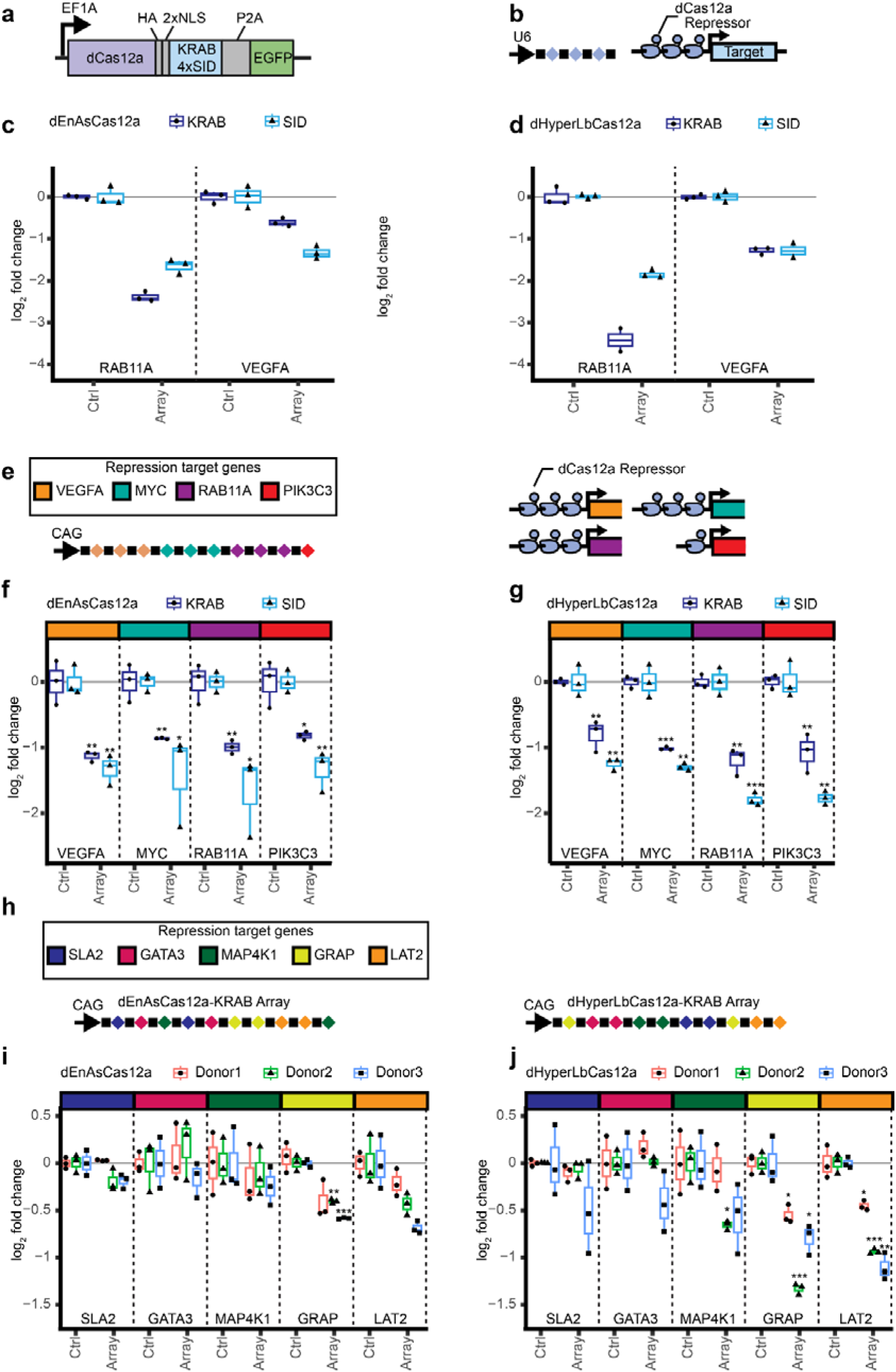
Multiplexed gene repression by dCas12a-KRAB and dCas12a-4xSID. **(a)** dCas12a repressor constructs **(b)** Experimental design for testing multiplexed repression of a single gene by dCas12a-KRAB and dCas12a-4xSID constructs. Repressors were co-transfected into HEK293T cells along with a pre-crRNA of three crRNAs targeting a gene promoter of interest or a control pre-crRNA containing three non-targeting crRNAs. Two days post-transfection, dCas12a-positive cells were sorted on EGFP expression, RNA was harvested, and gene expression changes were assessed via RT-qPCR**. (C)** Repression of the *RAB11A* (nTPM = 295.3) and *VEGFA* (nTPM = 9.7) promoters by dEnAsCas12a-KRAB and 4xSID fusions (n = 3 independent experiments per transfection condition) **(d)** Repression of the *RAB11A* and *VEGFA* promoters by dHyperLbCas12a KRAB and 4xSID fusions (at least two independent experiments per transfection condition) **(e)** Experimental design for testing highly multiplexed gene repression by dCas12a repressors. A pre-crRNA of ten crRNAs targeting four genes: *VEGFA* (nTPM = 9.7), *MYC* (nTPM = 154.0), *RAB11A* (nTPM = 295.3), and *PIK3C3* (nTPM = 14.7) was designed to test highly multiplexed repression. HEK293T cells were co-transfected with dCas12a repressors and either this array or a control pre-crRNA containing ten non-targeting crRNAs and relative gene expression was determined as in **(b). (f)** Repression of four target genes by dEnAsCas12a KRAB and 4xSID fusions using a pre-crRNA of length ten (n = 3 independent experiments per transfection condition) **(g)** Repression of four target genes by dHyperLbCas12a KRAB and 4xSID fusions using a a pre-crRNA of length ten (n = 3 independent experiments per transfection condition) * p < 0.05, ** p < 0.01, *** p < 0.001, two-sided FDR-adjusted student’s t-test **(h)** Experimental design for testing dCas12a-KRAB repressors in primary human T cells. A pre-crRNA of ten crRNAs targeting five genes: *SLA2* (Ctrl Ct = 25-26)*, GATA3* (Ctrl Ct = 24-25)*, MAP4K1* (Ctrl Ct = 25)*, GRAP* (Ctrl Ct = 29-30)*, and LAT2* (Ctrl Ct = 27-28) was designed to test highly multiplexed repression. T cells from three donors were co-transduced with dCas12a repressors and either this array or a control pre-crRNA containing ten non-targeting crRNAs. Cells were selected for dCas12a expression using Blasticidin, RNA was harvested 17 days post-transduction, and gene expression changes were assessed via RT-qPCR. **(i)** Repression of five target genes by dEnAsCas12a-KRAB using a pre-crRNA of length ten (n = 3 independent donors. Individual data points show qPCR technical replicates) **(j)** Repression of five target genes by dHyperLbCas12a-KRAB using a pre-crRNA of length ten (n = 3 independent donors. Individual data points show qPCR technical replicates). * p < 0.05, ** p < 0.01, *** p < 0.001, two-sided FDR-adjusted Student’s T-test

Next, we tested multiplexed repression by targeting several genes simultaneously. To do so, we co-transfected the dCas12a-based repressors with a pre-crRNA array targeting four different genes into HEK293T cells **(Fig. 2e)**. All repressors significantly repressed all four target genes (p < 0.05 for EnAs repressors, p < 0.005 for HyperLb repressors, FDR-adjusted Student’s T-test), including *PIK3C3* which is only targeted by a single crRNA. The dCas12a-4xSID fusions were as effective as dCas12a-KRAB fusions at repressing gene expression **(Fig. 2f-g)**. Taken together, these data demonstrate that dCas12a-SID repressors are highly effective and can serve as an alternative to dCas12a-KRAB.

### Lentiviral delivery of dHyperCas12a-KRAB enables strong gene repression

High-throughput, multiplexed screens require stably expressed dCas12a fusions, typically achieved via lentiviral delivery. A challenge is that previous studies have found that expression of dCas12a-KRAB fusions from low copy number integrated lentiviral vectors can result in low activity^10^. We hypothesized that dHyperLbCas12a effectors, which have the strongest activity under low crRNA expression (**Supp Fig. 1a**), can overcome that limitation.

To test that hypothesis, we created a HepG2 hepatocellular carcinoma cell line that stably expressed dHyperLbCas12a-KRAB. We then lentivirally transduced a construct expressing a pre-crRNA containing four crRNAs targeting the promoter of hexokinase domain containing 1 (HKDC1), a gene involved in maternal glucose homeostasis^58–60^. In doing so, we repressed *HKDC1* expression by 85%. That amount of repression was comparable to the ∼75% reduction in *HKDC1* transcript we observed in *HKDC1* knockout cell lines **(Supp. Fig 2c).** From those results, we conclude that lentiviral delivery of dHyperLbCas12a-KRAB is highly effective at repressing the expression of individual genes.

We further generalized that outcome to multiplexed repression in two additional cell types. First, We created A549 lung adenocarcinoma cell lines that stably expressed dHyperLbCas12a-KRAB or dHyperLbCas12a-4xSID. We then transduced the cell lines with a construct expressing a pre-crRNA with crRNAs targeting the promoter of *MYC*, *PIK3C* (crRNA spacer sequence originally reported in Ref 44), *RAB11A*, and *VEGFA*. In that system, dHyperLbCas12a-KRAB significantly repressed all four target genes, while dHyperLbCas12a-4xSID repressed two (p ≤ 0.05, FDR-adjusted Student’s T-test). For dHyperLbCas12a-KRAB, repression ranged from only ∼10% repression for *MYC* to 55-60% repression for *VEGFA* and *RAB11A*. Meanwhile, dHyperLbCas12a-4xSID was unable to repress *MYC* or *VEGFA*, but achieved ∼15% repression of *PIK3C3* and 35% repression of *RAB11A* **(Supp. Fig 2d)**. Because *MYC* and *PIK3C3* are both essential genes in A549 cells, it is possible that the weaker repression of those target genes resulted from effects on cell fitness^61^. In support of that possibility, we used flow-cytometry to sort A549 repressor cell lines transfected with the multiplexed pre-crRNA plasmid, and found that dCas12a repressor expression was several-fold lower in array-expressing cells than in a bulk transfected population (**Supp. Fig 2e)**.

Next, we co-transduced human T cells from three independent donors with dHyperLbCas12a-KRAB or dEnAsCas12a-KRAB and either a pre-crRNA targeting the promoters of *SLA2, GATA3, MAP4K1, GRAP, and LAT2* or a control pre-crRNA containing ten non-targeting crRNAs **(Fig. 2h)**. We observe similar trends for both Cas12a orthologues. For example, both dEnAsCas12a- and dHyperLbCas12a-KRAB showed the strongest repression of *GRAP* and *LAT2*, though dHyperLbCas12a showed stronger repression (mean log_2_ fold change across donors of −0.89 and −0.87 vs. −0.47 and −0.42) consistent with our previous findings **(Fig. 2i-j)**. While we do not observe effects for all target genes in this cell type, these results indicate the potential of using dCas12a for multiplexed repression in primary cells.

Taken together, these results demonstrate that dHyperLbCas12a repressors are suitable for use in multiplexed high-throughput screens. Also, as a step towards low multiplicity of infection screens, we demonstrate repression when pre-crRNA constructs are transduced and expressed from the genome at a copy number that is much lower than with transient transfection.

### dCas12a-P300 fusions robustly activate gene expression

Gene activation using dCas12a has most frequently been shown using fusions to the VPR activation domain^41,42,46,48^. While VPR is a potent activator of transcription, it has mostly been used to target gene promoters, not enhancer elements, and has been shown to be toxic in certain contexts^25,62–64^. Fusions of dCas9 to the histone acetyltransferase domain from P300 have been shown to be highly effective for targeting the non-coding genome, and a previous study showed that dCas9-P300 leads to stronger activation than dCas9-VPR when targeting a gene enhancer and promoter simultaneously^62,65^. dLbCas12a, but not dAsCas12a, has been shown to activate gene expression from both enhancers and promoters using single crRNAs^43^. We also tested activation using dAsCas12a-P300 and dLbCas12a-P300 fusions at two genomic loci using single crRNAs **(Supplementary Fig. 3a-b)**. We found that dLbCas12a-P300 activated both target genes, while dAsCas12a-P300 activated one of the genes, consistent with previous reports that LbCas12a is a stronger epigenome editor than AsCas12a, and our dCas12-VPR and KRAB comparisons **(Fig. 1a-b)**^41^. From this we conclude that the dLbCas12a-P300 fusions more strongly and more robustly activate gene expression than dAsCas12a-P300 fusions.

Above, we show that fusing KRAB and VPR to other variants of dCas12a can enhance repression or activation. Based on that result, we hypothesize that dEnAsCas12a-P300 and dHyperLbCas12a-P300 will also improve the efficacy of gene activation over fusions to wild type Cas12a proteins. Specifically, we created dEnAsCas12a-P300 and dHyperLbCas12a-P300 fusions in a plasmid containing a strong promoter and a NLS^48^ **(Fig. 3a)**. We then created pre-crRNA arrays of three crRNAs targeting each of the HBE1 promoter, the *MYOD1* promoter, or a *MYOD1* enhancer ∼5 kb upstream of the *MYOD1* promoter^65,66^. We co-transfected plasmids expressing the dCas12a activators with plasmids expressing the pre-crRNA arrays into HEK293T cells. **(Fig. 3b)**. All four fusion proteins modestly but significantly increased target gene expression (p < 0.005, FDR-adjusted Student’s T-test). dEnAsCas12a-P300 activation strength ranged from 1.2-to 18.2-fold, while dHyperLbCas12a activation strength ranged from 2.6- to 16.7-fold over non-targeting control. Contrary to our expectations, dCas12a-VPR led to stronger *MYOD1* activation from the distal enhancer than dCas12a-P300 for both orthologues **(Fig. 3c-d)**.

**Figure 3.**
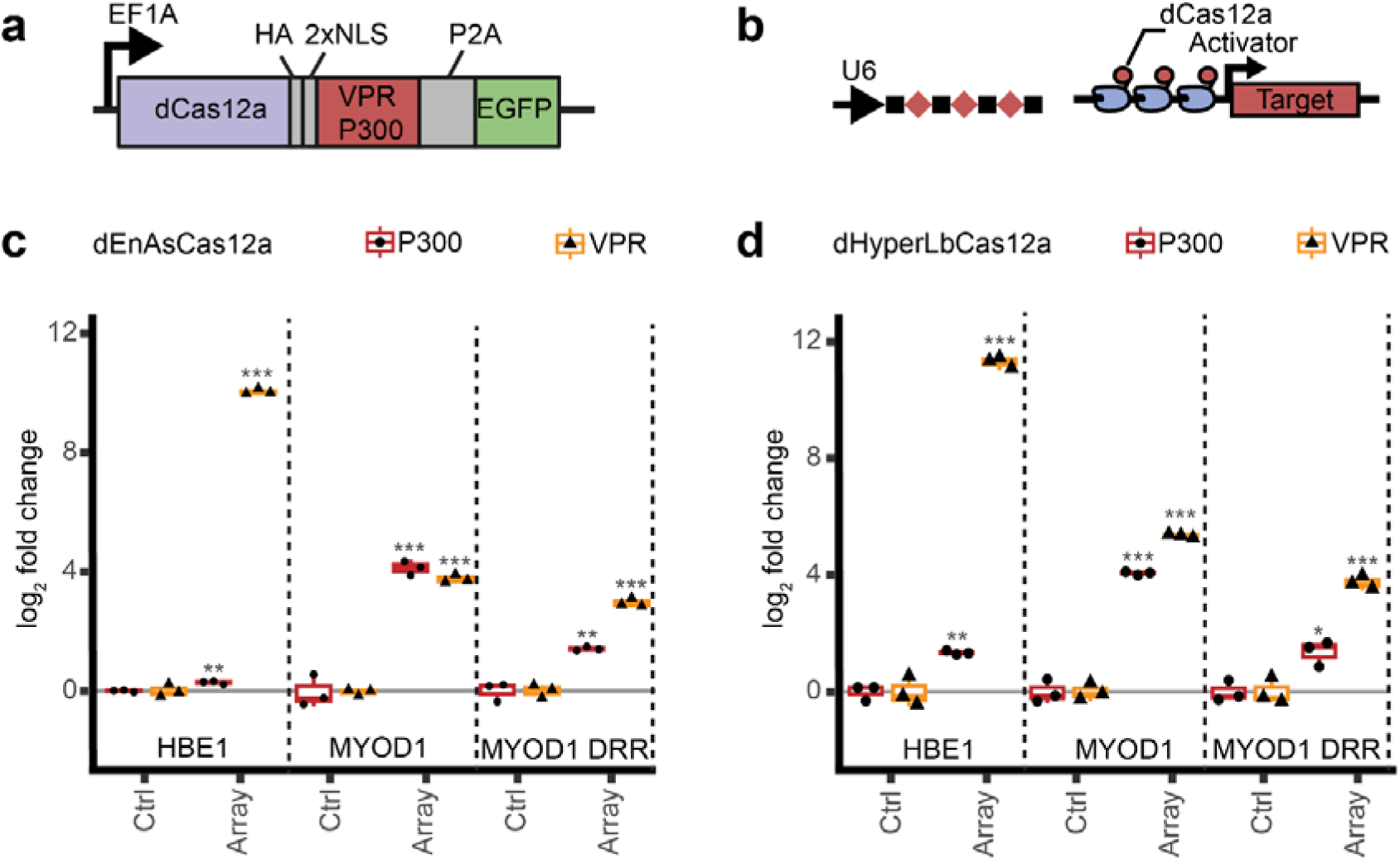
Activation of single genes using dCas12a. **(a)** dCas12a activator constructs **(b)** Experimental design for testing multiplexed repression by dCas12a-VPR and P300 constructs. Activator constructs were co-transfected into HEK293T cells along with a pre-crRNA of three crRNAs targeting a gene of interest or a control pre-crRNA of three non-targeting crRNAs. Two days post-transfection, dCas12a-expressing cells were sorted on EGFP, RNA was harvested, and gene expression changes were assessed via RT-qPCR**. (c,d)** Activation of the *HBE1* (nTPM = 0.1) promoter, *MYOD1* (nTPM = 0.0) promoter, or *MYOD1* distal regulatory region (DRR) by **(c)** dEnAsCas12a-VPR and P300 fusions (at least two independent experiments per transfection condition) or **(d),** dHyperLbCas12a-VPR and P300 fusions (n = 3 independent experiments per transfection condition)

We also evaluated whether all three crRNAs were needed, and whether there were additive or epistatic interactions between the crRNAs as reported previously^41,42,67,68^. We created pre-crRNA arrays that used three crRNAs to target either the *HBE1* or *MYOD1* promoter in all one-, two-, and three-way combinations of those crRNAs. We then co-transfected the pre-crRNA plasmids into HEK293T cells along with a dCas12a activator and assessed gene activation (**Supp Fig. 3c-f**). Generally, for dCas12a-VPR fusion proteins, multiple crRNAs increased activation strength. At both loci tested, the strongest pre-crRNA contained more than one crRNA, and combinations of three crRNAs were at least as strong as the best single crRNA. In contrast, for P300, we frequently observed reduced activation when we delivered multiple crRNAs, including examples where the combinations of crRNAs had weaker effects than the best single crRNA. Previous studies have also shown mixed effects from using multiple Cas9 gRNAs or Cas12a crRNAs, suggesting potential steric hindrance between the P300 fusions^43,67^.

### dCas12a-P300 fusions enable multiplexed gene activation in human cell lines

We next evaluated whether dCas12a activators are also capable of highly multiplexed gene activation. We first evaluated activity using transient delivery that typically achieves a high copy number of plasmids in each cell. To do so, we co-transfected dCas12a activators with a plasmid expressing a pre-crRNA of ten crRNAs from an RNA Pol II promoter into HEK293T cells. The crRNAs consisted of one crRNA targeting the *AR* promoter (originally reported in Ref 46), three crRNAs targeting the *HBE1* promoter, and six crRNAs targeting either the promoter or a distal regulatory region of *MYOD1* **(Fig. 4a)**. In all but one case, the dCas12a fusions significantly increased expression of all three target genes (p < 0.01, FDR-adjusted Student’s T-test, dHyperLbCas12a-VPR activation of *AR* was not significant) **(Fig. 4b-c)**. The range of activation varied among genes and between activators. For example, activation of *HBE1* was stronger with VPR fusions (up to ∼3200-fold) than with P300 fusions (up to 5-fold); whereas activation of *MYOD1* was consistently ∼16-fold for both P300 and VPR. Meanwhile, activation of AR was weaker overall (1.2-2.0-fold). That range of *AR* activation was the same order of magnitude as observed in previous studies^41^.

**Figure 4.**
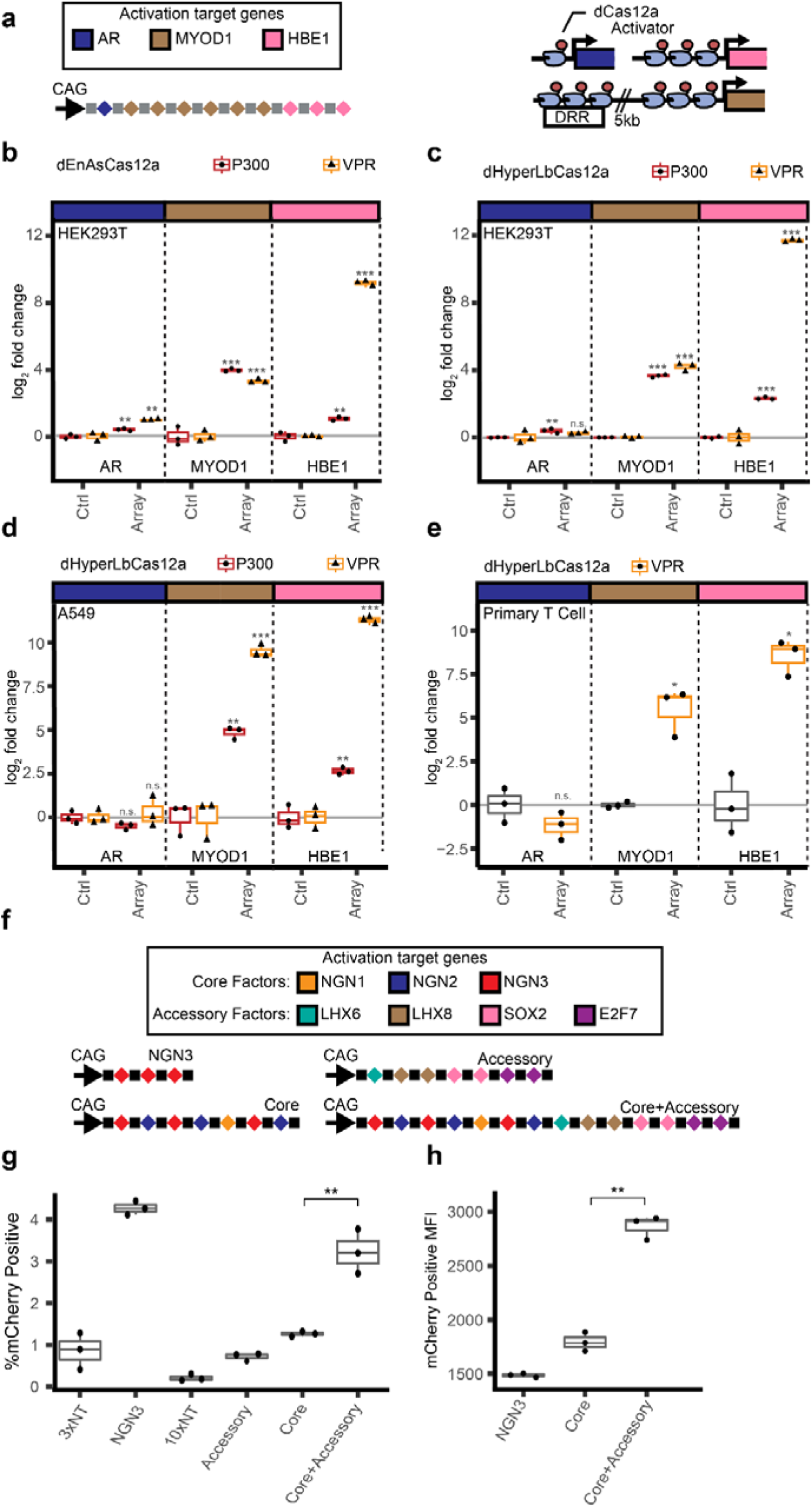
Activation of multiple genes using dCas12. **(a)** Experimental design for testing highly multiplexed gene activation by dCas12a activators. A pre-crRNA containing ten crRNAs targeting three genes: *AR*, *MYOD1*, and *HBE1*, was designed to test highly multiplexed activation. HEK293T cells were co-transfected with dCas12a activators and either this pre-crRNA or a control pre-crRNA of ten non-targeting crRNAs and gene expression was determined as in Fig. 3**. (b-e)** Activation of three target genes using the multiplexed pre-crRNA by **(b)** HEK293T cells transfected with dEnAsCas12a-VPR and P300 fusions (n = 3 independent experiments per transfection condition). HEK293T target expression levels: *AR* (nTPM = 2.2), *MYOD1* (nTPM = 0.0), and *HBE1* (nTPM = 0.1) **(c)** HEK293T cells transfected with dHyperLbCas12a-VPR and P300 fusions (n = 3 independent experiments per transfection condition), **(d)** A549 cells that stably express dHyperLbCas12a-VPR or dHyperLbCas12a-P300 (n = 3 independent experiments per transfection condition) A549 target expression levels: *AR* (nTPM = 4.0), *MYOD1* (nTPM = 0.0), and *HBE1* (nTPM = 0.4), and **(e)** primary T cells from three independent donors transduced with dHyperLbCas12a-VPR. RNA was harvested five days post-transduction. T Cell Expression: *AR* (Ctrl Ct = 35), *MYOD1* (Ctrl Ct = 40), and *HBE1* (Ctrl Ct = 33-36) **(f)** Experimental design for testing multiplexed activation of factors driving differentiation from iPSCs to neurons. pre-crRNA arrays were designed to target *NGN3*, a set of core neurogenic transcription factors (core), a set of accessory factors (accessory), or both (core+accessory). **(g,h)** An iPSC cell line with a *TUBB3A-*mCherry reporter that stably expresses dHyperLbCas12a-VPR was transduced with the pre-crRNA arrays in **(f)** or with control pre-crRNA arrays of non-targeting crRNAs. Cells were selected with puromycin, and after 4 days flow cytometry for mCherry was performed in order to determine the degree of *TUBB3A* expression in the cells as a proxy for neuronal differentiation. **(g)** Percent of mCherry positive cells and **(h)** MFI of mCherry positive cells were measured.

Many study designs require lentiviral delivery or integration of DNA expression cassettes. Those delivery approaches result in fewer effective molecules per cell, and thus require more potent effector proteins. We therefore evaluated whether the dCas12a system could drive robust, multiplexed gene activation when delivered lentivirally. We generated A549 lung adenocarcinoma cell lines that stably expressed dHyperLbCas12a-P300 or dHyperLbCas12a-VPR. We then transfected those cell lines with the same pre-crRNA targeting *AR, MYOD1, and HBE1* and found comparable effects on gene expression **(Fig. 4d)**. Both *MYOD1* and *HBE1* were significantly upregulated (p < 0.005, FDR-adjusted Student’s T-test), with effects ranging from 6-fold to ∼2500-fold; whereas there was no significant effect on *AR* expression.

To evaluate generality to other cell models, we also evaluated multiplexed activation in a HepG2 liver cancer cell line stably expressing dHyperLbCas12a-P300. In that model, *MYOD1* and *HBE1* were also significantly upregulated (p < 0.001, FDR-adjusted Student’s T-test) with effects ranging from ∼100-600-fold. In that HepG2 model, the effect on *MYOD1* was stronger than the effect on *HBE1* **(Supp. Fig 4a)**.

To generalize to additional gene loci, we also created two unique pre-crRNA arrays that each contain two crRNAs targeted to the promoter of *HKDC1*. We find that dHyperLbCas12a-P300 is able to significantly upregulate *HKDC1* expression using either array (p < 0.001, FDR-adjusted Student’s T-test) with similar effect size of ∼3-4 fold upregulation **(Supp. Fig 4b)**. This result underscores the robustness of stably expressed dHyperLbCas12a-P300 since, unlike *HBE1* and *MYOD1*, *HKDC1* has moderate basal expression in these cells (nTPM = 17.1), making it harder to detect increases in expression.

### dHyperLbCas12a-VPR fusions enable multiplexed gene activation in cultured primary human cells

Model systems for studying primary cell differentiation commonly involve controlling the expression of several genes at once^16,69^. As a key step towards such highly multiplexed control of gene regulation in primary cells, we focused on demonstrating dCas12a-based gene activation in T cells and in an human induced pluripotent stem cell (iPSC)-based model of neuronal differentiation.

To demonstrate multiplexed control of gene expression in T cells, we co-transduced dHyperLbCas12a-VPR along with the pre-crRNA targeting *AR*, *HBE1* and *MYOD1* into primary human CD3+ T cells from three different donors. The results were similar to those observed in cell lines, namely that there was significant activation (p < 0.05, One-Sided Student’s Paired T-Test) of *HBE1* and *MYOD1*, with effects ranging from ∼45-350-fold upregulation **(Fig.4e)**. Also as in the cell lines, there was little or no activation of AR gene expression.

To demonstrate multiplexed control of gene expression in iPSC cell differentiation, we used an iPSC cell line that had previously been modified to express mCherry cotranslationally with *TUBB3A*, a marker of neuronal differentiation^13^. We further modified the line to stably express dHyperLbCas12a-VPR. We designed a pre-crRNA array of three crRNAs to target the promoter of *NGN3*, a strong driver of neuronal differentiation, to act as a positive control. We also designed a pre-crRNA array containing seven crRNAs that target three core neurogenic transcription factors (deemed “Core”): *NGN1*, *NGN2*, and *NGN3*. We designed a pre-crRNA array containing seven crRNAs that target four accessory genes that have been shown to enhance neuronal differentiation when co-activated in tandem with a core factor (deemed “Accessory”): *LHX6*, *LHX8*, *SOX2*, and *E2F7*^13^. Lastly, we designed a pre-crRNA array containing all 14 crRNAs and targeting both core and accessory factors together (deemed “Core+Accessory”) **(Fig. 4f)**.

We transduced the dHyperLbCas12a-VPR cell line with these pre-crRNAs and then measured the percentage of cells that were mCherry positive when gating such that ∼1% of cells transduced with a pre-crRNA array containing non-targeting control crRNAs are mCherry positive **(Fig. 4g, Supp. Fig 5)**. We find that ∼4% of cells that received the *NGN3* targeting array are mCherry positive, while ∼1.25% of cells that received the “core” pre-crRNA array and ∼3.5% of cells that received the “Core+Accessory” pre-crRNA array are mCherry positive. As expected, activation of the accessory factors alone did not lead to an increase in mCherry positive cells over control **(Fig. 4h)**. While we did not observe higher rates of mCherry positive cells in the Core or Core+Accessory conditions vs. the *NGN3*-targeting array, we do observe an (∼3x) increase in mCherry positive cells between Core and Core+Accessory conditions, suggesting increased differentiation efficiency due to multiplexed activation. Furthermore, we observe increased (∼2x) MFI of the mCherry positive population in the Core+Accessory condition vs. both the *NGN3* and core conditions, suggesting higher *TUBB3A* expression in these cells **(Fig. 4i).**

Together, these results demonstrate that dCas12a activators are capable of multiplexed activation of gene expression in multiple primary cell systems in which dCas12a effectors have been integrated and remain stably expressed.

### Multiplexed activation and repression within a single cell using dCas12a effectors

The use of epigenome editors from two different Cas12a orthologues created an opportunity for simultaneous gene activation and repression within a single cell. Since Cas12a will process crRNAs from a transcript containing other elements, and each Cas12a orthologue specifically recognizes its own direct repeat sequence, we hypothesized that a hybrid array containing both As and Lb Cas12a crRNAs could enable programmed activation and repression from a single array. To test that possibility, we created a hybrid pre-crRNA array containing four AsCas12a crRNAs targeting the *HBE1* or *MYOD1* promoters and four LbCas12a crRNAs targeting the *RAB11A* or *MYC* promoters, separated by a non-targeting crRNA (**Fig. 5a)**. We validated the hybrid pre-crRNA array with single dCas12a effectors by co-transfecting with either dHyperLbCas12a-KRAB or dEnAsCas12-VPR into HEK293T cells. Doing so increased expression of *HBE1* and *MYOD1* with dEnAsCas12-VPR, and decreased expression of *MYC* and *RAB11A* with dHyperLbCas12a-KRAB **(Supp. Fig 6a-b)**.

**Figure 5.**
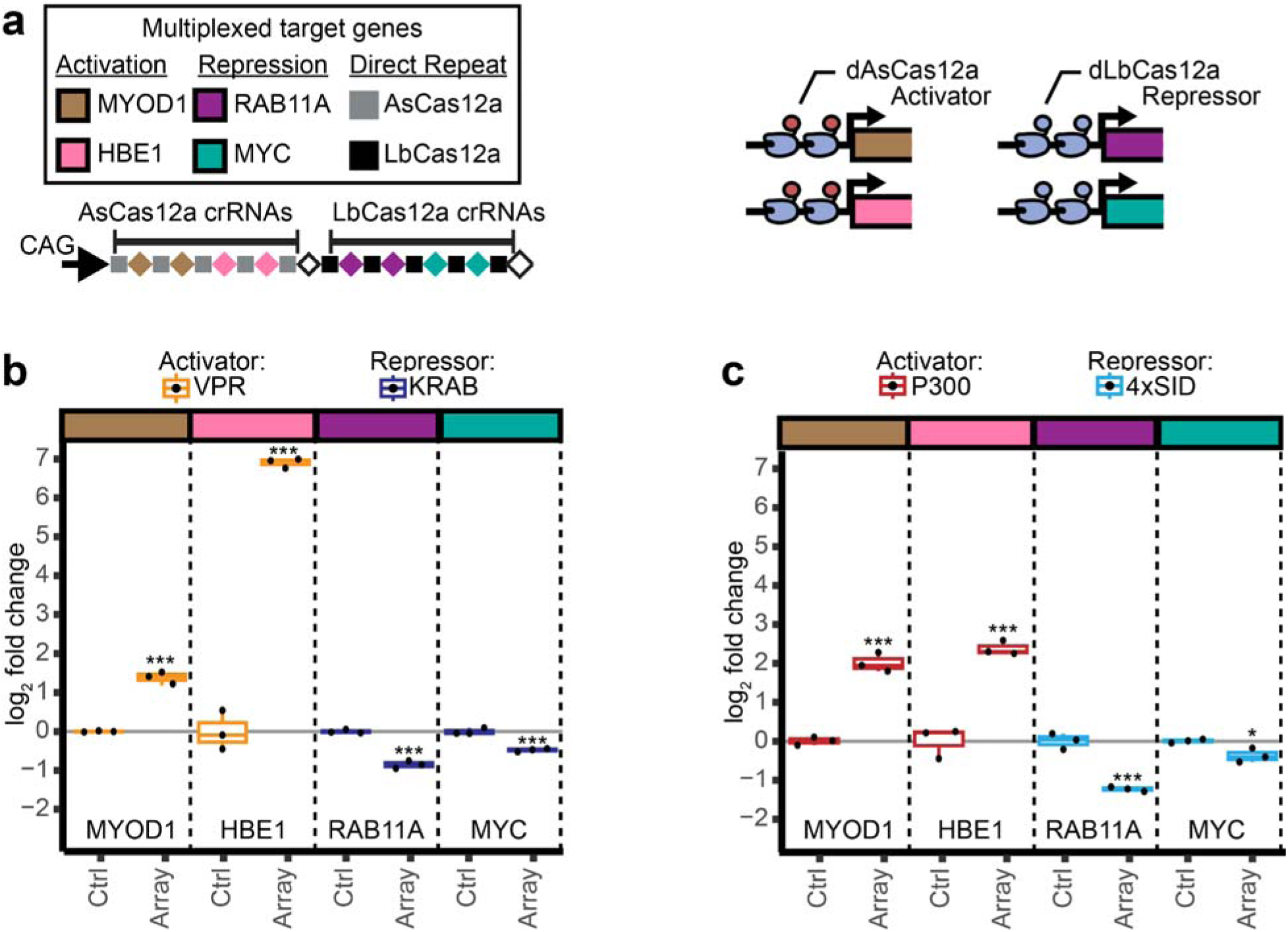
Programmed, simultaneous activation and repression within a single cell. (**a)** Experimental design for simultaneous activation and repression. A hybrid pre-crRNA composed of both Lb and As Cas12a crRNAs is co-transfected with a dEnAsCas12a activator and a dHyperLbCas12a repressor in order to achieve programmed activation and repression within a single cell. The hybrid pre-crRNA or a control pre-crRNA of ten non-targeting crRNAs were co-transfected into HEK293T cells with a dEnAsCas12a activator and a dHyperLbCas12a repressor. Cells were sorted and changes in target gene expression were assessed as in Fig 2**-4. (b,c)** Simultaneous activation and repression by **(b)** dEnAsCas12a-VPR and dHyperLbCas12a-KRAB or **(c)** dEnAsCas12a-P300 and dHyperLbCas12a-4xSID (n = 3 independent experiments per transfection condition).

We then evaluated simultaneous activation and repression by co-transfecting a dEnAsCas12 activator, a dHyperLbCas12a repressor, and hybrid pre-crRNA arrays into HEK293T cells. We evaluated two combinations of activators and repressors: dEnAsCas12a-VPR for activation with dHyperLbCas12a-KRAB for repression; and dEnAsCas12a-P300 for activation with dHyperLbCas12a-4xSID for repression. For both pairs of effectors, we observed significant and simultaneous activation and repression of all the target genes (**Fig. 5b-c**, p < 0.05, FDR-adjusted Student’s T-test). From this, we demonstrate that hybrid pre-crRNA arrays can simultaneously activate and repress gene expression within the same cell.

To generalize this result, we also designed and tested a similar pre-crRNA containing the same crRNA sequences, but with dHyperLbCas12a activators and dEnAsCas12a repressors. As before, we first tested the ability of single dCas12a effectors to use the pre-crRNA and found that dHyperLbCas12a-VPR and dEnAsCas12a-KRAB could significantly activate or repress the target genes when co-transfected with the hybrid crRNA **(Supp. Fig 6c-d)**. We next co-transfected a dHyperLbCas12a Cas12a activator and a dEnAsCas12a repressor along with the hybrid pre-crRNA into HEK293T cells. We found that dHyperLbCas12-VPR was able to significantly activate both target genes (p < 0.05, FDR-adjusted Student’s T-test), but dEnAsCas12a-KRAB was unable to repress either target gene **(Supp. Fig 6e)**. Meanwhile, the combination of dHyperLbCas12a-P300 and dEnAsCas12a-SID was able to significantly activate *HBE1* and *MYOD1* expression while significantly repressing *RAB11A* expression (p < 0.01, FDR-adjusted Student’s T-test), but not *MYC* expression (p = 0.172, FDR-adjusted Student’s T-test) **(Supp. Fig 6f)**.

Cas12a nucleases can exhibit low activity, as measured by indel formation, using the crRNAs of other Cas12a orthologues^70^. If this low-level orthologous crosstalk activity is also observed with CRISPRi and CRISPRa, then it could interfere with the hybrid pre-crRNA system. However, since both Cas12a orthologues are co-transfected in our system, we wanted to test not just whether each dCas12a can use the orthologous crRNA, but also whether it can use the orthologous crRNA when competing with the homologous dCas12a.

To evaluate crosstalk in such a setting, we co-transfected dHyperLbCas12a-VPR with the hybrid pre-crRNA containing AsCas12a crRNAs for activation as well as either dEnAsCas12a or Cas9 as a transfection control for number and size of plasmids. We found that, by itself, dHyperLbCas12a-VPR can use AsCas12a crRNAs but with less efficacy than homologous LbCas12a crRNAs. Specifically, when dHyperLbCas12a-VPR was cotransfected with Cas9, there was 150-fold activation of *HBE1* and no significant activation of *MYOD1* **(Supp. Fig 6g)**. The observed activation of *HBE1* was ∼30-fold less than when dHyperLbCas12a-VPR was co-transfected with its homologous hybrid pre-crRNA array **(Supp. Fig 6a)**.

However, that crosstalk is alleviated when dHyperLbCas12a and dEnAsCas12a are both present and competing for crRNAs. Specifically, when dHyperLbCas12a-VPR was co-transfected with dEnAsCas12a, we observed no significant activation of either gene, and relative *HBE1* expression dropped to a 1.25-fold change over non-targeting control **(Supp. Fig 6g)**.

That result was essentially similar when we tested dEnAsCas12a-VPR in the same way. Specifically, dEnAsCas12a-VPR was able to significantly activate *HBE1* by ∼6.5 fold using LbCas12a crRNAs when co-transfected with Cas9. However, there was no effect when dEnAsCas12a-VPR was co-transfected with dHyperLbCas12a **(Supp. Fig 6h)**. Together, these results confirm that while dEnAsCas12a and dHyperLbCas12a can use orthologous crRNAs for activation, they are not able to do so when the homologous Cas12a is also present and competing for crRNAs, such as in the hybrid pre-crRNA system developed here.

### Large-scale comparison of dCas12a repressors in a low MO fitness screen

To enable large-scale comparison of the dCas12a repressors described above and to assess their suitability for use in screening applications, we performed a screen of genes that are predicted to impact cell proliferation. We created a library containing 171 pre-crRNAs targeting the promoters of 145 core essential genes^71^, 48 pre-crRNAs targeting the promoters of 20 cell surface control genes^45^, 50 non-targeting pre-crRNAs, and 25 intergenic control crRNAs. Each pre-crRNA contained four crRNAs, and we required that all gene targets have at least four crRNAs with CRISPick on-target efficiency score of > 30%. We created an As and Lb Cas12a version of this library using the same spacer sequences but As- or Lb-specific direct repeats.

Next, we transduced the appropriate library at an MOI of 0.2 into A549 cells stably expressing either dHyperLbCas12a-KRAB, dHyperLbCas12a-4xSID, dEnAsCas12a-KRAB, or dEnAsCas12a-4xSID, with three replicates per cell line. Cells were selected for crRNA expression using puromycin, and cell pellets were collected to assess relative pre-crRNA abundance at 5, 7, 14, and 21 days post-transduction (**Fig. 6a**).

**Figure 6.**
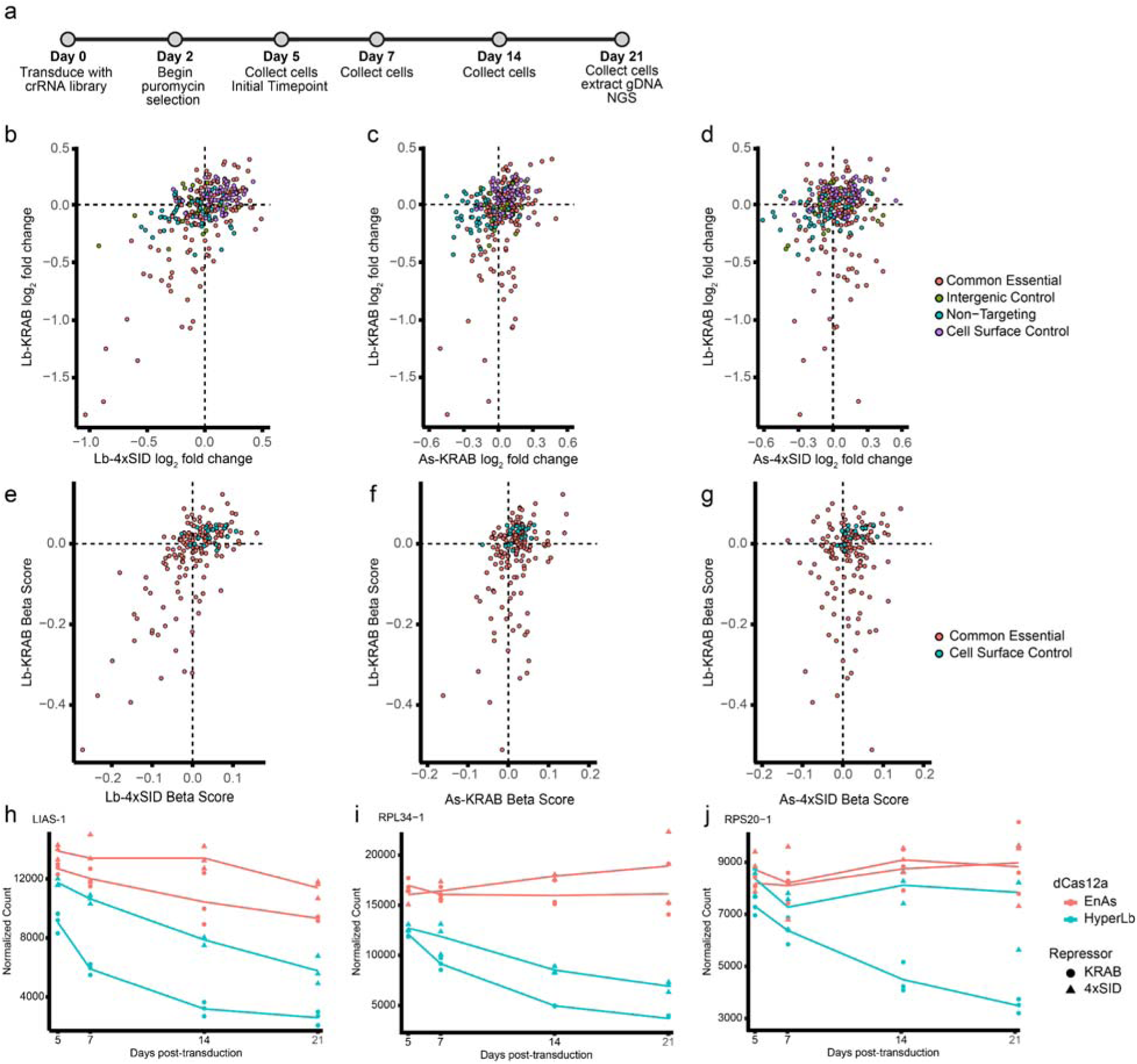
Fitness Screen Results. **(a)** Timeline of the fitness screen. A549 cells stably expressing dHyperLbCas12a-KRAB, dHyperLbCas12a-4xSID, dEnAsCas12a-KRAB, or dEnAsCas12a-4xSID were transduced in triplicate with a pre-crRNA array library targeting the promoters of essential genes (n = 171), cell surface controls (n =48), intergenic controls (n=25), and non-targeting controls (n = 50). They were then selected with puromycin, and cell pellets were collected at 5-, 7-, 14-, and 21-days post-transduction to track pre-crRNA abundance over time. **(b-d)** log_2_ fold-change of pre-crRNA arrays from day 21 vs. day 5 for **(b)** dHyperLbCas12a-KRAB vs. dHyperLbCas12a-4xSID, **(c)** dHyperLbCas12a-KRAB vs. dEnAsCas12a-KRAB, **(d)** dHyperLbCas12a-KRAB vs. dEnAsCas12a-4xSID **(e-g)** Beta scores from the MaGeck MLE module for change in crRNA abundance across the time course for **(e)** dHyperLbCas12a-KRAB vs. dHyperLbCas12a-4xSID, **(f)** dHyperLbCas12a-KRAB vs. dEnAsCas12a-KRAB, **(g)** dHyperLbCas12a-KRAB vs. dEnAsCas12a-4xSID **(h-j)** Examples of normalized read counts over time for crRNAs that significantly repressed their target genes **(h)** in both dHyperLbCas12a repressor cell lines and the dEnAsCas12a-KRAB cell lines, **(i) in** both dHyperLbCas12a repressor cell lines, and **(j)** in the dHyperLbCas112a-KRAB cell line only

There was strong concordance in read counts between replicates for all Cas12a repressor screens. Specifically, correlation between replicates at day 21 was R > 0.91 for the KRAB repressor screens and R > 0.79 for the SID repressor screens **(Supp. Fig. 7-8)**. The strongest and most consistent repressive effects occurred in the dHyperLbCas12a-KRAB screen. That screen had much higher correlation between replicates for log_2_ fold changes of pre-crRNAs targeting essential genes (R > 0.8) compared to all the other repressors (−0.081 < R < 0.276). That screen also had greater effects compared to the other repressors **(Fig. 6b-e, Supp. Fig. 9).** Representative examples for each system show genes that were significantly repressed by both dHyperLbCas12a repressors and dEnAsCas12-KRAB; by both dHyperLbCas12a repressors but not the dEnAsCas12a repressors; and by dHyperLbCas12a-KRAB alone **(Fig. 6h-j)**.

Overall, we conclude that KRAB is a stronger repressor than 4x SID for targeting gene promoters, and that dHyperLbCas12a-KRAB is suitable for use in low MOI screens, while dEnAsCas12a is not. We based that conclusion on comparing effects across the four screens. Using a linear model of change in log_2_ fold change over time and a threshold of FDR < 0.001, we detected 28 significantly depleted pre-crRNAs targeting essential genes in the dHyperLbCas12a-KRAB condition, five in the dHyperLbCas12a-4xSID condition, two in the dEnAsCas12a-KRAB condition, and zero in the dEnAsCas12a-4xSID condition using DeSeq2. We obtained similar results at the gene level when performing a time course analysis with the MAGeck MLE module and wald FDR < 0.05-31 genes were significantly depleted in the dHyperLbCas12a-KRAB condition, five in the dHyperLbCas12a-4xSID condition, one in the dEnAsCas12a-KRAB condition, and zero in the dEnAsCas12a-4xSID condition **(Supp. Table 2)**. Comparing these hits to DepMap gene essentiality data for A549 cells, we find that the dHyperLbCas12a-KRAB hits are strongly enriched in highly essential genes (gene effect ≤ −1)^72^ **(Supp. Fig 10)**. Furthermore, we note that the effects of dHyperLbCas12a-4xSID were stronger than those of either As repressor and the most closely correlated with dHyperLbCas12a-KRAB (R = 0.68 vs. R = 0.38 for dEnAsCas12a-KRAB and R = 0.13 for dEnAsCas12a-4xSID), even if they did not achieve significance.

### High-throughput multiplexed epigenome editing screens via dHyperLbCas12a-KRAB

The above findings create the foundation for high-throughput combinatorial epigenome editing screens with the dCas12a platform. To demonstrate that possibility, we performed a combinatorial screen of two enhancers that previously were hypothesized to control activation of Period 1 (PER1) gene expression in response to glucocorticoids. Glucocorticoids are steroid hormones that have a major role in peripheral circadian rhythms throughout the body^73^. *PER1* is a key gene in that regulation. In A549 cells, *PER1* stands out because its expression is uniquely sensitive to very low concentrations of glucocorticoids^8^. The mechanism by which such low concentrations of glucocorticoids activate *PER1* expression remains unresolved. Based on ChIP-seq and reporter assays studies, there are two promoter-proximal glucocorticoid-responsive regulatory elements, denoted here as Enhancer A and Enhancer B, that may cause glucocorticoid-dependent *PER1* activation. Furthermore, Enhancer A can control reporter gene expression at very low glucocorticoid concentrations, suggesting it may be responsible for the *PER1* gene expression response at those low concentrations^8^ **(Fig. 7a)**.

**Figure 7.**
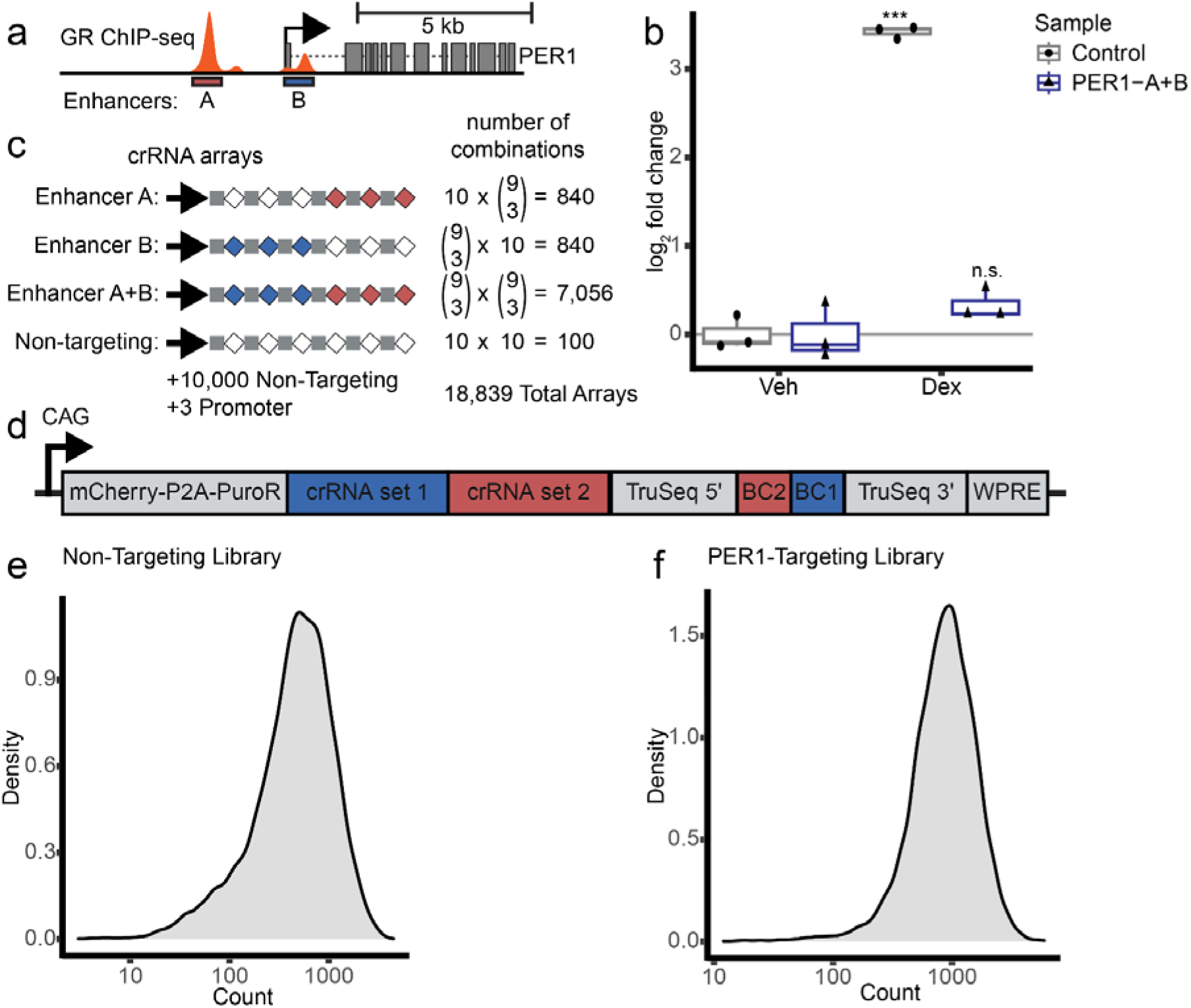
Construction of a crRNA library to target two *PER1* enhancers. **(a)** two enhancers, labeled A and B, control the glucocorticoid-responsiveness of the gene *PER1*. These sites are bound by the glucocorticoid receptor after treatment with the synthetic glucocorticoid dexamethasone **(b)** dHyperLbCas12a-KRAB is capable of dampening the *PER1* glucocorticoid response through dual targeting of the enhancers in (**a)**. A549 cells stably expressing dHyperLbCas12a-KRAB and either a pre-crRNA of four crRNAs targeting the *PER1* enhancers (2 crRNAs per enhancer) or a control pre-crRNA of four non-targeting crRNAs were created. These cell lines were treated with either 100nM dexamethasone or vehicle for three hours before RNA was harvested and changes in *PER1* expression were determined using RT-qPCR. **(c)** Schematic showing makeup of the 8,836 member *PER1*-targeting pre-crRNA library. Pre-crRNAs of six crRNAs were created to target each of the *PER1* enhancers either individually or in combination **(d)** schematic showing the completed library array constructs. Two sets of 3 crRNAs were cloned into a CAG expression vector to create dual-targeting pre-crRNAs of 6 crRNAs with a bipartite barcode flanked by TruSeq adapter sequences. The crRNAs are also located on the same transcript as the mCherry-P2A-puroR selection marker. **(e,f)** Density plots showing the distribution of crRNA counts for each library. The distribution of pre-crRNA barcode counts for **(e)** the 10,003 member non-targeting library. **(f)** the 8,836 member *PER1*-targeting library

We first definitively established that Enhancer A and Enhancer B together control the PER1 glucocorticoid response. To do so, we transduced A549 cells stably expressing dHyperLbCas12a-KRAB with either a pre-crRNA array targeting both Enhancer A and Enhancer B, or a non-targeting pre-crRNA array. We then selected cells for pre-crRNA expression, and treated them for three hours with either 100 nM dexamethasone (Dex) or ethanol as a vehicle control. With non-targeting crRNAs, *PER1* expression increased nearly 11-fold with dex treatment; while with crRNAs targeting Enhancer A and Enhancer B, *PER1* expression increased by only ∼1.3-fold in the dex condition (p = 8.8 × 10^−6^, Student’s T-Test) **(Fig. 7b)**. Together, these results indicate that Enhancer A and Enhancer B together account for all or nearly all of the *PER1* glucocorticoid response.

Enhancer A and Enhancer B were also sensitive to specific chromatin modifications. The *PER1* response was not significantly altered when we repeated the experiments in A549 cells expressing dHyperLbCas12a-4xSID instead of dHyperLbCas12a-KRAB (p = 0.45, Student’s T-Test). That finding suggests the *PER1* glucocorticoid response is differentially sensitive to KAP1-mediated vs Sin3a-mediated repression; and that the dCas12a repressors described here can identify such differential sensitivities. Additionally, activating the enhancers with VPR or P300 in the absence of glucocorticoids did not significantly increase *PER1* expression (p = 0.48, p =0.20, Student’s T-Test). **(Supp. Fig. 11)**, indicating that glucocorticoid receptor binding and co-factor recruitment may be necessary for *PER1* activation via those enhancers.

Next, to dissect the independent and combinatorial contributions of Enhancer A and Enhancer B, we designed a library of pre-crRNA arrays to target dCas12a-KRAB to the enhancers alone or in combination **(Fig. 7c)**. Overall, our design was to deliver pre-crRNAs containing six crRNAs, where the first three crRNAs target Enhancer B, and the second three target Enhancer A. To target the enhancers together, we used on-target crRNAs in all six positions. To target Enhancer A or Enhancer B individually, we used non-targeting crRNAs in the first three or second three positions, respectively. In total, we designed nine on-target crRNAs per enhancer, and assembled DNA sequences for all 84 unique triples thereof **(Fig. 7c, Supp. Fig. 12a)**. We also designed ten sets of three non-targeting crRNAs to target enhancers individually.

To construct the screening library, we used a nested cloning strategy to simultaneously assemble the pre-crRNA array and a series of 8 bp barcodes that encode the contents of the array **(Supp. Fig 12)**^74,75^. In the first round of cloning, we inserted the crRNAs targeting Enhancer B or non-targeting crRNAs (**Supp. Fig 12b**). In the second round of cloning, we inserted the crRNAs targeting Enhancer A or non-targeting crRNAs **(Supp. Fig 12c)**. That resulted in a library of 8,836 unique pre-crRNA arrays composed of 7,056 pre-crRNA arrays targeting both enhancers, 1,680 pre-crRNA arrays targeting the enhancers individually, and 100 pre-crRNA arrays targeting neither enhancer **(Fig. 7c-d, Supp. Fig 12d)**. In parallel, we separately constructed a control library of 10,000 non-targeting six-crRNA arrays and three pre-crRNAs targeting the *PER1* promoter **(Supp. Fig 13)**. Since most (80%) of the *PER1*-targeting library consists of pre-crRNAs targeting both enhancers, this large control library will enable better separation between cells with high and low *PER1* expression. When constructing the targeting and control libraries, we also enabled sorting crRNA-expressing cells by co-transcribing a selectable marker with the pre-crRNA^44^ **(Fig. 7d, Supp. Fig 12)** and high-throughput sequencing of the barcode by including TruSeq adapter sequences flanking the embedded barcode **(Supp. Fig 12)**. We confirmed the library construction via sequencing on an Illumina Nextseq instrument. We recovered all on-target pre-crRNA arrays, and 9,890 of the 10,000 negative control pre-crRNA arrays. The missing negative control pre-crRNA arrays were explained by missing or low-abundant crRNAs during DNA synthesis. Aside from those dropouts, the pre-crRNA arrays were evenly represented in the assay library **(Fig. 7 e-f)**.

To assay the pre-crRNA libraries, we lentivirally transduced them into A549 cells stably expressing dHyperLbCas12a-KRAB **(Fig. 8a)**. Ten days post-transduction, we treated crRNA-expressing cells with either low (200 pM) or high (100 nM) concentrations of the synthetic glucocorticoid dexamethasone, or with ethanol to control for solvent effects. After three hours, we collected cells with the highest and lowest 12% *PER1* expression^76^ **(Supp. Fig. 14a, Supp. Fig 15)**. Finally, we estimated the abundance of each array in the sorted cell populations via high-throughput sequencing of the barcodes. We also confirmed that pre-crRNA arrays remained intact and correctly associated to the corresponding barcode throughout the screen, consistent with findings from other Cas12a studies^77^ (**Supp. Fig 16**, Supplementary text).

**Figure 8.**
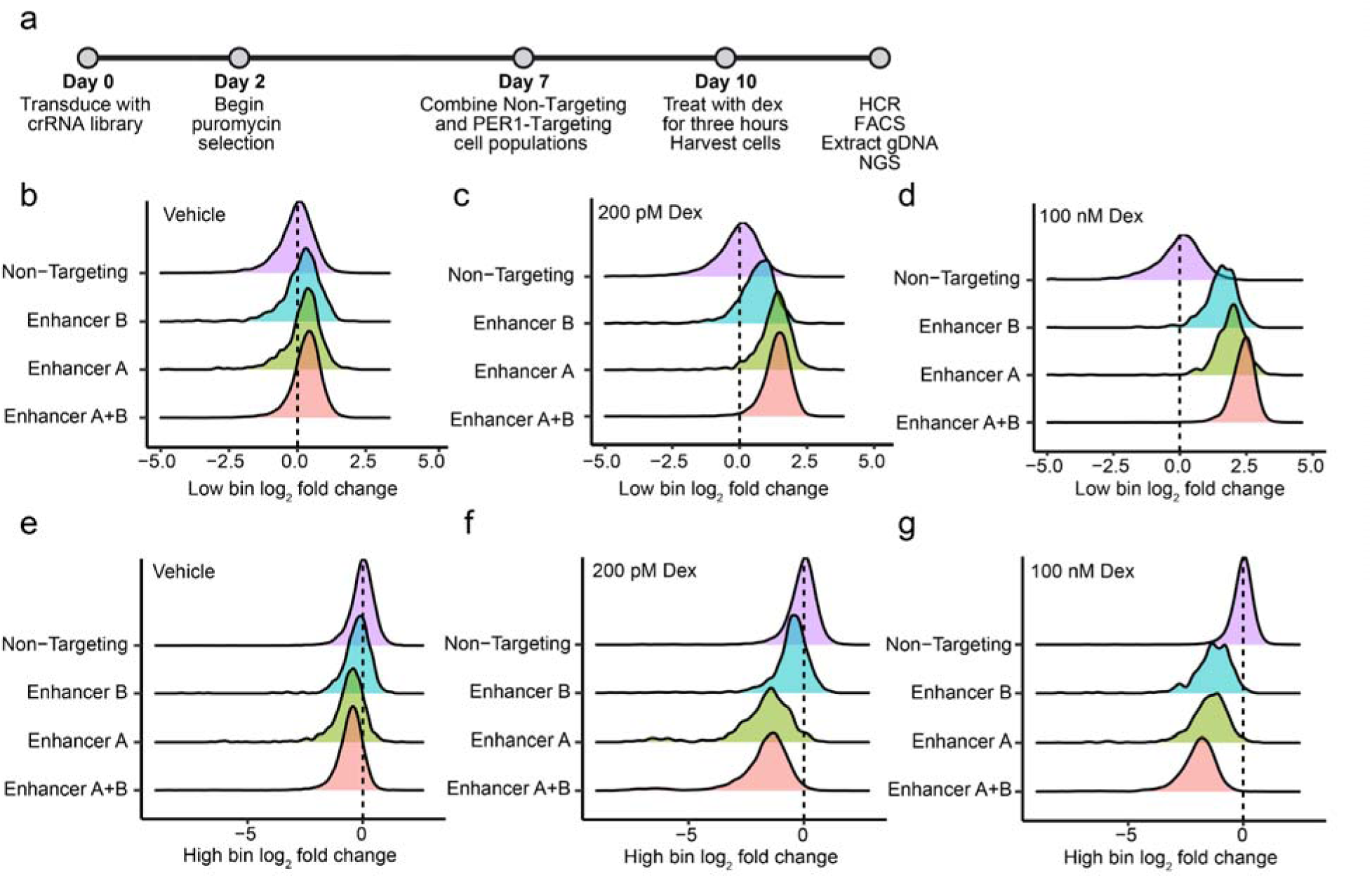
*PER1* Screen Results. **(a)** Timeline of the *PER1* screen. A549 cells stably expressing dHyperLbCas12a-KRAB were transduced with either the library of *PER1*-targeting arrays (n = 8,836) or the library of non-targeting arrays and promoter-targeting arrays (n=10,003). They were then selected with puromycin, after which the cell populations for the two different libraries were combined. Cells were treated with either an ethanol control (vehicle), 0.2nM dexamethasone (low Dex), or 100nM dexamethasone (high Dex) for three hours before being harvested for HCR. Cells were then sorted into high and low bins (top and bottom 12%) based on *PER1* expression. gDNA was recovered and sequenced and array abundance in high and low bins wa compared to input. **(b,c,d)** Enrichment in pre-crRNA barcode abundance in low bin vs. input for **(b)** vehicle **(c)** 0.2nM Dex and **(d)** 100nM Dex. **(e,f,g)** Enrichment in pre-crRNA barcode abundance in high bin vs. input for **(e)** vehicle **(f)** 0.2nM Dex and **(g)** 100nM Dex.

Overall, we found that Enhancer A controls the *PER1* response to low glucocorticoid concentrations, and that both Enhancers A and B together control the *PER1* response at high glucocorticoid concentrations. Specifically, at 200 pM dexamethasone, pre-crRNA arrays targeting Enhancer A were ∼2.6-fold enriched in low-*PER1*-expressing cells, and depleted by 2.9-fold in high-*PER1*-expressing cells. In contrast, pre-crRNA arrays targeting only Enhancer B were 1.7-fold enriched in low-*PER1*-expressing cells and depleted by 1.4-fold in high-PER1-expressing cells **(Fig. 8c)**. At 100 nM dexamethasone, pre-crRNA arrays that target both Enhancer A and Enhancer B were ∼5.4-fold enriched in low-*PER1*-expressing cells and 3.7-fold depleted in high-*PER1*-expressing cells; and pre-crRNA arrays targeting only Enhancer A or only Enhancer B had intermediate effects (3.9-fold and 3.0-fold enriched, and 2.7- and 2.4-fold depleted, respectively) **(Fig. 8d)**. Finally, in the ethanol treatment condition, the on-target pre-crRNA arrays were only weakly depleted from high-*PER1*-expressing cells (up to 1.4-fold) **(Fig. 8b)**. Together, these results reveal glucocorticoid-dose-dependent enhancer activity controlling *PER1* expression.

We did not find strong evidence for substantial heterogeneity in the effects of the individual crRNAs. Specifically, we tested if, when targeting enhancer A or B individually, any of the crRNAs consistently increased or decreased the effect of the pre-crRNA array **(Supp. Table 3)**. In the 100 nM Dex high bin, 2 of the 9 crRNAs targeting enhancer A had significant effects and 6 of the 9 crRNAs targeting enhancer B had a significant effect (FDR < 0.05). Of those eight crRNAs with significant effects, four increased crRNA array activity, and four decreased that activity. However, the magnitude of those effects were small, with changes in mean pre-crRNA enrichment of between 10% and 36% when those crRNAs were present **(Supp. Fig 17a-c)**. The individual crRNA effects also did not correlate with GC content or the on-target activity score predicted by CRISPick **(Supp. Table 3)**.

## Discussion

We systematically evaluated and improved technologies for dCas12-based epigenome editing. First, we consistently evaluated a wide range of dCas12a-based epigenome effectors delivered by both transient plasmid transfection and sustained lentiviral transduction. Through that effort, we identified a number of more effective epigenome effectors, and demonstrated that the effectors work when lentivirally integrated into the genome at lower copy number, including in cultured primary human cells. Second, we demonstrated that expressing pre-crRNAs using RNA polymerase II allows for longer and more complex multiplexing than using RNA polymerase III. Finally, combining those advances, we present what is to the best of our knowledge the first successful dCas12a CRISPRi screens, and the first dCas12a screen on changes in gene expression. Those screens involve simultaneous delivery of six crRNAs, demonstrating the potential for highly multiplexed CRISPRi screens on human gene expression.

A key challenge in epigenome editing screens is that effectors and crRNAs typically must be delivered at low copy number. Previous dCas12a-based CRISPRi studies have required delivery of high levels of dCas12a and pre-crRNA into HEK293T cells^42,44^; and others have found similar results for dCas12a-based gene activation^25^. Those limitations can be overcome by using a nicking-biased version of AsCas12a to improve repression. While effective, doing so also introduces insertions of deletions in up to ∼7% of target sites^10^. Alternatively, co-expressed nanobodies have been used to activate promoters genome-wide; but doing so increases complexity by adding components to the screening system^25^. By instead basing our screen on dHyperLbCas12a, which we and others have found to be more effective under low copy number conditions^48^, we were able to screen directly using a dCas12a-based epigenome effector. Further, we expect other dHyperLbCas12a epigenome editing fusions can be used in similarly designed screens. Finally, by reading out on gene expression directly using HCR^76,78^ rather than on cell proliferation as in previous Cas12a-based epigenome screening studies, we discover new mechanisms controlling the expression of individual genes.

A key enabling feature for our screens is the creation of barcoded pre-crRNA libraries using a nested cloning strategy^74,75^. Up to four crRNAs and a barcode sequence can be added to the pre-crRNA during each round of cloning, allowing for the generation of large combinatorial libraries from small and easy to synthesize oligonucleotide pools. For example, the 10,000 member non-targeting library in this study was generated from 200 oligonucleotides that were 235 nt each. The end result is a library of uniquely barcoded pre-crRNAs containing all possible combinations of crRNA sets added during each round of cloning. The barcode allows for screening with pre-crRNAs that exceed the length of high-throughput short-read sequencing methods. We believe that, compared to dCas9-based multiplexed epigenome editing strategies, this type of combinatorial screening allows for much more highly-multiplexed experiments, can enable more efficient gene activation or repression via multiple crRNAs targeting the same promoter, and is cheaper to synthesize due to shorter crRNA length. Additionally, while high MOI screens coupled with single cell sequencing technologies can allow for delivery of many dCas9 gRNAs, these experiments limit library size and are much more expensive and technically complex than the screening strategy presented here. Together, those advances make the screening method flexible and applicable to a wide range of highly-multiplexed combinatorial studies of gene expression.

We also develop and validate a number of dCas12a-based technologies that can be used for future highly-multiplexed epigenome editing studies. We introduce a suite of newly-validated dCas12a epigenome editors that expand the available options for multiplexed targeting of the non-coding genome. We selected two epigenome-modifying domains, P300 and SID, that had been used successfully to modulate gene expression from both promoters and enhancers when fused to dCas9 or other DNA-binding proteins^4,43,57,65^. We successfully expand these results by creating dCas12a-P300 and dCas12a-4xSID fusions for both As and LbdCas12a variants, and demonstrate highly-multiplexed repression and activation with pre-crRNAs of up to ten crRNAs, including when dCas12a effectors are stably expressed in A549 cells.

Many aspects of health and disease involve both increases and decreases in gene expression. Modeling those systems via simultaneous activation of some genes and repression of others has been previously demonstrated using dCas9 orthologs and a dual sgRNA expression vector^13,18,19^. Here, we demonstrate an alternative approach using a hybrid dCas12a pre-crRNA to simultaneously activate two genes and repress two genes. That strategy can be expanded in several ways. For example, it is likely, given the compatibility with dCas12a-VPR, P300, KRAB, and 4xSID, that the system could be expanded to other dCas12a epigenetic fusions or nuclease active Cas12a. It is also likely that other CRISPR proteins that process their own pre-crRNAs, such as Cas13a, could be added^79–81^. The approach could also be used for high-throughput multiplexed screens that combine activation and repression. Together, those advances have the potential to greatly improve modeling of biological systems that involve activation and repression.

The results of this study also point to areas for future mechanistic investigation. We and others have found that using multiple gRNAs or crRNAs to target a genomic element generally often leads to stronger activation or repression for dCas9 or dCas12a fusions to KRAB and VPR, but that for P300 adding additional gRNAs or crRNAs can lead to weaker activation, even in cases where the same combination of crRNAs leads to stronger activation for dCas12a-VPR fusions^41–43,67,68^. Therefore, there may be rules for effective multiplexed activation with P300 that have not yet been discovered. Additionally, whether all dHyperLbCas12a epigenome editors are suitable for high-throughput screening and the upper limit for pre-crRNA length in a high-throughput screening setting have yet to be determined. A final challenge will be understanding the profile of off-target effects and how they scale with the number of crRNAs contained in each array. We expect that a full understanding of those off-target events would require genome editing to stably record^82,83^.

Finally, though the experimental technologies now exist for highly multiplexed epigenome editing screens, the potential scale of such studies will demand corresponding statistical methods for experimental design and interpretation. A key observation from our screen of the PER1 locus is that identifying epistatic interactions in gene regulation requires a great degree of replication, and we expect that the outcomes of our study can be useful for power analyses to design future such screens. Though our screen focused on two enhancers controlling glucocorticoid dependent *PER1* expression, we demonstrated high activity of pre-crRNAs containing six crRNAs that could be used to target combinations of six regulatory elements. The scale of combinatorial libraries that can be generated with six crRNAs can already exceed practical limitations in the number of cells or sequencing required. For example, there are 10^16^ unique combinations of six transcription factors, far exceeding the number of cells that could be reasonably cultured; and a screen of one million three-way combinations of transcription factors would only cover less than 0.2% of the possible combinations. Meanwhile, the pre-crRNA libraries could readily be expanded to include more crRNAs due to the high processivity of Pol II. For those reasons, highly multiplexed combinatorial screens will necessarily rely on and generate very sparse observations relative to the number of possible combinations, and statistical methods to prioritize informative combinations and interpret effects at that sparsity will be needed to realize the full potential of highly multiplexed epigenome editing screens.

## Acknowledgements

This work was supported by NHGRI RM1 HG011123 to TER, NHGRI UM1 HG012053 and NIMH R01 MH125236 to TER and CAG, NCI R01 CA289574 to CAG, Yosemite Fund, and the Duke-Coulter Translational Partnership. We thank Joshua B. Black for guidance on neuronal differentiation experiments. We also acknowledge the use of biorender.com to create some of the figures in this manuscript.

## Contributions

T.E.R. conceived the study, and T.E.R. and C.A.G. funded, and supervised the study. T.E.R. and S.M.M. designed and planned experiments. M.C.H. cloned LbCas12a pre-crRNA expression plasmids used in **Figs.2c-d, 3c-d, and Supp. Fig.3c-f**, and completed the dLbCas12a activation experiments in **Supp. Fig. 3c-f**. C.M.A. and S.M.M. designed and performed T cell experiments in Figs. 2 and 4. S.M.M. completed all other experiments and analyzed resulting data. T.E.R. and S.M.M. wrote the manuscript, with input from C.A.G., C.M.A., and M.C.H.

## Methods

### Plasmid generation

Unless otherwise noted, cloning was performed using PCR of desired insert fragments with Q5 (New England Biolabs (NEB)), followed by Gibson Assembly using NEBuilder Hifi 2x Master Mix (NEB) to add inserts into the desired digested plasmid backbone. Cloning reactions were transformed into chemically competent NEB Stable E. Coli (NEB), purified using the Zyppy Plasmid Miniprep Kit (Zymo), and validated using Sanger sequencing (Azenta). All Cas12a ORFs and epigenome editing domain sequences can be found in **Supplemental Information.** All key plasmids from this study will be posted on Addgene.

First-generation dCas12a effector fusions were cloned into the FUGW plasmid backbone. FUGW-dLbCas12a was a gift from Dr. Charles Gersbach, and dLbCas12a sequence was originally derived from pY109 (Addgene 847740). dAsCas12a and dEnAsCas12a were derived from Addgene plasmids (114078 and 107943, respectively)^46^. Mutations to make dAsCas12a Ultra were introduced into the dAsCas12a plasmid. A gene fragment (Twist) with the dHyperLbCas12a ORF was synthesized and cloned into the same FUGW backbone. Effector fusions were generated using Gibson assembly or by standard subcloning methods. The 4xSID sequence was derived from Addgene 106399^4^ and the 3xKRAB sequence was taken from Addgene 128132^44^. Versions of each dCas12a effector were made with either a T2A-EGFP or T2A-puroR using subcloning. These constructs were used in **Fig. 1a-b, Supp. Fig 1, Supp. Fig 2a-b, and Supp. Fig 3a-f**.

The FUGW vector was then modified using Gibson assembly and subcloning to create a second generation of dCas12a effector fusions. The hUbc promoter was replaced with the EF1A promoter. The nucleoplasmin NLS on the C-terminus of dEnAsCas12a was exchanged for two copies of the c-Myc NLS. Lastly, the T2A-puroR was replaced by T2A-BlasticidinR. These dCas12a effectors were used in **Fig. 1c-d**, **Fig. 2-8, Supp. Fig 2c-e, Supp. Fig 4, and Supp. Fig 6-10.**

For expression of Cas12a pre-crRNA arrays using U6, pRDA_052 (Addgene 136474)^45^ was used for AsCas12a and an FUGW plasmid was used for LbCas12a. These constructs were used in **Fig. 1a-b, Fig. 2c-d, Fig. 3c-d, Supp. Fig 1, Supp. Fig 2a-b, and Supp. Fig 3a-f**. The FUGW LbCas12a crRNA expression vector was modified for U6 expression of either As or LbCas12a crRNAs by inserting an Esp3I cloning site flanked by two direct repeat sequences just downstream of the U6 promoter. These plasmids also included an hUbc promoter to express mCherry-P2A-PuroR for selection. These plasmids were used in **Fig. 1c-d**. A pHAGE plasmid was modified using Gibson assembly for CAG expression of either As or LbCas12a crRNAs. First, the promoter was changed from EF1A to CAG. Next, an mCherry-P2A-puroR ORF, followed by the MALAT1 triplex sequence used in a previous study^44^, followed by two direct repeat sequences flanking an Esp3I cloning site were inserted into the vector downstream of the CAG promoter. Finally, a kanamycin resistance cassette was inserted between the direct repeat sequences in order to limit recombination. This plasmid was used in **Fig. 1c-d**, **Fig. 2f-j**, **Fig. 4-7, Supp. 2c-e, Supp. 3g, Supp. Fig. 4b.** The sequence of this transcript can be found in **Supplemental Information**.

Cloning of crRNAs into expression vectors was performed using one-pot golden gate cloning **(Fig. 1a-b, Fig.2c-d, Fig. 3c-d**, **Fig. 7, Supp. Fig 1, Supp. Fig 2a-c, Supp. Fig 3a-f, Supp. Fig. 4, and Supp. Fig 10a)**. All crRNA spacer sequences used in this study can be found in **Supplemental Table 1**. Briefly, pairs of oligos were annealed, then phosphorylated using T4 Polynucleotide Kinase (NEB) in a 50 μL reaction. Next, 40 fmol of plasmid backbone and 2 μL of each annealed oligo pair were added to a one-pot golden gate reaction with Esp3I and T4 DNA Ligase (NEB) as has been done in previous studies^79,84^. Long pre-crRNA arrays in **Fig. 1c-d, Fig. 2f-j**, **Fig. 4-5, Supp. 2d-e, Supp. Fig 4b, and Supp. Fig. 6** were constructed using PCR followed by golden gate cloning. Briefly, the crRNA array was created by using multiple rounds of low cycle number PCR. Primers added one crRNA to each end of the growing array with each round of PCR. The last round of PCR also added Esp3I sites for Golden Gate cloning. This amplicon was then cloned into the plasmid backbone using one pot Golden Gate cloning with 40 fmol of plasmid and 120 fmol of PCR product. Long pre-crRNA arrays in **Fig. 2i-j and Fig. 4g-h** were synthesized as Golden Gate-compatible gene fragments (Twist) with Esp3I sites and 5’ and 3’ primer sequences for PCR amplification. Gene fragments were first amplified using low cycle number PCR. PCR products were purified using DNA Clean and Concentrator (Zymo), and then the amplicon was then cloned into the plasmid backbone using one pot Golden Gate cloning with 40 fmol of plasmid and 120 fmol of PCR product.

### CRISPRi Library Generation

First, CRISPick was used to design a library targeting the promoters of 291 previously identified core essential gene, 25 cell surface controls, and a handful of A549-specific essential genes using the CRISPRi window (TSS-50 to TSS+300) and the TTTV PAM^45,71^. We also used CRISPick to generate 200 non-targeting crRNAs and 100 intergenic control crRNAs. We identified 145 core essential genes and 20 cell surface control genes that had at least four crRNAs with on-target activity score ≥ 0.3. We also designed crRNAs to target thirteen synthetic lethal gene pairs that had at least two crRNAs with on-target activity score ≥ 0.3 either individually or in combination. Using these spacers, we designed a library of pre-cRNA arrays containing four crRNAs each. The final library was composed of 171 pre-crRNA arrays targeting the promoters of 145 core essential genes, 48 pre-crRNA arrays targeting the promoters of 20 cell surface control genes, 50 non-targeting pre-crRNA arrays, 25 intergenic control crRNA arrays, and 600 pre-crRNA arrays targeting the synthetic lethal genes either individually or in combination for a total of 893 pre-crRNA arrays.

The pre-crRNA arrays were ordered as two separate oligo pools (Twist), one for dHyperLbCas12a and one for dEnAsCas12a. These contained the same spacer sequences, but different direct repeat sequences. Each oligo pool was amplified via 8 low cycle number PCR reactions (Q5, NEB). PCR products were purified using the DNA Clean and Concentrator Kit (Zymo). These DNA fragments were then input into 8 ligation one-pot Golden gate reactions using 360 fmol of PCR product and 40 fmol of the CAG crRNA expression vector described above. These products were pooled and concentrated using the DNA Clean and Concentrator kit before being electroporated into Endura chemically competent cells (Lucigen). The 893-member library was purified using the Midi Plus kit (Qiagen).

### PER1 and Non-Targeting crRNA library generation

First, the CAG crRNA expression vector described above was modified to have the 3’ TruSeq adapter sequence downstream of the Esp3I cloning site instead of a flanking direct repeat sequence. This was done with Gibson Assembly. This sequence can be found in **Supplemental Information**. Spacer sequences used in the *PER1* and non-targeting libraries and oligos ordered for cloning these libraries can be found in **Supplemental Table 3**.

Next, nine unique crRNAs were designed to target each *PER1* enhancer using CRISPick. From those nine crRNAs 84 (9 choose 3) unique sets of three crRNAs were created to target each enhancer. Using those sets of crRNAs, a pool of 188 oligos, composed of two different groups was designed (Twist). Group 1 was composed of oligos containing the 84 sets of three enhancer B-targeting crRNAs plus 10 sets of three non-targeting crRNAs, as well as a nested cloning site and a unique 8nt barcode sequence **(Supp. Fig 12a)**. Group 2 was composed of oligos containing the 84 sets of three enhancer A-targeting crRNAs plus 10 sets of three non-targeting crRNAs, as well as the 5’ TruSeq adapter sequence for NGS and a unique 8nt barcode sequence. Barcode sequences were designed with a Hamming Distance of 3. Group 1 and group 2 oligos were amplified from the pool using separate PCR primer pairs **(Supp. Table 1)**. Then, Golden Gate cloning was used to insert the group 1 amplicons into the plasmid backbone. The plasmid was digested with Esp3I (NEB), while the PCR amplicon was digested with BsaI (NEB). These were purified using the DNA Clean and Concentrator Kit (Zymo). These DNA fragments were then input into 8 ligation reactions using 120fmol of PCR product and 40 fmol of digested plasmid. Ligated products were pooled and concentrated using the DNA Clean and Concentrator kit before being electroporated into Endura chemically competent cells (Lucigen). The 94-member library was purified using the Midi Plus kit (Qiagen). Group 2 amplicons were then cloned into the 94-member library using 8x one pot Golden Gate cloning with Esp3I as described in plasmid generation methods. This library was electroporated and purified as before, resulting in a final library with six crRNAs per array and 8,836 total crRNA arrays.

The non-targeting library was created by designing 50 non-targeting control crRNAs as well as three promoter-targeting crRNAs using CRISPick. To create a 10,000 member library 200 sets of 3 crRNAs were randomly chosen from the 50 non-targeting crRNAs and broken up into 100 sets for group 1 and 100 sets for group 2. Three sets of three promoter-targeting crRNAs were also included with group 1. The non-targeting library was then cloned the same way as the PER1 enhancer-targeting library. The three promoter-targeting crRNAs did not contain a nested cloning site and were therefore unchanged during the second round of cloning.

### Cell Culture

HEK293T, A549, and HepG2 cell lines were obtained from ATCC. Cell lines were authenticated using STR profiling through the Duke Cell Culture and DNA Analysis facility. Mycoplasma testing was performed by Eurofins. All cell lines were grown in humidified incubators at 37 °C and 5% CO_2_. Cells were cultured using the following media:

HEK293T: DMEM (Gibco) + 10% FBS + 1% penicillin-streptomycin
A549: F12K (Gibco) + 10% FBS + 1% penicillin-streptomycin
HepG2: EMEM (ATCC) + 10% FBS + 1% penicillin-streptomycin

The RVR-iPSC TUBB3A-mCherry reporter cell line has been characterized in previously published work^13^. iPSCs were maintained on Matrigel (Corning) coated plates in mTESR supplemented with mTESR plus 5x Supplement (Stem Cell Technologies). For neuronal differentiation, media was changed to DMEM/F-12 Nutrient Mix (Gibco, 11320), 1x B-27 serum-free supplement (Gibco), 1x N-2 supplement (Gibco), and 25 μg/mL gentamicin (Gibco) 24 hours post-transduction with pre-crRNA arrays.

Primary T cells were obtained by isolating CD3+ T cells from a leukopak (StemCell Technologies) using the EasySep^TM^ Human T cell isolation kit (StemCell Technologies). They were activated with anti-CD3/CD28 Dynabeads (Gibco) at a 3:1 bead to cell ratio and maintained in PRIME-XV media (Fujifilm) supplemented with 100 U/mL of IL-2, 5% human platelet lysate (Compass Biomedical), 100 U ml^−1^ penicillin, and 100 μg ml^−1^ streptomycin.

### Transfection

For experiments performed in 24-well plates **(Figs 1a, 2-5, Supp. 3a-b, Supp. 2e)**, cells were co-transfected with 400 ng of dCas12a effector plasmid and 250 ng array plasmid 24 hours after seeding using Lipofectamine 3000. For the experiments in **Figure 5 and Supp. Fig 6**, cells were co-transfected with 250 ng of each dCas12a effector plasmid, and 250 ng of hybrid crRNA, or 125ng of each control array (As and Lb Cas12a crRNAs). 48 hours post-transfection, cells were either harvested for flow cytometry **(Figs 1a, 2-5, Supp. Fig 6, Supp. 2e)**, or RNA was extracted directly from bulk transfected cells **(Supp. Fig 3a-b)**

For experiments performed in 96-well plates **(Figs. 1b, 1d-e, 4d, 7b, Supp. 1, Supp. 2a-d, Supp. 3c-f, Supp. 4c)** cells were reverse transfected with 80 ng of dCas12a effector plasmid and 50 ng array plasmid using Lipofectamine 3000. For the low crRNA condition in **Supp. Fig 1A**, cells were transfected with 5 ng of array plasmid. RNA was harvested from cells 48 hours post-transfection.

### RT-qPCR

For all experiments except those specifically listed below, RNA extraction and cDNA synthesis were completed using the two step Cells to Ct Kit (Thermo). 2 μL of cDNA was used in each PCR reaction. Multiplexed qPCR was performed using TaqMan Gene Expression Master Mix (Thermo), a FAM-labeled Taqman assay (Thermo) for the appropriate target gene, and a VIC-labeled, primer-limited endogenous control assay (Thermo) for ACTB. qPCR was performed on a StepOne Plus thermal cycler under the following conditions:

95 °C for 10 min
40x

95 °C for 15 seconds
60 °C for 1 minute

In instances where gene targets failed to amplify due to low/no expression, a Ct value of 40 was used.

For experiments in **Fig. 2i-j**, **Fig 4e, Supp. Fig 3a-b**, qPCR was performed as above except that RNA was extracted using the RNeasy mini kit (Qiagen). cDNA was then created using the High Capacity cDNA Reverse Transcription Kit (Applied Biosciences) or Super Script III Reverse Transcription Kit (Thermo) using 500 ng −1 μg of RNA. 1 μL of cDNA was used for qPCR.

For experiments to test dCas12a effector expression **(Supp. Fig 2e)**, the Cells to Ct kit was used as above and 2 μL of cDNA was used in the qPCR reaction. qPCR was performed using SYBR Green (Thermo) and primers to detect the Cas12a transcript or primers to detect ACTB. Primer sequences can be found in **Supp. Table 1**. Thermal cycling conditions were as follows:

95 °C for 10 min
40x

95 °C for 15 seconds
60 °C for 1 minute

All nTPM values indicated for qPCR target genes are from the Human Protein Atlas (proteinatlas.org)

### Stable cell line generation

In order to create A549 and HepG2 cell lines that stably express a dCas12a effector, lentivirus was created by the Duke Viral Vectors Core using standard reagents (psPAX2 and MD2.G). A549 and HepG2 cells were transduced with lentivirus at 1x concentration in media containing 10 μg/mL polybrene (Thermo). Media was changed 24 hours post-transduction. Cells were selected with 2.5 μg/mL Blasticidin-S-HCl. Once selection and expansion was complete, cell lines were validated by transduction or transfection of array constructs.

In order to create the stable crRNA array-expressing cell lines in **Fig. 7b**, **Supp. Fig 2c-d, Supp. Fig 3g, and Supp. Fig 10** and the Cas9 knockout cell lines in **Supp. Fig 2c**, we created lentivirus using standard reagents. Briefly, 90% confluent HEK293T cells in a 6 well plate were transfected with 600 ng psPax2, 450 ng of the lentiviral vector of interest, and 150 ng of MD2.G using Lipofectamine 3000. Media was changed 24 hours post-transfection, and supernatant was harvested 72 hours post-transfection. A549 and HepG2 cells were transduced with lentivirus at 1x concentration in media containing 10 μg/mL polybrene Media was changed 24 hours post-transduction. HepG2 cells were selected with 0.75 μg/mL puromycin, while A549 cells were selected with 1.0 μg/mL puromycin. Once selection and expansion was complete, cell lines were validated by RT-qPCR.

In order to create iPSCs and T cells that stably express a dCas12a effector and pre-crRNA array, dCas12a and pre-crRNA array lentivirus were created using a previously reported protocol for high titer lentivirus production^85^.

T Cells were transduced 24 hours post-activation in a 24 well plate at a concentration of 1.0×10^6^ cells/mL using 10% v/v of virus in cell culture media. 24 hours post-transduction, cells were split evenly into five wells of a 96 well plate and transduced with pre-crRNA array lentivirus at a concentration of 10% v/v. Cells were expanded and selected using 4 μg/mL Blasticidin-S-HCl. Once expansion and selection was complete, RNA was harvested and qPCR was performed to assess levels of target gene activation or repression.

iPSCs were transduced with dHyperLbCas12a-VPR in single cell suspension in mTESR (Stem Cell Technologies) supplemented with Rock inhibitor (Y-27632, Stem Cell Technologies) at a concentration of 5×10^5^ cells/mL. After 24 hours, media was changed to mTESR without Rock inhibitor. 30 hours after transduction, 5 μg/mL Blasticidin-S-HCl was added directly to the media. Cells were selected and expanded to 10 cm dishes. Cells were single cells passaged for the first passage after transduction to facilitate complete selection. For neuronal differentiation experiments, 60,000 cells were seeded in single cell suspension to each well of a matrigel-coated 24 well plate in mTESR supplemented with Rock inhibitor and 1 μL of pre-crRNA virus was added to each well (three independent transductions per pre-crRNA array construct). The next day, media was changed to neurogenic media (see above). At 30 hours post-transduction, puromycin was added directly to the media for a final concentration of 1 μg/mL. Daily media changes using neurogenic media plus 1 μg/mL puromycin were performed, and cells were harvested and analyzed via flow cytometry four days post-transduction.

### Flow Cytometry

For studies involving transfected cells, live cells were harvested from a 24 well plate 48 hours post-transfection and resuspended in 200 μL MACS Buffer (DPBS + 0.5% BSA + 2 mM EDTA). Cells were then sorted on a SONY SH800 sorter using a 100 μm chip. Populations were gated on FSC/SSC for single cells, and then gated for GFP positive cells, a marker of Cas12a expression **(Supp. Fig 18)**. Between 15,000 and 50,000 cells were sorted per sample, after which we proceeded with RNA extraction using the Cells to Ct kit.

For the iPSC studies, live cells were washed with PBS and harvested from a matrigel-coated 24 well plate four days post-transduction using Accutase (Stem Cell Technologies). Cells were resuspended in 150 μL MACS Buffer and analyzed on the Attune NxT Flow Cytometer (Thermo). Populations were gated on FSC/SSC to determine single cells, and then gated on mCherry using a threshold of ∼1% mCherry positive cells in a population of cells transduced with a non-targeting control pre-crRNA array **(Supp. Fig 5)**

To determine lentiviral titers for the CRISPRi libraries, A549 cells stably expressing dHyperLbCas12a-KRAB were transduced with serial dilutions of the lentiviral pre-crRNA libraries (either As or LbCas12a version). Media was changed 24 hours post-transduction, and cells here harvested 48 hours post-transduction, resuspended in 150 μL MACS Buffer, and analyzed on the Attune NxT Flow Cytometer (Thermo). Single cells were gated on using FSC/SSC, and then mCherry positive population was determined by gating on untransduced cells **(Supp. Fig 19)**. mCherry fluorescence was used as a marker of transduction and used for MOI calculations.

For the dHyperLbCas12a-KRAB screen, flow cytometry was performed by the flow cytometry core at the Duke Cancer Institute using an MoFlo Astrios EQ High Speed Sorter (Beckman Coulter). Cells were first gated on single cells using FSC/SSC. Next cells were gated on compensated mCherry, a marker of crRNA expression, and compensated FITC-488, a marker of PER1 transcripts by HCR. The top and bottom 12% of FITC-expressing cells were then sorted **(Supp. Fig 15)**.

### PER1 CRISPRi Screen

First, lentivirus was created for the PER1-Targeting and Non-Targeting crRNA libraries. For each library, two T-75s containing 90% confluent HEK293T cells were transfected with 4 μg of psPAX2, 3 μg of array plasmid, and 2 μg of MD2.G using 27 μL of PEI (1 mg/mL). Media was changed 24 hours post-transfection, and supernatant was harvested 72 hours post-transfection. For each library, 24 mL of supernatant was concentrated 10x using Lenti-X concentrator (Takara Bio) as directed.

Next, for each library 1×10^8^ cells in 500 cm^2^ dishes were transduced at low MOI using the concentrated lentivirus in media containing 10 μg/mL polybrene to achieve 1,000x library coverage. After 48 hours, cells were passaged 1:4 into media containing 0.75 μg/mL puromycin. Cells were passaged two more times (5 more days) to ensure that selection was complete. In order to ensure good library coverage, at each passage, cells from all plates per library were pooled and the number of cells passaged was at least 2,000x library size. After 3 passages in puromycin (7 days post-transduction), cells from each of the two libraries were pooled in equal numbers and co-cultured. Ten days post-transduction, cells were treated for 3 hours with 0.1% ethanol (vehicle), 200 pM dexamethasone, or 100 nM dexamethasone, with 2×10^8^ cells per condition. 1.4×10^7^ cells were taken from each condition, mixed, and pelleted in order to create a bulk sample to use as a comparison for crRNA enrichment. Additionally, 5×10^6^ A549 dHyperLbCas12a-KRAB cells (untransduced) were harvested for use as flow cytometry controls.

Cells were harvested and HCR Flow-FISH was performed^76^. PER1 transcripts were detected using a custom set of 20 probe pairs containing the B3 amplifier. Probe was used at 2x concentration compared to the standard protocol. Amplification was performed for 22 hours using the B3-488 amplifier. Flow cytometry controls were created as follows: negative control cells were taken from A549 cells that expressed dCas12a-KRAB, but were not transduced with a crRNA library. They were hybridized with probe, but set aside after the probe wash step. mCherry single color controls were taken from cells transduced with the PER1 and Non-targeting libraries. They were hybridized with probe, but set aside after the probe wash step. FITC-488 single color controls were taken from A549 cells that expressed dCas12a-KRAB, but were not transduced with a crRNA library. These cells went through the whole HCR protocol and had labeled PER1 transcripts. All cells were resuspended in MACS buffer and sorted as described above.

### CRISPRi Fitness Screen

Lentivirus was created for the As and LbCas12a pre-crRNA array libraries using the same method as the *PER1* screen. Next, since the pre-crRNA construct contains an mCherry marker, titer was determined via flow cytometry as described above (under flow cytometry).

Next, for each A549 cell line expressing a dCas12a effector, 5×10^6^ cells were seeded to a T-75 flask and transduced with the appropriate pre-crRNA array library at an MOI of 0.2. This was done in triplicate for each effector. Media was changed 24 hours post-transduction. 48 hours post-transduction, cells were passaged 1:4 into four new T-75s containing F12K media supplemented with 0.75 μg/mL puromycin. After selection (5 days post-transduction), an initial cell pellet was collected from each sample (2×10^6^ cells/pellet). Cells were selected with puromycin for one week, and then selected for one passage with 2.5 μg/mL Blasticidin-S-HCl to ensure that all cells continued to express the dCa12a repressor construct. Cell pellets were also collected at 7-, 14-, and 21-days post-transduction (1-2×10^6^ cells/pellet) to compare pre-crRNA abundance at those times to the initial timepoint.

### NGS Library Prep and Sequencing

For the *PER1* screen, genomic DNA was extracted from bulk-transduced cells that did not undergo HCR using the Puregene Cell Kit (Qiagen). Meanwhile, genomic DNA from cells sorted into high and low *PER1* expression bins was extracted using the FFPE miniprep kit (Zymo). Genomic DNA purified using the FFPE miniprep kit was then cleaned using Axiprep Magnetic Beads (Axygen) at a ratio of 0.9x in order to remove any small DNA fragments. Genomic DNA was then analyzed on a TapeStation 4200 in order to confirm that DNA was high quality and average fragment size was large (> 20 kb).

Next, crRNAs were amplified from genomic DNA (primer sequences can be found in **(Supp. Table 1)** using Q5 2x Master Mix (NEB) in 100 μL reactions with the following thermal cycling parameters:

98 °C for 45 seconds

27x

98 °C for 15 seconds
60 °C for 20 seconds
72 °C for 1 minute and 15 seconds

72°C for 5 minutes

We performed 18 PCR reactions for each sorted cell sample and 96 PCR reactions for the bulk unsorted sample. Axiprep magnetic beads were used at a 0.9x ratio to clean and size select 200 μL of PCR product in order to remove any adapter dimer. PCR products were quantified using a qPCR standard curve (KAPA ABI Prism qPCR Master Mix). Preparation of libraries from plasmid DNA used the same PCR conditions except with 12 cycles and 25 PCR reactions per plasmid library. In both cases, the number of cycles of PCR was determined empirically by first performing a qPCR using Q5 2x Master Mix and 1x EvaGreen (Thermo), and choosing the number of cycles that gave about one half of the maximum signal. Libraries were run using a 600 cycle paired-end kit NextSeq 2000 at a concentration of 950 pM. Custom primers were used for sequencing (Read 1, Index 1, Index 2, and Read 2, **Supp. Table 1**)

For the fitness screen, genomic DNA was extracted from cell pellets using the Puregene Cell Kit (Qiagen). Next, pre-crRNA arrays were amplified from genomic DNA using KAPA 2x PCR Master Mix (Roche) for 28-29 cycles. The plasmid libraries were also amplified using 13 cycles of PCR. We performed 8x 50 μL PCR reactions per sample. PCR cycle numbers were determined empirically by performing a qPCR using KAPA 2x PCR Master Mix with EvaGreen, similarly to the *PER1* library. Axiprep magnetic beads were used at a 0.9x ratio to clean and size select 200 μL of PCR product in order to remove any adapter dimer. PCR products were quantified using the Quant-It (Invitrogen). Libraries were run using a 300 cycle paired-end kit on the NextSeq 2000 at a concentration of 750 pM and with a 20% Phi X spike in. Standard Illumina primers were used for sequencing.

### Sequencing Data Processing

For the PER1 screen, FastQ files were generated using BCLConvert in BaseSpace. To extract barcode sequences, reads containing the barcode were first trimmed using Cutadapt, aligned to a custom reference of barcode sequences using Bowtie2, and counts for each barcode were generated using Samtools with the following parameters:

~~~
cutadapt -a AGATCGGAAGAGCGTCGTGTAGGGAAAGAatctacacttagtagaaatt -m20 –
discard-untrimmed
bowtie2 -x new.primers.BC.only -p 12 --very-sensitive –nofw
samtools view -bS Input.umi.sam > Input.umi.bam
samtools sort Input.umi.bam -o Input.umi_Sorted.bam
samtools index Input.umi_Sorted.bam
samtools idxstats Input.umi_Sorted.bam > Input.umi.idxstats.txt
~~~

Only reads with a perfect match to a barcode sequence were considered. MaGeck was then run, using control normalization to the non-targeting crRNAs, on the counts table to quantify barcode enrichment in high and low bins compared to bulk unsorted cells. High and low for each condition were analyzed separately. Raw read counts and log_2_ fold changes for each pre-crRNA can be found in **Supp. Table 3**.

To assess whether barcode sequences were paired with the correct crRNA array sequences, barcode sequences were extracted from each read using UMI-tools. Reads were then trimmed using Cutadapt, and read 2 was aligned to a custom crRNA array reference using Bowtie2. Samtools was used to create a sorted bam file, and then UMI-tools was used to group reads by barcode sequence. The specific parameters used were

~~~
umi_tools extract --extract-method=regex
--bc-pattern=’^(?P<umi_1>.{8})(?P<discard_2>TCGG)(?P<umi_2>.{8})’
cutadapt -g ^NNNNNNNatcctggtattggtctgcgaaatttctactaagtgtagat -m70 –
discard-untrimmed
bowtie2 -x new.primers.array -p 12 --norc -N 0 --score-min C,0,0
samtools view -bS Input.umi.sam > Input.umi.bam
samtools sort Input.umi.bam -o Input.umi_Sorted.bam
samtools index Input.umi_Sorted.bam
samtools idxstats Input.umi_Sorted.bam > Input.umi.idxstats.txt
Umi_tools group -I Input.umi_Sorted.bam --group-
out=Input.umi_counts.txt --output-bam -S Input.umi_grouped.bam
~~~

For each pre-crRNA array, the number of reads associated with the correct barcode was determined and the percentage of reads with the correct array was calculated as 100 x (# of reads for correct array/ # total reads)

For the fitness screen, FastQ files were generated using BCLConvert in BaseSpace. To align and count pre-crRNA array sequences, reads were first trimmed used cutadapt, aligned to a custom reference using Bowtie2, and then counted using samtools. Only perfect matches to the reference were considered. The specific parameters used were:

~~~
cutadapt -g ^NNNNNNNatcctggtattggtctgcgataatttctactcttgtagat -m70 –
discard-untrimmed
 -G ^CGTAatctacaagagtagaaatta -m70 --discard-untrimmed
bowtie2 -x As -p 12 --end-to-end -N 0 --score-min C,0,0 -I 152 -X 152
samtools view -bS As.Plasmid.sam > As.Plasmid.bam
samtools sort As.Plasmid.bam -o As_Sorted.Plasmid.bam
samtools index As_Sorted.Plasmid.bam
samtools idxstats As_Sorted.Plasmid.bam > As.Plasmid.idxstats.txt
~~~

To model changes in pre-crRNA array abundance over time, we used DESeq2 with the number of days in culture included as an independent variable in the model. We subtracted five from the number of days such that our initial observation at day 5 would serve as the intercept in the fit models. We evaluated statistical significance using a likelihood ratio test compared to a model without the day parameter. We estimated false discovery rates using the Benamini-Hochberg method as implemented in DESeq2. We modeled each screen independently, and plotted results using data where counts for all four screens were jointly normalized for differences in read depth by DESeq2. Code and examples for the analysis is available at https://github.com/ReddyLab/Cas12a_screen_analysis.

We also used our time course data in conjunction with the MAGeck MLE to determine which genes were essential and depleted over time using information from all pre-crRNAs targeting each gene. The associated beta scores and wald FDR values for each essential gene and cell surface control are reported in **Supp. Table 2** and **Fig. 6**. Gene effect score data in **Supp. Fig. 10** is from the DepMap Public 25Q1 dataset (depmap.org)

## Data Availability

Supplementary data has been made available with this study. Raw sequence data and associated processed files have been deposited in NCBI’s Gene Expression Omnibus (Edgar et al., 2002) and are accessible through GEO Series accession number GSE278336 (https://www.ncbi.nlm.nih.gov/geo/query/acc.cgi?acc=GSE278336).

## Code Availability

Code and examples for the Deseq2 analysis of the fitness screen data is available at https://github.com/ReddyLab/Cas12a_screen_analysis.

