## Supplemental Information for "HyperCas12a enables multiplexed CRISPRi screens"

#### Supplemental Figures

##### Supplemental Figure 1

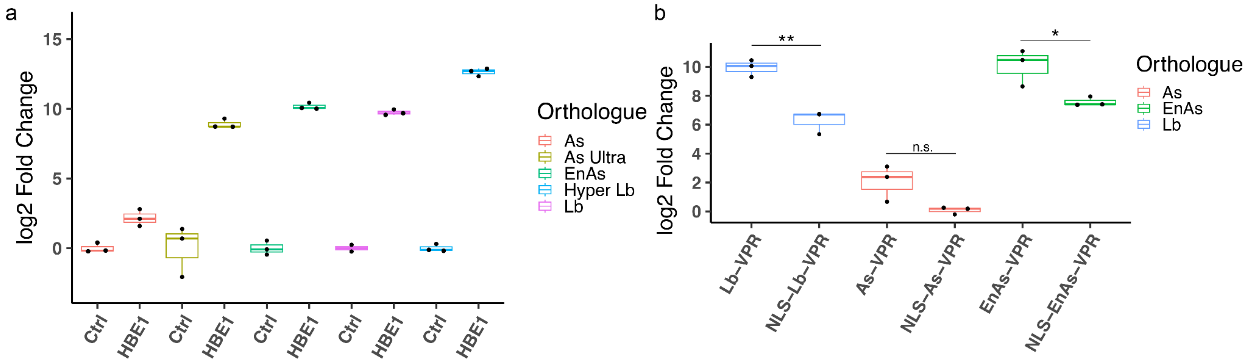


**a,** Comparison of dCas12a activators. dCas12a-VPR variants were compared and treated as in **Fig. 1b**, except with 1/10th the amount of crRNA plasmid transfected (low crRNA condition) (two or more independent experiments per transfection condition) **b,** Comparison of dCas12a activators with or without N-terminal NLS fusion. dCas12a activators for three different Cas12a variants were created with or without an N-terminal SV40 NLS. The As and Lb dCas12a-VPR fusions were transfected into HEK293T cells with or without a plasmid containing a crRNA targeting the *HBB* promoter (HEK293T nTPM = 0.0), while the dEnAsCas12a-VPR construct was co-transfected with either a crRNA targeting the *HBB* promoter or a crRNA targeting the *AR* promoter to serve as a negative control. After two days, RNA was harvested and relative *HBB* expression levels were quantified using RT-qPCR (n = 3 independent experiments per transfection condition) * $p < 0.05$, ** $p < 0.01$, two-sided student’s t-test

### Supplemental Figure 2

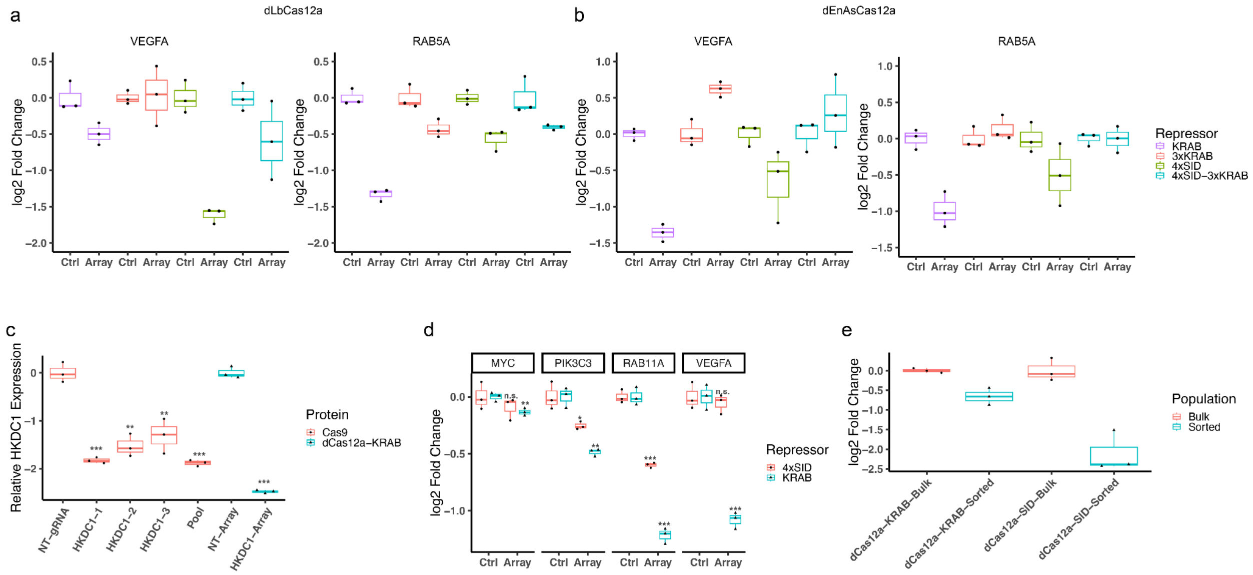


**a,** Four different dLbCas12a repressor fusions were co-transfected in HEK293T cells with a pre-crRNA of three crRNAs targeting the *VEGFA* (nTPM = 9.7) or *RAB5A* (nTPM = 22.6) promoter or a control pre-crRNA containing a crRNA targeting the *HBB* promoter. After two days, RNA was harvested and target gene expression was assessed using RT-qPCR (n = 3 independent experiments per transfection condition) **b,** Four different dEnAsCas12a repressor fusions were evaluated as in **a** (n = 3 independent experiments per transfection condition) **c,** *HKDC1* expression in HepG2 knockdown and knockout cell lines. *HKDC1* (nTPM = 17.1) knockout cell lines and controls were created by transducing HepG2 cells with Cas9 and a non-targeting gRNA, one of three *HKDC1*-targeting gRNAs, or a pool of three *HKDC1* gRNAs (pool). The *HKDC1* knockdown cell line and control were created by transducing HepG2 cells that stably express dHyperLbCas12a-KRAB with either an array of four crRNAs targeting the *HKDC1* promoter or a control array of four non-targeting crRNAs. After selecting with puromycin for 11 days, RNA was harvested and *HKDC1* expression was assayed using RT-qPCR (n = 3 independent experiments per transfection condition, n.s. Non-significant, * $p \leq 0.05$, ** $p \leq 0.01$, *** $p \leq 0.001$, FDR-adjusted two-sided student's t-test) **d,** A549 cells stably expressing dHyperLbCas12a-KRAB or dHyperLbCas12a-4xSID were transduced with the multiplexed pre-crRNA from **Fig 2e**. Cells were selected with 0.75ug/mL puromycin for eight days before RNA was harvested and target gene expression was assayed using RT-qPCR (n = 3 independent experiments per transfection condition, n.s. Non-significant, * $p \leq 0.05$, ** $p \leq 0.01$, *** $p \leq 0.001$, FDR-adjusted two-sided student's t-test)). **e,** Relative dHyperLbCas12a repressor expression in stable A549 cell lines that were transfected with the pre-crRNA from **d**. Repressor transcript levels were assayed both in bulk transfected cells and in cells sorted on array expression (at least two independent experiments per transfection condition)

#### Supplemental Figure 3:

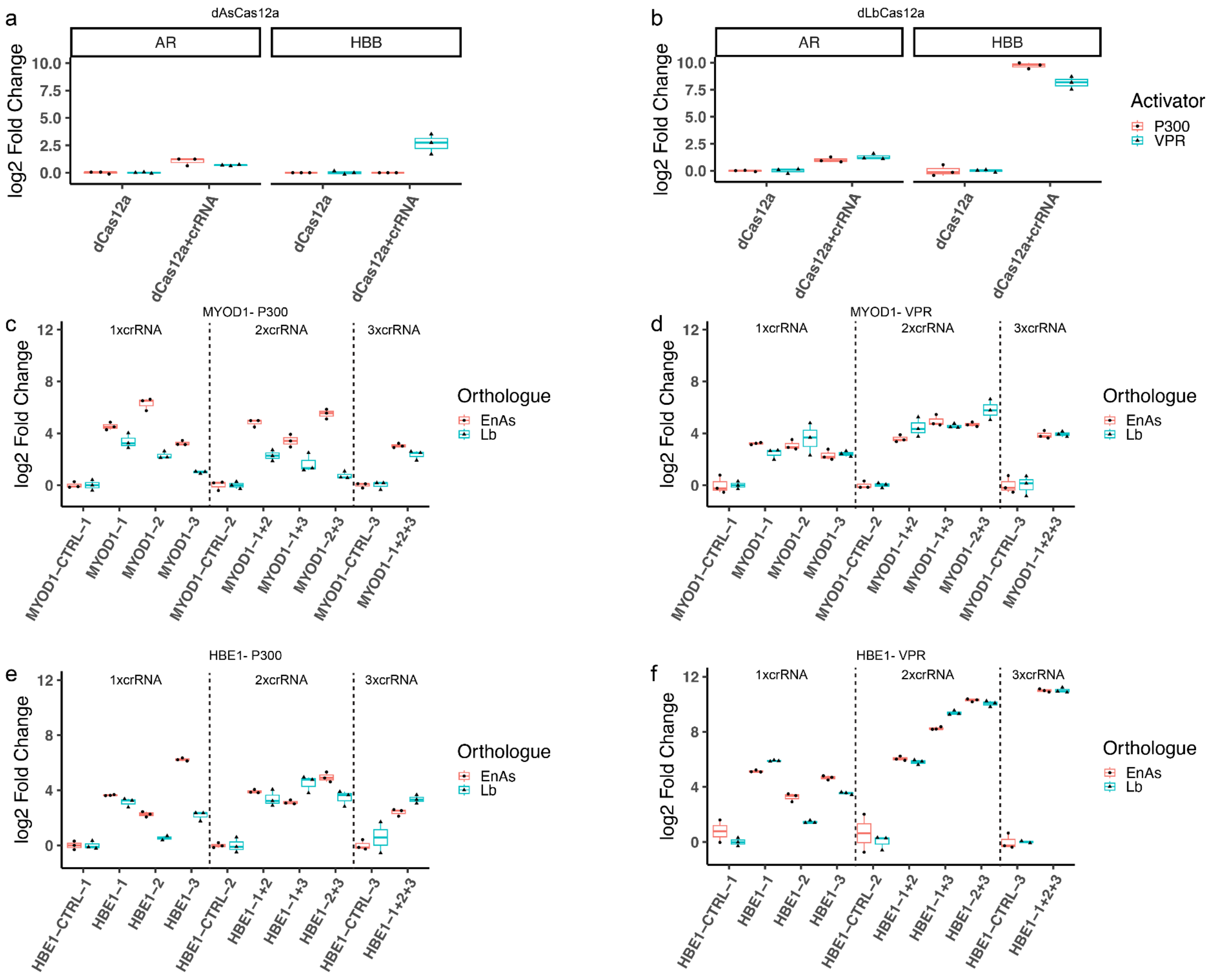


**a,b** Gene activation by wild type Cas12a activators using a single crRNA **a,** dAsCas12a or **b,** dLbCas12a activators were transfected into HEK293T cells with or without a single crRNA targeting either the *AR* (nTPM = 2.2) or *HBB* promoter (nTPM = 0.0) (crRNA sequences first reported in Ref. 46). Spacer sequences used were originally reported in Ref. 46. After two days, RNA was harvested and target gene expression was assessed using RT-qPCR (n = 3 independent experiments per transfection condition) **c,d** *MYOD1* (nTPM = 0.0) activation by different combinations of crRNAs. Three crRNAs were designed against the *MYOD1* promoter, and pre-crRNAs with all 1-3 way combinations of these crRNAs were constructed. These pre-crRNAs or pre-crRNAs with an equal number of non-targeting crRNAs were cotransfected with either **c,** dCas12a-P300 or **d,** dCas12a-VPR fusions and gene expression was assessed as in **a** (n = 3 independent experiments per transfection condition). **e,f** *HBE1* (nTPM = 0.1) activation by different combinations of crRNAs.Three crRNAs were designed against the HBE1 promoter, and pre-crRNAs with all 1-3 way combinations of these crRNAs were constructed. These pre-crRNAs or pre-crRNAs with the same number of non-targeting crRNAs were cotransfected with either **e,** dCas12a-P300 or **f,** dCas12a-VPR fusions and gene expression was assessed as in **a** (n = 3 independent experiments per transfection condition).

#### Supplemental Figure 4:

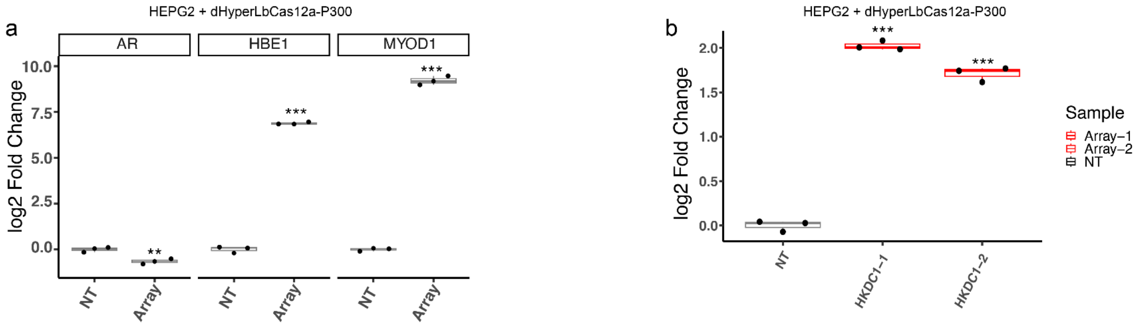


**a,** Multiplexed gene activation in a stable HEPG2 dHyperLbCas12a-P300 cell line. Cells stably expressing dHyperLbCas12a-P300 were transduced with the multiplexed crRNA array from **Fig. 4**. or a non-targeting pre-crRNA array containing ten non-targeting crRNAs. Cells were selected with puromycin, and gene expression was assessed via RT-qPCR 16 days post-transduction (n = 3 independent wells per cell line) *AR* nTPM = 0.0 , *HBE1* nTPM =2.4, *MYOD1* nTPM = 0.0. **b,** Activation of *HKDC1* (nTPM = 171.) in a stable HEPG2 dHyperLbCas12a-P300 cell line. Cells stably expressing dHyperLbCas12a-P300 were transduced with two different pre-crRNA arrays, each containing two crRNAs targeting the *HKDC1* promoter, or a control pre-crRNA containing three non-targeting crRNAs. Cells were selected with puromycin, and gene expression was assessed via RT-qPCR 16 days post-transduction (n = 3 independent wells per cell line). N.s. not significant, *** p < 0.001, FDR-adjusted Student's one-sided t-test

#### Supplemental Figure 5:

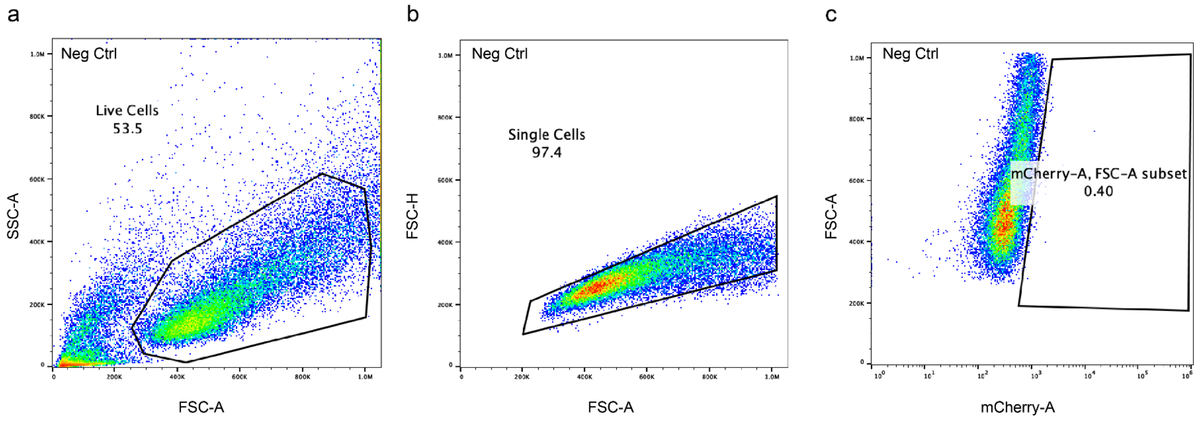


Flow cytometry gating strategy for defining mCherry positive iPSCs during neuronal differentiation experiments. A negative control that was transduced with a pre-crRNA array containing three non-targeting crRNAs was used to establish gates for **a,** live cells and **b,** single cells using FSC and SSC as shown in the first two plots. **c,** FSC and mCherry (FL2-A) signal was used to gate on mCherry positive cells.

#### Supplemental Figure 6:

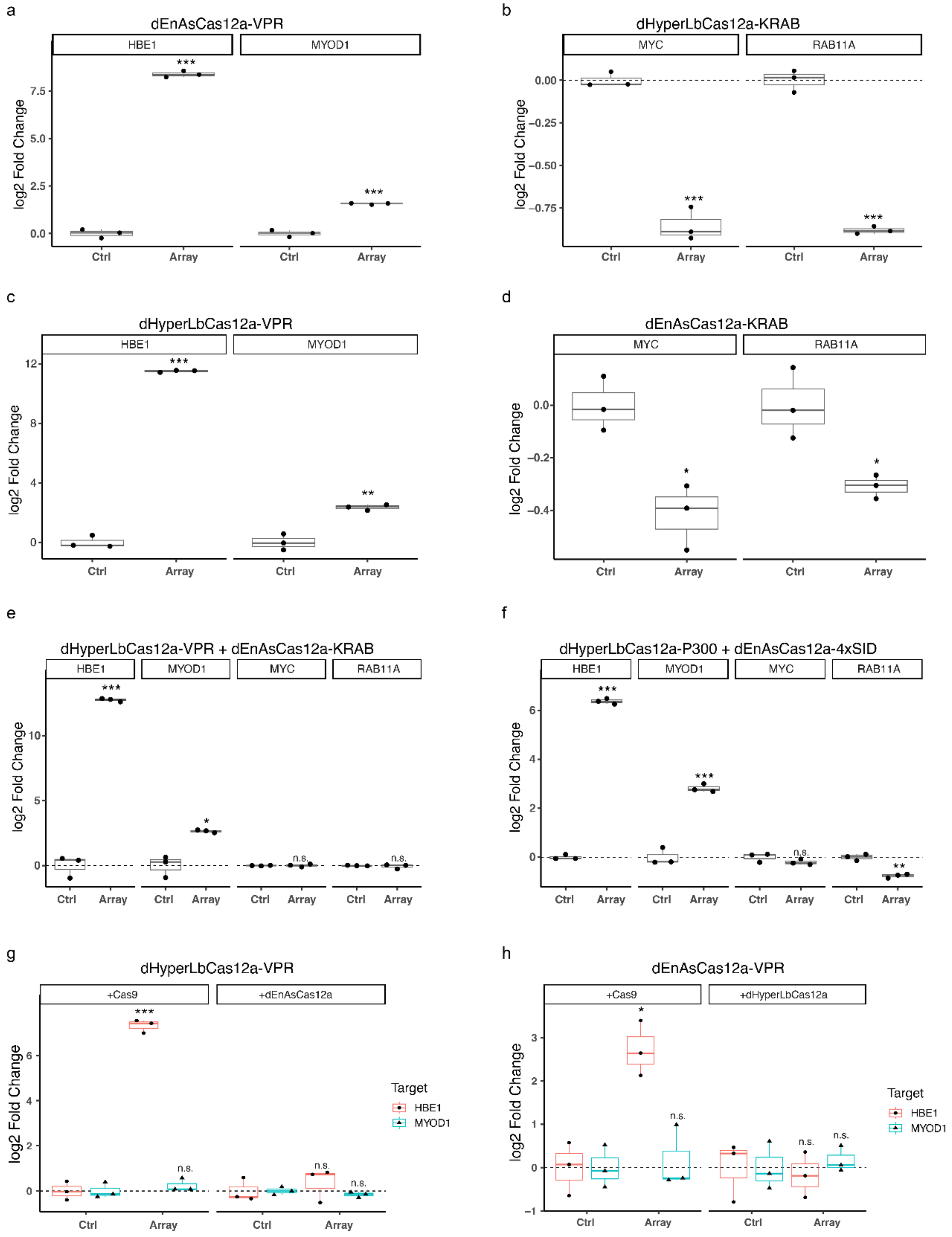

**a,** dEnAsCas12a-VPR, **b,** dHyperLbCas12a-KRAB, **c,** dHyperLbCas12a-VPR, and **d,** dEnAsCas12a-KRAB are capable of using a hybrid pre-crRNA in order to modulate gene expression. dCas12a fusions were co-transfected into HEK293T cells with either the hybrid pre-crRNA from **Fig. 5** or a control pre-crRNA of 10 non-targeting crRNAs. After 2 days, cells were sorted on GFP as a marker of Cas12a expression, and gene expression changes were assessed as before (n = 3 independent experiments per transfection condition). **(e,f)** Simultaneous activation and repression by **(e)** dHyperLbCas12a-VPR and dEnAsCas12a-KRAB or **(f)** dHyperLbCas12a-P300 and dEnAsCas12a-4xSID. Gene expression was assessed as in **(a-d)** (n = 3 independent experiments per transfection condition). **(g,h)** Ability of **(g)** dHyperLbCas12a-VPR and **(h)** dEnAsCas12a-VPR to use orthologous crRNAs for activation in the presence of either Cas9 or the autologous dCas12a. Plasmids were co-transfected into HEK293T cells and sorted on GFP as a marker of dCas12a-VPR expression, and gene expression changes were assessed as before (n = 3 independent experiments per transfection condition). N.s. not significant, *** $p < 0.001$, FDR-adjusted Student's two-sided t-test

### Supplemental Figure 7:

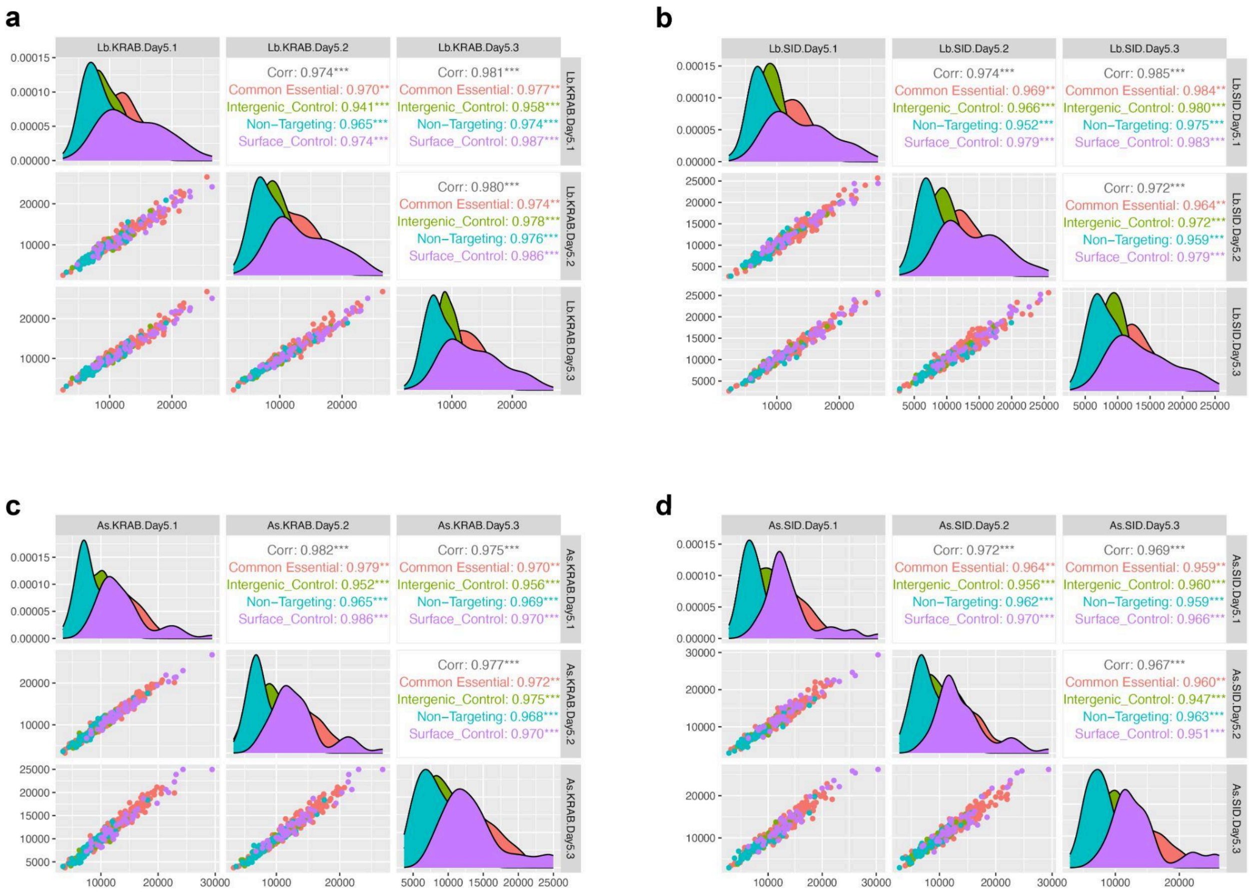


Correlation of initial (day 5) normalized read counts between replicates for **a,** dHyperLbCas12a-KRAB, **b,** dHyperLbCas12a-4xSID, **c,** dEnAsCas12a-KRAB, and **d,** dEnAsCas12a-4xSID

#### Supplemental Figure. 8:

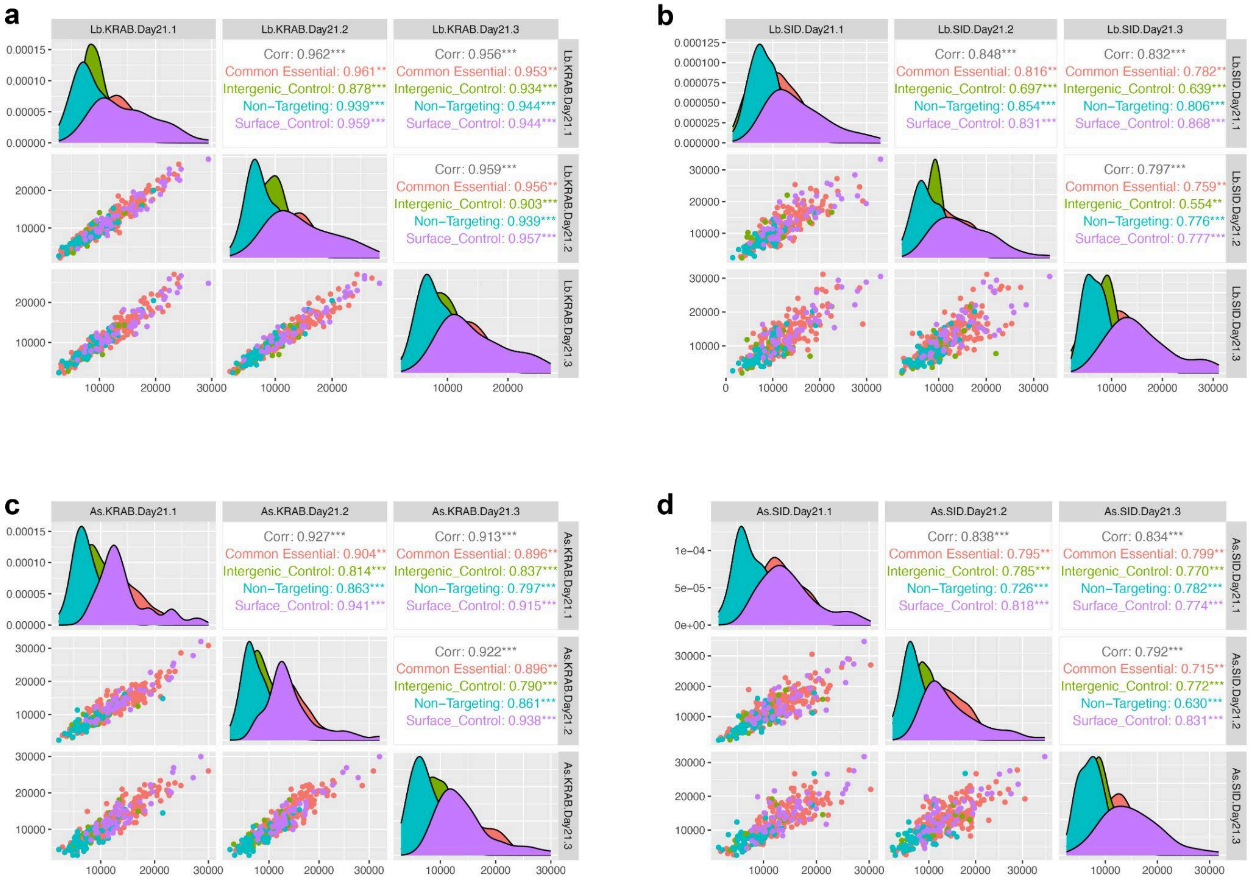


Correlation of final (day 21) normalized read counts between replicates for **a,** dHyperLbCas12a-KRAB, **b,** dHyperLbCas12a-4xSID, **c,** dEnAsCas12a-KRAB, and **d,** dEnAsCas12a-4xSID

#### Supplemental Figure 9:

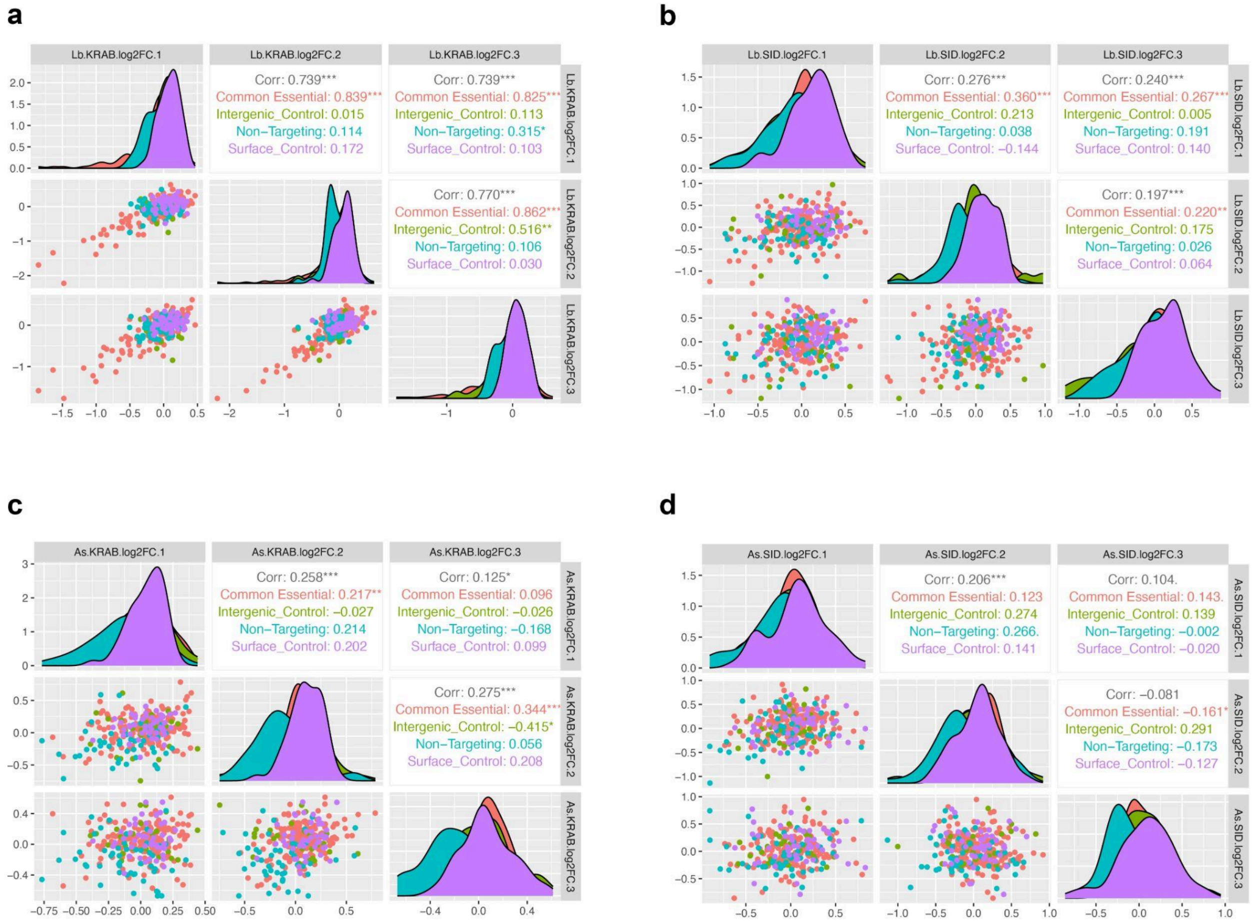


Correlation of log2 fold change of normalized read counts from Day 5 to Day 21 between replicates for **a,** dHyperLbCas12a-KRAB, **b,** dHyperLbCas12a-4xSID, **c,** dEnAsCas12a-KRAB, and **d,** dEnAsCas12a-4xSID

**Supplemental Figure 10:**

a

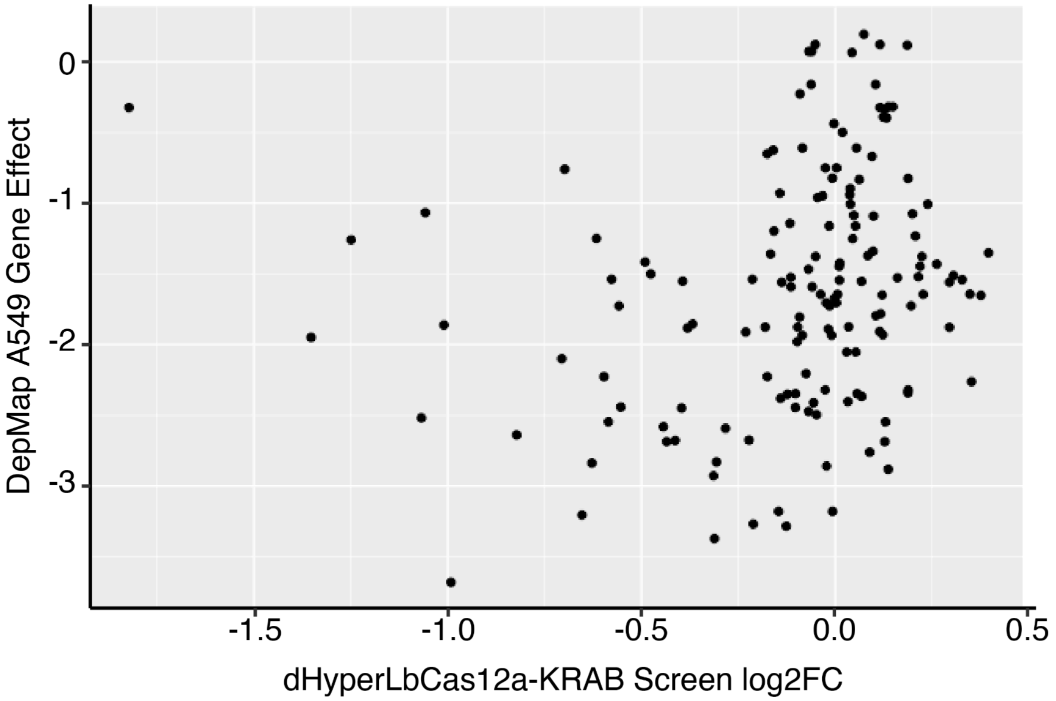


**a,** Comparison between log2 fold changes for pre-crRNA arrays targeting essential genes (day 5 vs. day 21) and DepMap gene effect scores for these genes in A549s. A gene effect of zero represents a gene with no effect on cell fitness while more negative scores represent essential genes. A gene effect of -1 corresponds to the median of all common essential genes. 131/145 essential genes targeted were present in both datasets.

#### Supplemental Figure 11:

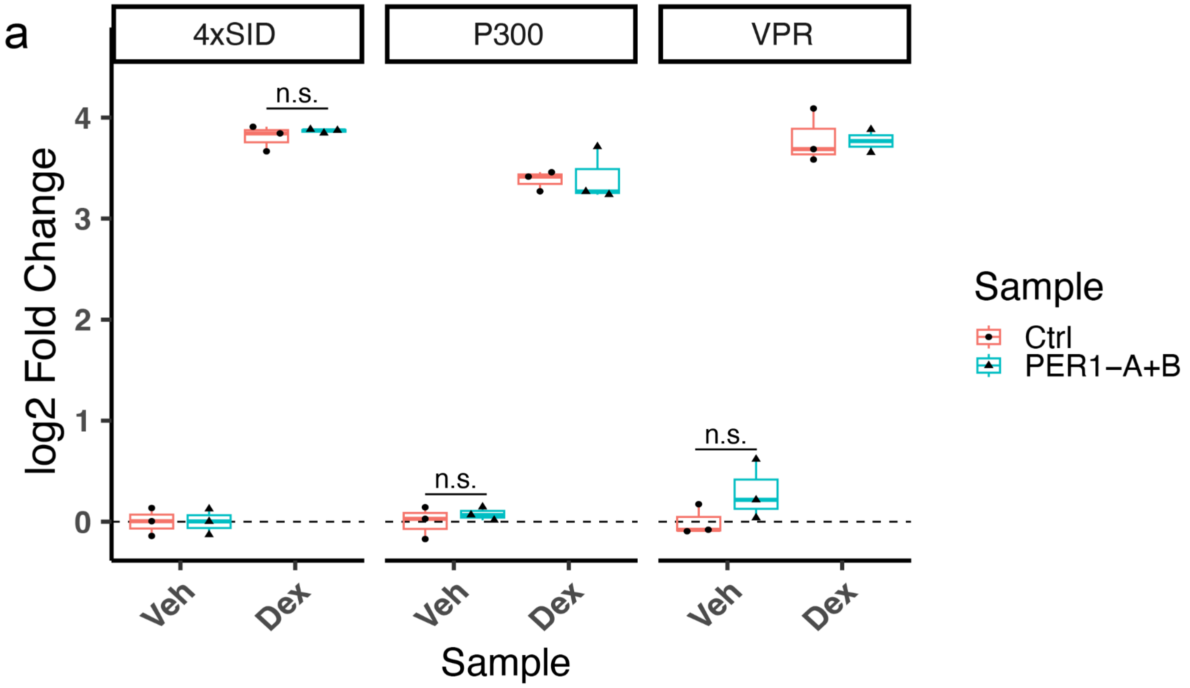


**a,** A549 cells stably expressing dHyperLbCas12a- 4xSID, P300, or VPR fusions were transduced with either a control pre-crRNA of four non-targeting crRNAs or a pre-crRNA of four crRNAs containing two crRNAs targeting each *PER1* enhancer (*PER1* nTPM = 6). Cells were selected with puromycin for eight days and then treated with 100nM dexamethasone or an ethanol control for three hours before RNA was harvested and RT-qPCR was performed to assess changes in *PER1* expression (at least 2 independent experiments per cell line and drug condition). N.S.- No significant change in *PER1* induction after treatment with dexamethasone between control pre-crRNA arrays and pre-crRNAs targeting the two *PER1* enhancers.

#### Supplemental Figure 12:

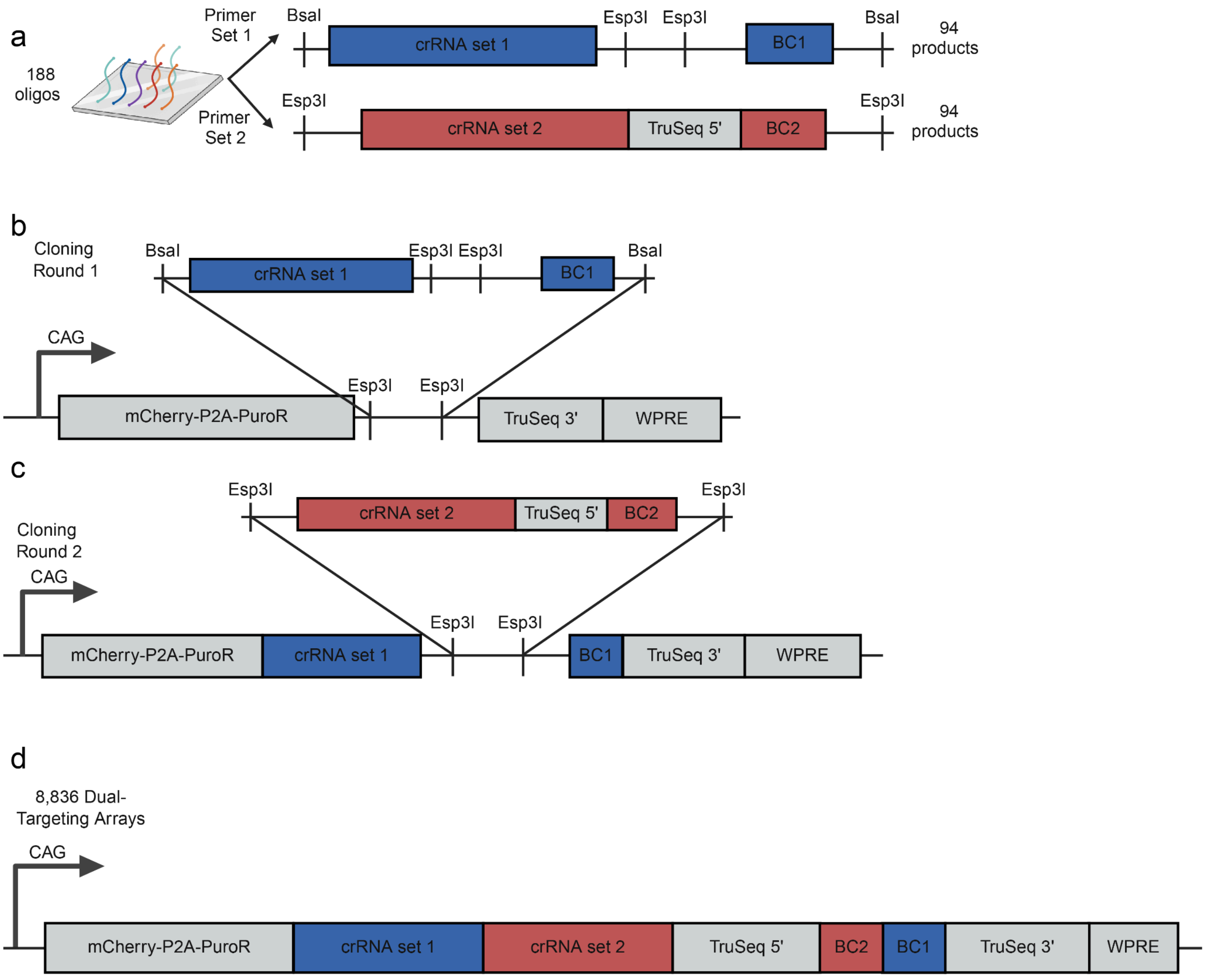


**a,** oligos containing sets of 3 crRNAs plus a barcode are synthesized. Two different pairs of primers are used to amplify crRNA group 1 (n = 94 oligos) and crRNA group 2 (n = 94 oligos). **b,** The crRNA group 1 pool is digested with BsaI and ligated into the crRNA expression vector which has been digested with Esp3I. **c,** The crRNA group 2 pool is cloned into the library of 94 crRNA group 1 plasmids using one pot Golden Gate cloning with Esp3I. **d,** The final product of the cloning is n = 8,836 (94x94) unique pre-crRNA arrays associated with a unique bipartite barcode that is flanked by TruSeq adapter sequences

**Supplemental Figure 13:**

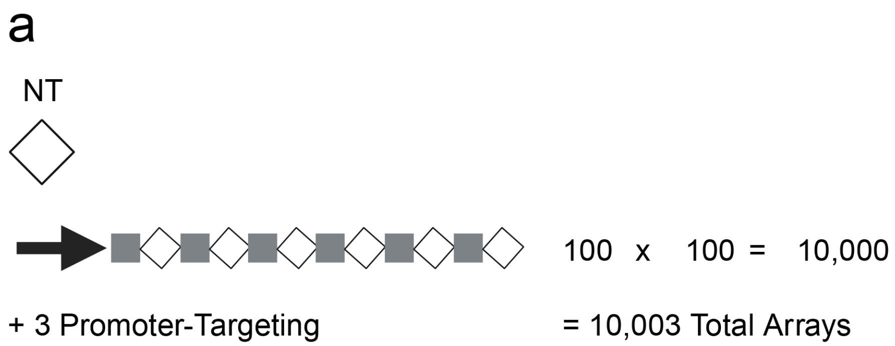


**a,** A non-targeting library of 10,003 pre-crRNAs containing six crRNAs each was assembled by using the same two-step cloning method as for the *PER1* enhancer-targeting library **(Fig 6 and Supp. Fig 5)**. Cloning 100 sets of non-targeting crRNAs in each round led to 10,000 uniquely barcoded pre-crRNAs in the final library. There were also three promoter-targeting, barcoded, crRNA arrays that were assembled after only one round of cloning and were not able to be modified after that.

#### Supplemental Figure 14:

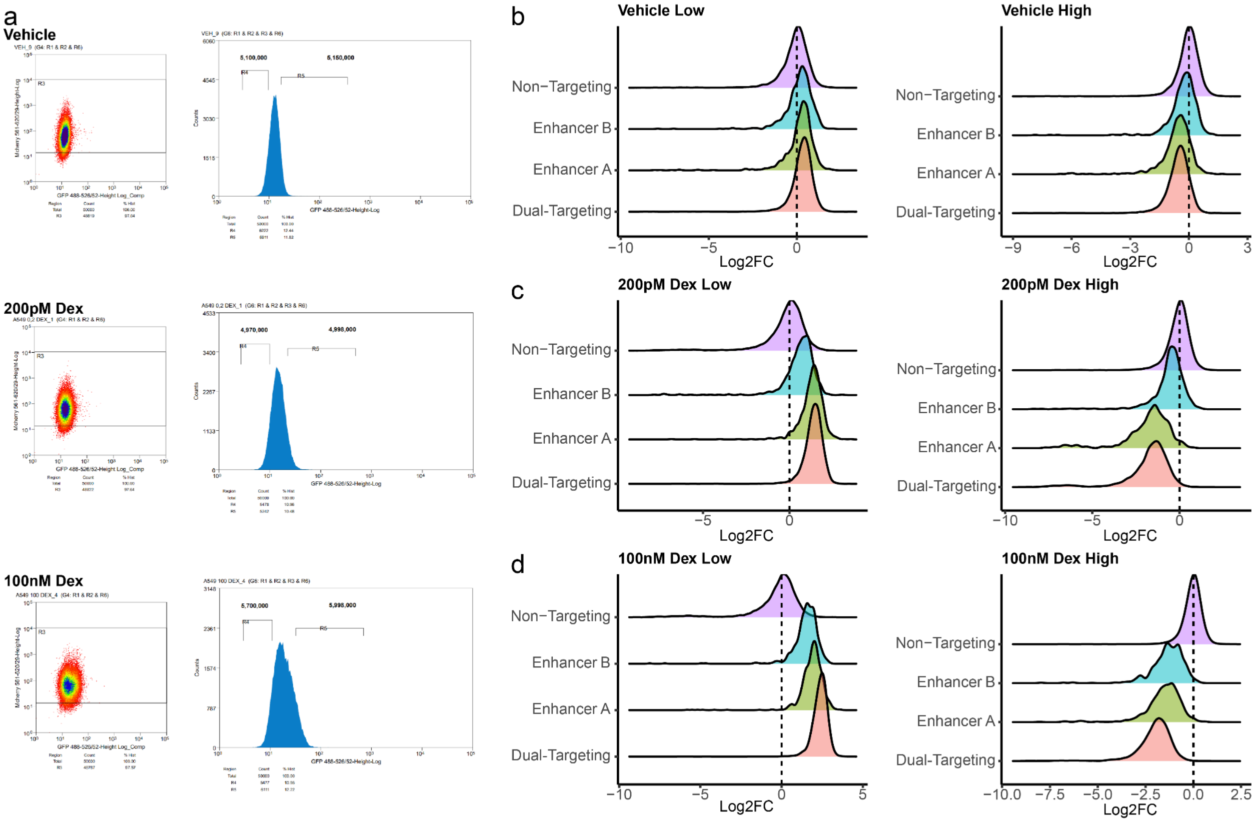


**a,** Flow cytometry plots showing the mCherry (crRNA array) and FITC (*PER1* mRNA) signal for cell populations subjected to different drug treatments. Numbers above the tails indicate the numbers of cells sorted into high and low bins. **b,c,d** density plots displaying enrichment of barcode sequences in high and low bins for **b,** cells treated with vehicle **c,** cells treated with 200pM dexamethasone and **d,** cells treated with 100nM dexamethasone

#### Supplemental Figure 15:

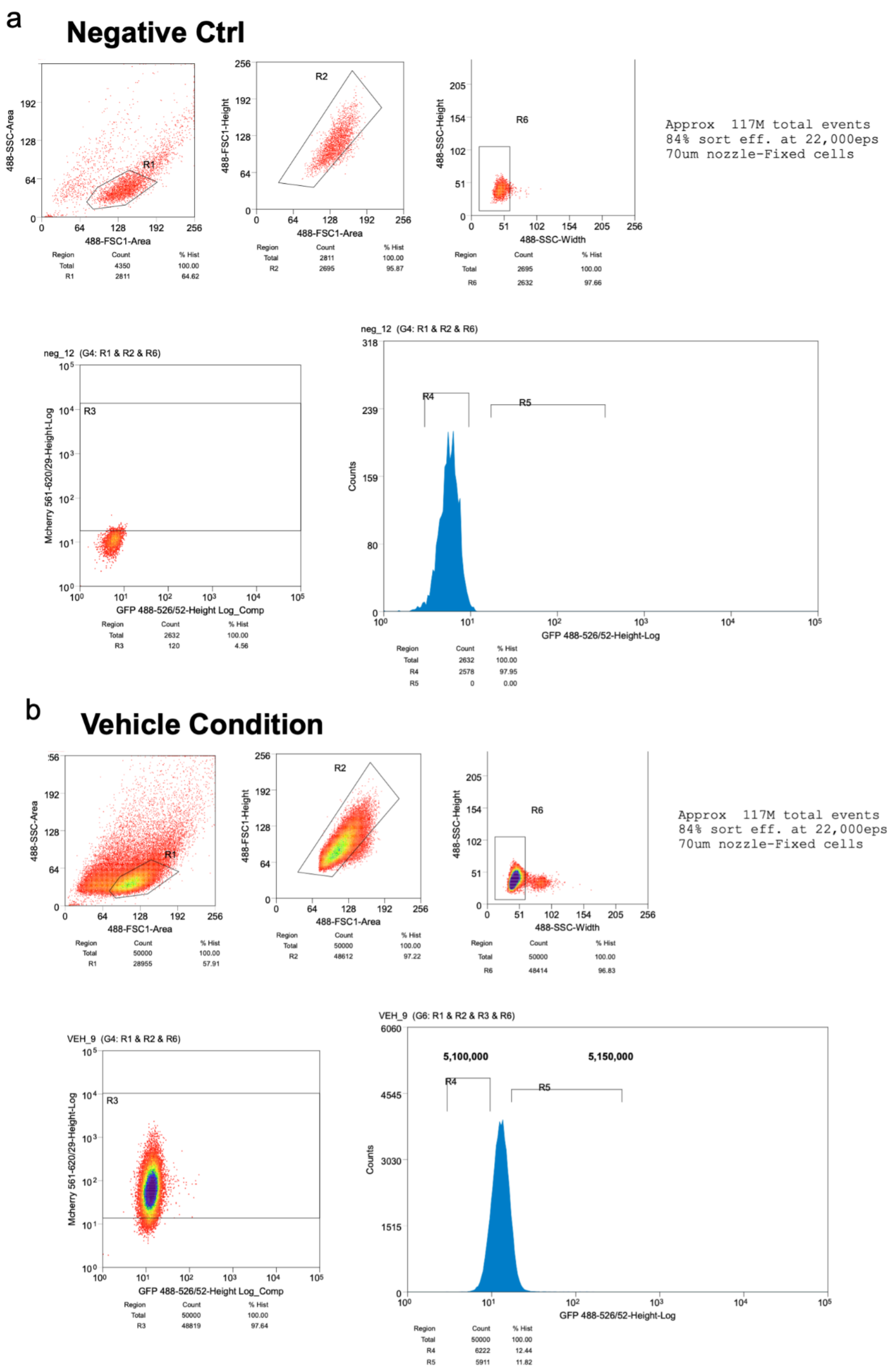


**a,** Flow cytometry gating strategy for sorting the PER1 screen. A negative control was used to establish gates for single cells using FSC and SSC as shown in the top row. It was also used to define the mCherry positive population and set thresholds for FITC signal

**b,** The same gating strategy showing the vehicle sample from the screen including gates defining the top and bottom 12% PER1 expression

**Supplemental Figure 16:**

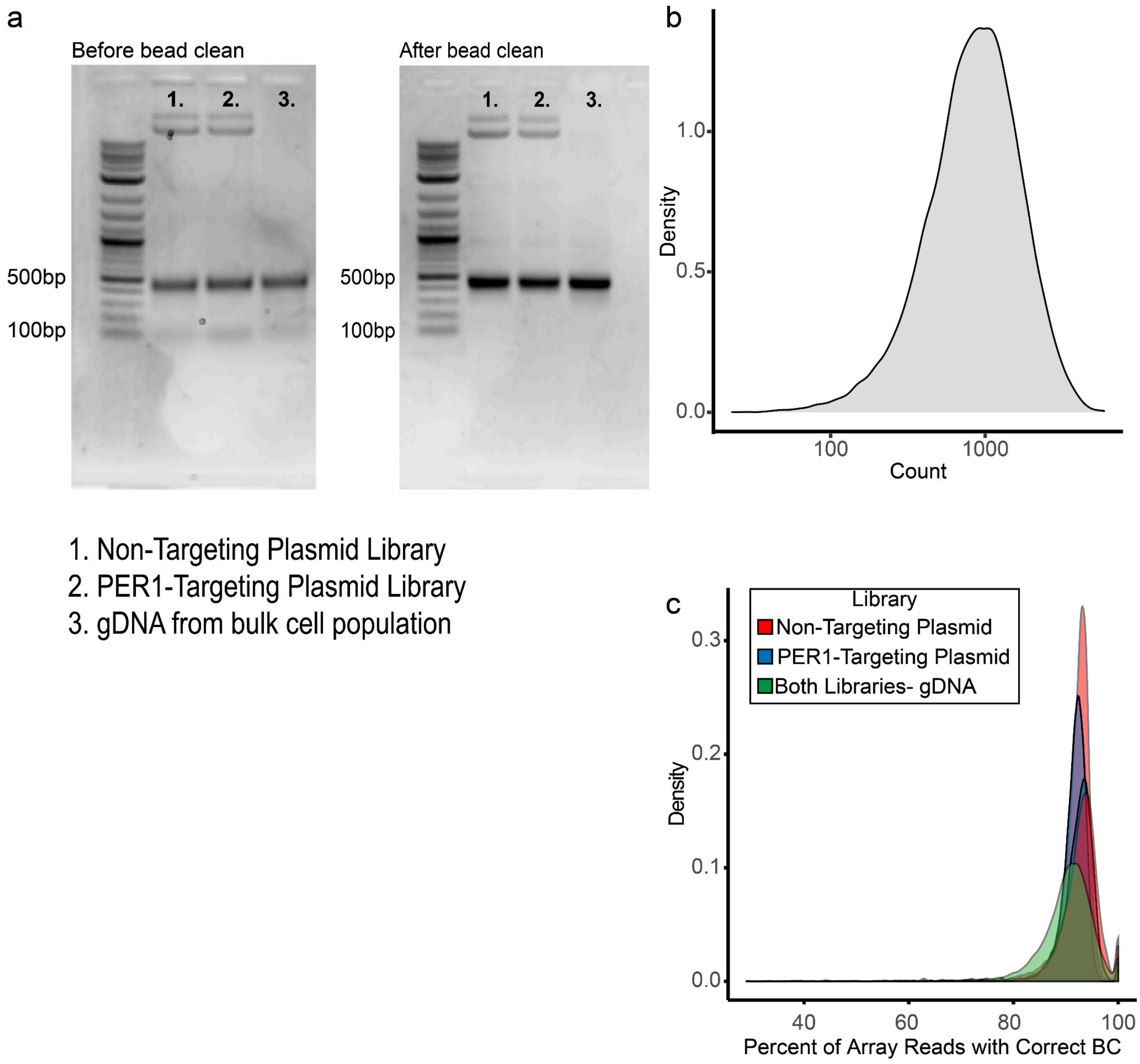


**a,** pre-crRNA arrays were amplified from either a plasmid library or genomic DNA from transduced A549 cells using PCR. PCR products were run on an agarose gel before and after cleanup using magnetic beads. **b,** Density plot showing the distribution of barcode counts from the bulk unsorted cell population. **c,** All of the sequencing reads associated with each pre-crRNA arraywere aligned and quantified. The percentage of total reads belonging to the correct barcode (BC) was quantified and is displayed here as a density plot. Each plasmid library was sequenced twice, while the genomic DNA library was sequenced once.

#### Supplemental Figure 17:

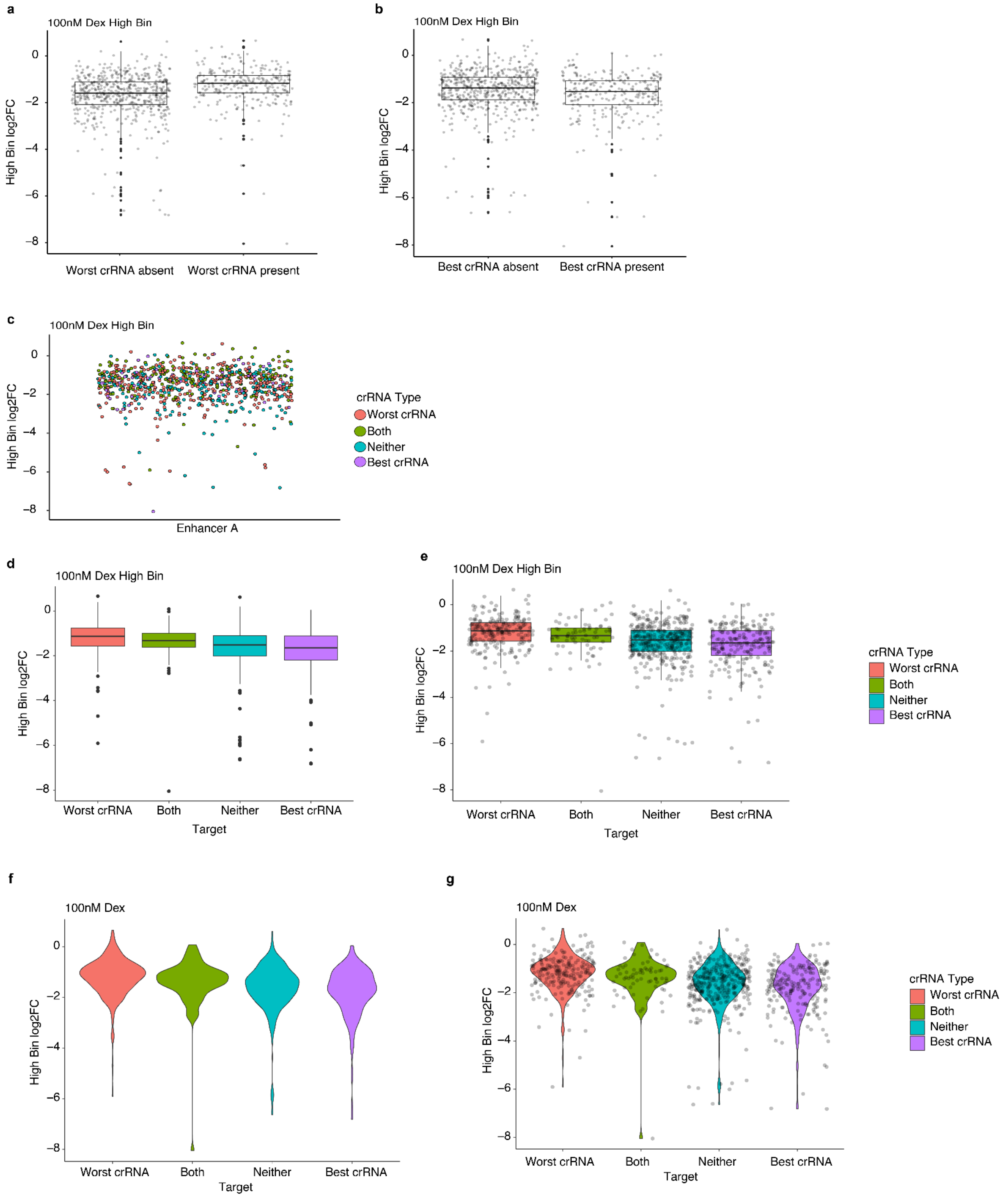


Effects of individual crRNAs on single enhancer targeting pre-crRNAs in the 100 nM Dex high bin. **a,** Boxplot showing the distribution of log2 fold changes when the worst crRNA targeting enhancer A is absent (left) or present (right) in an array. Average log2 fold change increases by 0.44 when the worst crRNA is present. **b,** Boxplot showing the distribution of log2 fold changes when the best crRNA targeting enhancer A is absent (left) or present (right) in an array. Average log2 fold change decreases by 0.22 when the best crRNA is present. **(c,d)** The distribution of log2 fold changes for pre-crRNAs containing the worst crRNA only, the best crRNA only, both, or neither as **(c)** a jitter plot or **(d)** as boxplots. There is a difference of 0.59 between the average log2 fold change of pre-crRNAs containing the worst crRNA only and those containing the best crRNA only.

**Supplemental Figure 18:**

a

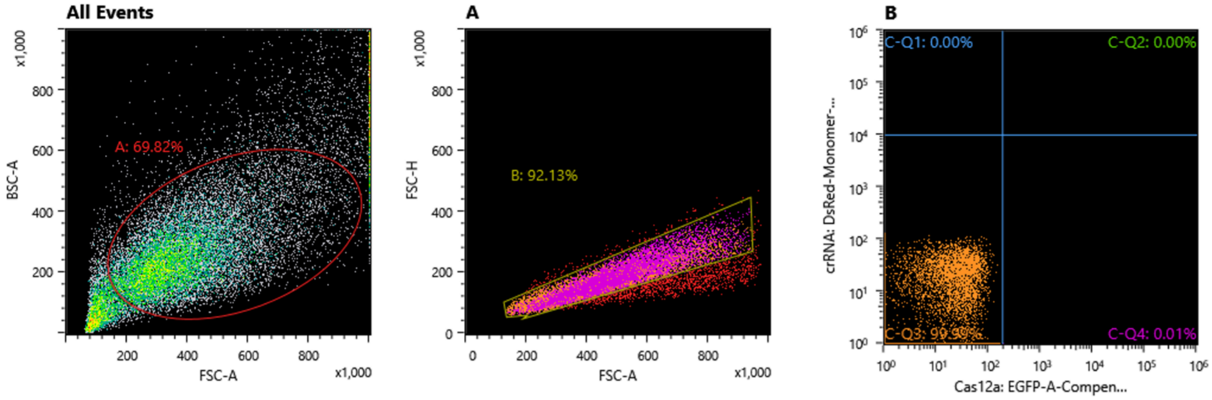


b

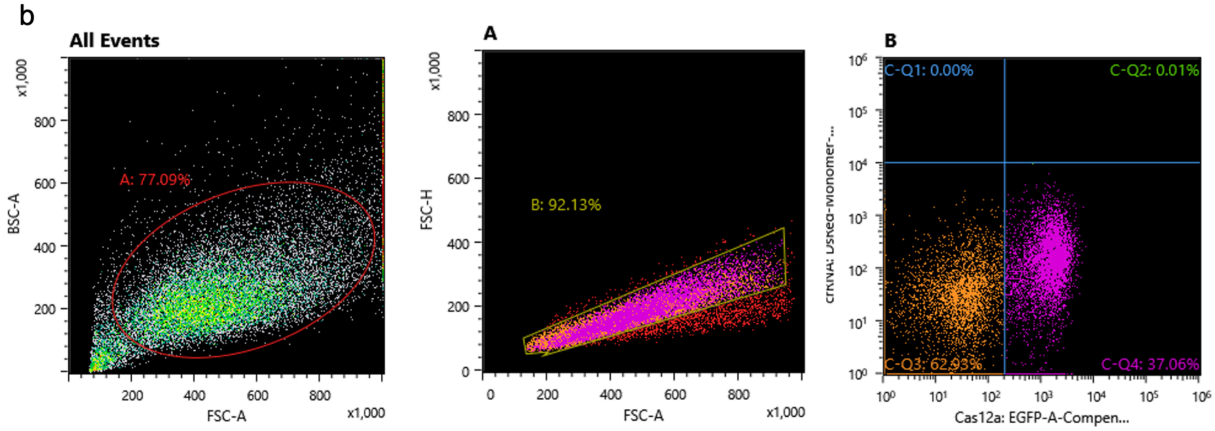


**a,** Flow cytometry gating strategy for sorting HEK293T cells transfected with Cas12a and crRNA expression plasmids. A negative control was used to establish gates for single cells using FSC and BSC as shown in the first two plots. It was also used to define the GFP positive population

**b,** The same gating strategy showing a sample transfected with dCas12a and crRNA expression plasmids. Some crRNA plasmids used in these studies expressed mCherry, but this ignored and only GFP positive cells were sorted.

#### Supplemental Figure 19:

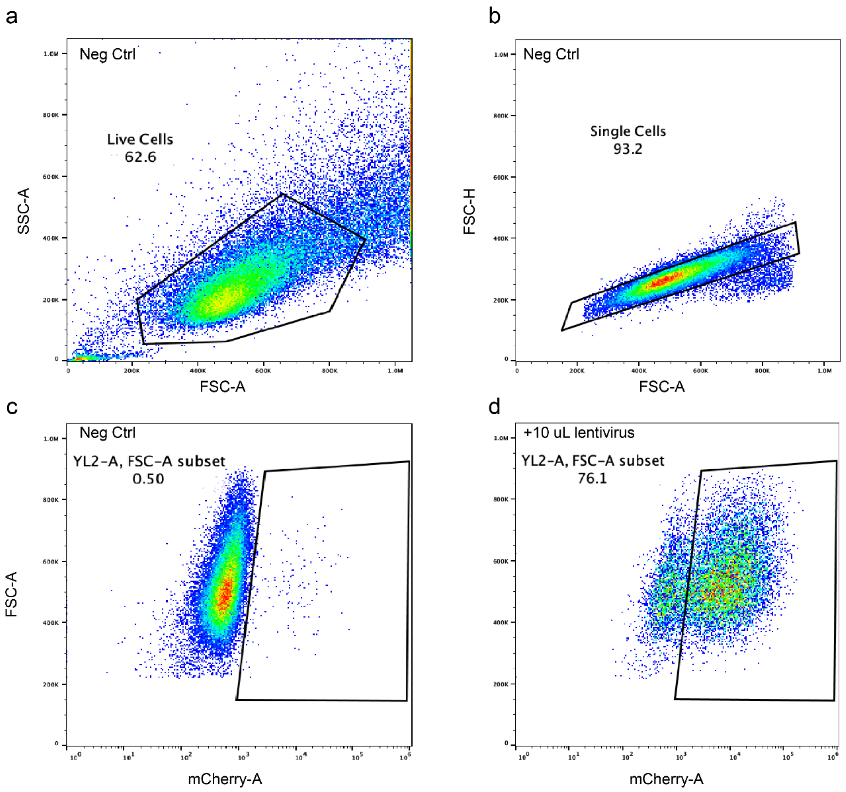


Flow cytometry gating strategy for defining mCherry positive A549 cells during lentiviral titration experiments. An untransduced negative control was used to establish gates for **a,** live cells and **b,** single cells using FSC and SSC as shown in the first two plots. **c,** This negative control was also used to establish gating for the mCherry positive population. **d,** Example of a sample transduced with a high amount of lentivirus and which has a large mCherry positive population (76% of all single cells).

### Supplemental Text

#### Barcodes remain associated with crRNA arrays throughout screening

In addition to evaluating the performance of the dHyperLbCas12a-KRAB construct, we also assessed crRNA array integrity. One concern with multiplexed screening constructs is their propensity to recombine and become uncoupled from associated barcode sequences. This is especially concerning for Cas9 dual gRNA expression vectors, which are estimated to recombine and uncouple from barcode sequences at a rate of ~10-50% during, with this recombination mostly occurring during lentivirus production[36-39]. Meanwhile, Cas12a constructs have been shown to be more stable due to their relatively short 20bp repeat sequences[71].

To assess recombination in our libraries, we amplified and sequenced the entire Cas12a pre-crRNA construct plus its barcode for each of the plasmid libraries, as well as for gDNA taken from a bulk population of transduced cells. After library amplification, we observed that amplicon size remained the same between plasmid constructs and gDNA, and also that gDNA amplification yielded a clean band of the correct size, not a smear or ladder **(Supp. Fig 13a)**. After sequencing, we extracted barcodes from reads. We recovered 18,735/18,736 crRNA arrays that were present in the initial plasmid library, and found that array abundance was uniform, as expected **(Supp. Fig 13b)**.

To assess the amount of recombination that occurred during lentivirus production, we extracted barcode sequences from reads and aligned just the pre-crRNA portion of the read to a custom reference (see methods for more details). We then grouped all the barcodes associated with each pre-crRNAs sequence and quantified the percentage of reads containing the correct barcode associated with that pre-crRNA. We performed two different sequencing runs for the plasmid libraries and found a median of 93.6% and 93.9% of reads containing the correct pre-crRNA-barcode pair for the non-targeting library, and 92.7% and 92.9% of reads containing the correct pre-crRNA-barcode pair for the PER1-targeting library **(Supp. Fig 13c)**, likely due to challenges with aligning highly homologous, repetitive sequences. When we performed this same analysis for the pre-crRNAs amplified from gDNA, we found a median of 90.4% of reads containing the correct array-barcode pair, which is in line with what we observe from the plasmid library and suggests that there is not a large amount of barcode swapping occurring during lentivirus production **(Supp. Fig 13c)**.

### Construct Sequences

dAsCas12a:
atgacacagttcgagggctttaccaacctgtatcaggtgagcaagacactgcggtttgagctgatcccacagggcaagaccctgaag
cacatccaggagcagggcttcatcgaggaggacaaggcccgcaatgatcactacaaggagctgaagcccatcatcgatcggatct
acaagacctatgccgaccagtgcctgcagctggtgcagctggattgggagaacctgagcgccgccatcgactcctatagaaagga
gaaaaccgaggagacaaggaacgccctgatcgaggagcaggccacatatcgcaatgccatccacgactacttcatcggccggac
agacaacctgaccgatgccatcaataagagacacgccgagatctacaagggcctgttcaaggccgagctgtttaatggcaaggtgc
tgaagcagctgggcaccgtgaccacaaccgagcacgagaacgccctgctgcggagcttcgacaagtttacaacctacttctccggc
ttttatgagaacaggaagaacgtgttcagcgccgaggatatcagcacagccatcccacaccgcatcgtgcaggacaacttccccaa
gtttaaggagaattgtcacatcttcacacgcctgatcaccgccgtgcccagcctgcgggagcactttgagaacgtgaagaaggccatc
ggcatcttcgtgagcacctccatcgaggaggtgttttccttccctttttataaccagctgctgacacagacccagatcgacctgtataacc
agctgctgggaggaatctctcgggaggcaggcaccgagaagatcaagggcctgaacgaggtgctgaatctggccatccagaaga
atgatgagacagcccacatcatcgcctccctgccacacagattcatcccccctgtttaagcagatcctgtccgataggaacaccctgtctt
tcatcctggaggagtttaagagcgacgaggaagtgatccagtccttctgcaagtacaagacactgctgagaaacgagaacgtgctg
gagacagccgaggccctgtttaacgagctgaacagcatcgacctgacacacatcttcatcagccacaagaagctggagacaatca
gcagcgccctgtgcgaccactgggatacactgaggaatgccctgtatgagcggagaatctccgagctgacaggcaagatcaccaa
gtctgccaaggagaaggtgcagcgcagcctgaagcacgaggatatcaacctgcaggagatcatctctgccgcaggcaaggagct
gagcgaggccttcaagcagaaaaccagcgagatcctgtcccacgcacacgccgccctggatcagccactgcctacaaccctgaa
gaagcaggaggagaaggagatcctgaagtctcagctggacagcctgctgggcctgtaccacctgctggactggtttgccgtggatga
gtccaacgaggtggaccccgagttctctgcccggctgaccggcatcaagctggagatggagccttctctgagcttctacaacaaggc
cagaaattatgccaccaagaagccctactccgtggagaagttcaagctgaactttcagatgcctacactggcctctggctgggacgtg
aataaggagaagaacaatggcgccatcctgtttgtgaagaacggcctgtactatctgggcatcatgccaaagcagaagggcaggta
taaggccctgagcttcgagcccacagagaaaaccagcgagggctttgataagatgtactatgactacttccctgatgccgccaagat
gatcccaaagtgcagcacccagctgaaggccgtgacagcccactttcagacccacacaacccccatcctgctgtccaacaatttcat
cgagcctctggagatcacaaaggagatctacgacctgaacaatcctgagaaggagccaaagaagtttcagacagcctacgccaa
gaaaaccggcgaccagaagggctacagagaggccctgtgcaagtggatcgacttcacaagggattttctgtccaagtataccaaga
caacctctatcgatctgtctagcctgcggccatcctctcagtataaggacctgggcgagtactatgccgagctgaatcccctgctgtacc
acatcagcttccagagaatcgccgagaaggagatcatggatgccgtggagacaggcaagctgtacctgttccagatctataacaag
gactttgccaagggccaccacggcaagcctaatctgcacacactgtattggaccggcctgttttctccagagaacctggccaagaca
agcatcaagctgaatggccaggccgagctgttctaccgccctaagtccaggatgaagaggatggcacaccggctgggagagaag
atgctgaacaagaagctgaaggatcagaaaaccccaatccccgacaccctgtaccaggagctgtacgactatgtgaatcacagac
tgtcccacgacctgtctgatgaggccagggccctgctgcccaacgtgatcaccaaggaggtgtctcacgagatcatcaaggataggc
gctttaccagcgacaagttctttttccacgtgcctatcacactgaactatcaggccgccaattccccatctaagttcaaccagagggtga
atgcctacctgaaggagcaccccgagacacctatcatcggcatcgcccggggcgagagaaacctgatctatatcacagtgatcgac
tccaccggcaagatcctggagcagcggagcctgaacaccatccagcagtttgattaccagaagaagctggacaacagggagaag
gagagggtggcagcaaggcaggcctggtctgtggtgggcacaatcaaggatctgaagcagggctatctgagccaggtcatccacg
agatcgtggacctgatgatccactaccaggccgtggtggtgctggagaacctgaatttcggctttaagagcaagaggaccggcatcg
ccgagaaggccgtgtaccagcagttcgagaagatgctgatcgataagctgaattgcctggtgctgaaggactatccagcagagaaa
gtgggaggcgtgctgaacccataccagctgacagaccagttcacctcctttgccaagatgggcacccagtctggcttcctgttttacgtg
cctgccccatatacatctaagatcgatcccctgaccggcttcgtggaccccttcgtgtggaaaaccatcaagaatcacgagagccgca
agcacttcctggagggcttcgactttctgcactacgacgtgaaaaccggcgacttcatcctgcactttaagatgaacagaaatctgtcctt
ccagaggggcctgcccggctttatgcctgcatgggatatcgtgttcgagaagaacgagacacagtttgacgccaagggcacccctttc
atcgccggcaagagaatcgtgccagtgatcgagaatcacagattcaccggcagataccgggacctgtatcctgccaacgagctgat

cgccctgctggaggagaagggcatcgtgttcagggatggctccaacatcctgccaaagctgctggagaatgacgattctcacgccat
cgacacgatggtggccctgatccgcagcgtgctgcagatgcggaactccaatgccgccacaggcgaggactatatcaacagcccc
gtgcgcgatctgaatggcgtgtgcttcgactcccggtttcagaacccagagtggccaatggacgccgatgccaatggcgcctaccac
atcgccctgaagggccagctgctgctgaatcacctgaaggagagcaaggatctgaagctgcagaacggcatctccaatcaggact
ggctggcctacatccaggagctgcgcaac

dEnAsCas12a:
Atgacacagttcgagggctttaccaacctgtatcaggtgagcaagacactgcggtttgagctgatcccacagggcaagaccctgaag
cacatccaggagcagggcttcatcgaggaggacaaggcccgcaatgatcactacaaggagctgaagcccatcatcgatcggatct
acaagacctatgccgaccagtgcctgcagctggtgcagctggattgggagaacctgagcgccgccatcgactcctatagaaagga
gaaaaccgaggagacaaggaacgccctgatcgaggagcaggccacatatcgcaatgccatccacgactacttcatcggccggac
agacaacctgaccgatgccatcaataagagacacgccgagatctacaagggcctgttcaaggccgagctgtttaatggcaaggtgc
tgaagcagctgggcaccgtgaccacaaccgagcacgagaacgccctgctgcggagcttcgacaagtttacaacctacttctccggc
ttttatagaaacaggaagaacgtgttcagcgccgaggatatcagcacagccatcccacaccgcatcgtgcaggacaacttccccaa
gtttaaggagaattgtcacatcttcacacgcctgatcaccgccgtgcccagcctgcgggagcactttgagaacgtgaagaaggccatc
ggcatcttcgtgagcacctccatcgaggaggtgttttccttccctttttataaccagctgctgacacagacccagatcgacctgtataacc
agctgctgggaggaatctctcgggaggcaggcaccgagaagatcaagggcctgaacgaggtgctgaatctggccatccagaaga
atgatgagacagcccacatcatcgcctccctgccacacagattcatcccccctgtttaagcagatcctgtccgataggaacaccctgtctt
tcatcctggaggagtttaagagcgacgaggaagtgatccagtccttctgcaagtacaagacactgctgagaaacgagaacgtgctg
gagacagccgaggccctgtttaacgagctgaacagcatcgacctgacacacatcttcatcagccacaagaagctggagacaatca
gcagcgccctgtgcgaccactgggatacactgaggaatgccctgtatgagcggagaatctccgagctgacaggcaagatcaccaa
gtctgccaaggagaaggtgcagcgcagcctgaagcacgaggatatcaacctgcaggagatcatctctgccgcaggcaaggagct
gagcgaggccttcaagcagaaaaccagcgagatcctgtcccacgcacacgccgccctggatcagccactgcctacaaccctgaa
gaagcaggaggagaaggagatcctgaagtctcagctggacagcctgctgggcctgtaccacctgctggactggtttgccgtggatga
gtccaacgaggtggaccccgagttctctgcccggctgaccggcatcaagctggagatggagccttctctgagcttctacaacaaggc
cagaaattatgccaccaagaagccctactccgtggagaagttcaagctgaactttcagatgcctacactggccagaggctgggacgt
gaatagagagaagaacaatggcgccatcctgtttgtgaagaacggcctgtactatctgggcatcatgccaaagcagaagggcaggt
ataaggccctgagcttcgagcccacagagaaaaccagcgagggctttgataagatgtactatgactacttccctgatgccgccaaga
tgatcccaaagtgcagcacccagctgaaggccgtgacagcccactttcagacccacacaacccccatcctgctgtccaacaatttca
tcgagcctctggagatcacaaaggagatctacgacctgaacaatcctgagaaggagccaaagaagtttcagacagcctacgccaa
gaaaaccggcgaccagaagggctacagagaggccctgtgcaagtggatcgacttcacaagggattttctgtccaagtataccaaga
caacctctatcgatctgtctagcctgcggccatcctctcagtataaggacctgggcgagtactatgccgagctgaatcccctgctgtacc
acatcagcttccagagaatcgccgagaaggagatcatggatgccgtggagacaggcaagctgtacctgttccagatctataacaag
gactttgccaagggccaccacggcaagcctaatctgcacacactgtattggaccggcctgttttctccagagaacctggccaagaca
agcatcaagctgaatggccaggccgagctgttctaccgccctaagtccaggatgaagaggatggcacaccggctgggagagaag
atgctgaacaagaagctgaaggatcagaaaaccccaatccccgacaccctgtaccaggagctgtacgactatgtgaatcacagac
tgtcccacgacctgtctgatgaggccagggccctgctgcccaacgtgatcaccaaggaggtgtctcacgagatcatcaaggataggc
gctttaccagcgacaagttctttttccacgtgcctatcacactgaactatcaggccgccaattccccatctaagttcaaccagagggtga
atgcctacctgaaggagcaccccgagacacctatcatcggcatcgcccggggcgagagaaacctgatctatatcacagtgatcgac
tccaccggcaagatcctggagcagcggagcctgaacaccatccagcagtttgattaccagaagaagctggacaacagggagaag
gagagggtggcagcaaggcaggcctggtctgtggtgggcacaatcaaggatctgaagcagggctatctgagccaggtcatccacg
agatcgtggacctgatgatccactaccaggccgtggtggtgctggagaacctgaatttcggctttaagagcaagaggaccggcatcg
ccgagaaggccgtgtaccagcagttcgagaagatgctgatcgataagctgaattgcctggtgctgaaggactatccagcagagaaa
gtgggaggcgtgctgaacccataccagctgacagaccagttcacctcctttgccaagatgggcacccagtctggcttcctgttttacgtg

cctgccccatatacatctaagatcgatcccctgaccggcttcgtggacccccttcgtgtggaaaaccatcaagaatcacgagagccgca
agcacttcctggagggcttcgactttctgcactacgacgtgaaaaccggcgacttcatcctgcactttaagatgaacagaaatctgtcctt
ccagaggggcctgcccggctttatgcctgcatgggatatcgtgttcgagaagaacgagacacagtttgacgccaagggcaccccctttc
atcgccggcaagagaatcgtgccagtgatcgagaatcacagattcaccggcagataccgggacctgtatcctgccaacgagctgat
cgccctgctggaggagaagggcatcgtgttcagggatggctccaacatcctgccaaagctgctggagaatgacgattctcacgccat
cgacacgatggtggccctgatccgcagcgtgctgcagatgcggaactccaatgccgccacaggcgaggactatatcaacagcccc
gtgcgcgatctgaatggcgtgtgcttcgactcccggtttcagaacccagagtggccaatggacgccgatgccaatggcgcctaccac
atcgccctgaagggccagctgctgctgaatcacctgaaggagagcaaggatctgaagctgcagaacggcatctccaatcaggact
ggctggcctacatccaggagctgcgcaac

dAsCas12a Ultra:
atgacacagttcgagggctttaccaacctgtatcaggtgagcaagacactgcggtttgagctgatcccacagggcaagaccctgaag
cacatccaggagcagggcttcatcgaggaggacaaggcccgcaatgatcactacaaggagctgaagcccatcatcgatcggatct
acaagacctatgccgaccagtgcctgcagctggtgcagctggattgggagaacctgagcgccgccatcgactcctatagaaagga
gaaaaccgaggagacaaggaacgccctgatcgaggagcaggccacatatcgcaatgccatccacgactacttcatcggccggac
agacaacctgaccgatgccatcaataagagacacgccgagatctacaagggcctgttcaaggccgagctgtttaatggcaaggtgc
tgaagcagctgggcaccgtgaccacaaccgagcacgagaacgccctgctgcggagcttcgacaagtttacaacctacttctccggc
ttttatgagaacaggaagaacgtgttcagcgccgaggatatcagcacagccatcccacaccgcatcgtgcaggacaacttccccaa
gtttaaggagaattgtcacatcttcacacgcctgatcaccgccgtgcccagcctgcgggagcactttgagaacgtgaagaaggccatc
ggcatcttcgtgagcacctccatcgaggaggtgttttccttcccttttataaccagctgctgacacagacccagatcgacctgtataacc
agctgctgggaggaatctctcgggaggcaggcaccgagaagatcaagggcctgaacgaggtgctgaatctggccatccagaaga
atgatgagacagcccacatcatcgcctccctgccacacagattcatcccccctgtttaagcagatcctgtccgataggaacaccctgtctt
tcatcctggaggagtttaagagcgacgaggaagtgatccagtccttctgcaagtacaagacactgctgagaaacgagaacgtgctg
gagacagccgaggccctgtttaacgagctgaacagcatcgacctgacacacatcttcatcagccacaagaagctggagacaatca
gcagcgccctgtgcgaccactgggatacactgaggaatgccctgtatgagcggagaatctccgagctgacaggcaagatcaccaa
gtctgccaaggagaaggtgcagcgcagcctgaagcacgaggatatcaacctgcaggagatcatctctgccgcaggcaaggagct
gagcgaggccttcaagcagaaaaccagcgagatcctgtcccacgcacacgccgccctggatcagccactgcctacaaccctgaa
gaagcaggaggagaaggagatcctgaagtctcagctggacagcctgctgggcctgtaccacctgctggactggtttgccgtggatga
gtccaacgaggtggaccccgagttctctgcccggctgaccggcatcaagctggagatggagccttctctgagcttctacaacaaggc
cagaaattatgccaccaagaagccctactccgtggagaagttcaagctgaactttcagaGgcctacactggcctctggctgggacgt
gaataaggagaagaacaatggcgccatcctgtttgtgaagaacggcctgtactatctgggcatcatgccaaagcagaagggcaggt
ataaggccctgagcttcgagcccacagagaaaaccagcgagggctttgataagatgtactatgactacttccctgatgccgccaaga
tgatcccaaagtgcagcacccagctgaaggccgtgacagcccactttcagacccacacaacccccatcctgctgtccaacaatttca
tcgagcctctggagatcacaaaggagatctacgacctgaacaatcctgagaaggagccaaagaagtttcagacagcctacgccaa
gaaaaccggcgaccagaagggctacagagaggccctgtgcaagtggatcgacttcacaagggattttctgtccaagtataccaaga
caacctctatcgatctgtctagcctgcggccatcctctcagtataaggacctgggcgagtactatgccgagctgaatcccctgctgtacc
acatcagcttccagagaatcgccgagaaggagatcatggatgccgtggagacaggcaagctgtacctgttccagatctataacaag
gactttgccaagggccaccacggcaagcctaatctgcacacactgtattggaccggcctgttttctccagagaacctggccaagaca
agcatcaagctgaatggccaggccgagctgttctaccgccctaagtccaggatgaagaggatggcacaccggctgggagagaag
atgctgaacaagaagctgaaggatcagaaaaccccaatccccgacaccctgtaccaggagctgtacgactatgtgaatcacagac
tgtcccacgacctgtctgatgaggccagggccctgctgcccaacgtgatcaccaaggaggtgtctcacgagatcatcaaggataggc
gctttaccagcgacaagttcttAttccacgtgcctatcacactgaactatcaggccgccaattccccatctaagttcaaccagagggtga
atgcctacctgaaggagcaccccgagacacctatcatcggcatcgcccggggcgagagaaacctgatctatatcacagtgatcgac
tccaccggcaagatcctggagcagcggagcctgaacaccatccagcagtttgattaccagaagaagctggacaacagggagaag

gagagggtggcagcaaggcaggcctggtctgtggtgggcacaatcaaggatctgaagcagggctatctgagccaggtcatccacg
agatcgtggacctgatgatccactaccaggccgtggtggtgctggagaacctgaatttcggctttaagagcaagaggaccggcatcg
ccgagaaggccgtgtaccagcagttcgagaagatgctgatcgataagctgaattgcctggtgctgaaggactatccagcagagaaa
gtgggaggcgtgctgaacccataccagctgacagaccagttcacctcctttgccaagatgggcacccagtctggcttcctgttttacgtg
cctgccccatatacatctaagatcgatcccctgaccggcttcgtggaccccttcgtgtggaaaaccatcaagaatcacgagagccgca
agcacttcctggagggcttcgactttctgcactacgacgtgaaaaccggcgacttcatcctgcactttaagatgaacagaaatctgtcctt
ccagaggggcctgcccggctttatgcctgcatgggatatcgtgttcgagaagaacgagacacagtttgacgccaagggcacccctttc
atcgccggcaagagaatcgtgccagtgatcgagaatcacagattcaccggcagataccgggacctgtatcctgccaacgagctgat
cgccctgctggaggagaagggcatcgtgttcagggatggctccaacatcctgccaaagctgctggagaatgacgattctcacgccat
cgacacgatggtggccctgatccgcagcgtgctgcagatgcggaactccaatgccgccacaggcgaggactatatcaacagcccc
gtgcgcgatctgaatggcgtgtgcttcgactcccggtttcagaacccagagtggccaatggacgccgatgccaatggcgcctaccac
atcgccctgaagggccagctgctgctgaatcacctgaaggagagcaaggatctgaagctgcagaacggcatctccaatcaggact
ggctggcctacatccaggagctgcgcaac

dLbCas12a:
Atgagcaagctggagaagtttacaaactgctactccctgtctaagaccctgaggttcaaggccatccctgtgggcaagacccaggag
aacatcgacaataagcggctgctggtggaggacgagaagagagccgaggattataagggcgtgaagaagctgctggatcgctact
atctgtcttttatcaacgacgtgctgcacagcatcaagctgaagaatctgaacaattacatcagcctgttccggaagaaaaccagaac
cgagaaggagaataaggagctggagaacctggagatcaatctgcggaaggagatcgccaaggccttcaagggcaacgagggct
acaagtccctgtttaagaaggatatcatcgagacaatcctgccagagttcctggacgataaggacgagatcgccctggtgaacagctt
caatggctttaccacagccttcaccggcttctttgataacagagagaatatgttttccgaggaggccaagagcacatccatcgccttcag
gtgtatcaacgagaatctgacccgctacatctctaatatggacatcttcgagaaggtggacgccatctttgataagcacgaggtgcagg
agatcaaggagaagatcctgaacagcgactatgatgtggaggatttctttgagggcgagttctttaactttgtgctgacacaggagggc
atcgacgtgtataacgccatcatcggcggcttcgtgaccgagagcggcgagaagatcaagggcctgaacgagtacatcaacctgta
taatcagaaaaccaagcagaagctgcctaagtttaagccactgtataagcaggtgctgagcgatcgggagtctctgagcttctacggc
gagggctatacatccgatgaggaggtgctggaggtgtttagaaacaccctgaacaagaacagcgagatcttcagctccatcaagaa
gctggagaagctgttcaagaattttgacgagtactctagcgccggcatctttgtgaagaacggccccgccatcagcacaatctccaag
gatatcttcggcgagtggaacgtgatccgggacaagtggaatgccgagtatgacgatatccacctgaagaagaaggccgtggtgac
cgagaagtacgaggacgatcggagaaagtccttcaagaagatcggctccttttctctggagcagctgcaggagtacgccgacgccg
atctgtctgtggtggagaagctgaaggagatcatcatccagaaggtggatgagatctacaaggtgtatggctcctctgagaagctgttc
gacgccgattttgtgctggagaagagcctgaagaagaacgacgccgtggtggccatcatgaaggacctgctggattctgtgaagag
cttcgagaattacatcaaggccttctttggcgagggcaaggagacaaacagggacgagtccttctatggcgattttgtgctggcctacg
acatcctgctgaaggtggaccacatctacgatgccatccgcaattatgtgacccagaagccctactctaaggataagttcaagctgtat
tttcagaaccctcagttcatgggcggctgggacaaggataaggagacagactatcgggccaccatcctgagatacggctccaagta
ctatctggccatcatggataagaagtacgccaagtgcctgcagaagatcgacaaggacgatgtgaacggcaattacgagaagatc
aactataagctgctgcccggccctaataagatgctgccaaaggtgttcttttctaagaagtggatggcctactataaccccagcgagga
catccagaagatctacaagaatggcacattcaagaagggcgatatgtttaacctgaatgactgtcacaagctgatcgacttctttaagg
atagcatctcccggtatccaaagtggtccaatgcctacgatttcaacttttctgagacagagaagtataaggacatcgccggcttttaca
gagaggtggaggagcagggctataaggtgagcttcgagtctgccagcaagaaggaggtggataagctggtggaggagggcaag
ctgtatatgttccagatctataacaaggacttttccgataagtctcacggcacacccaatctgcacaccatgtacttcaagctgctgtttga
cgagaacaatcacggacagatcaggctgagcggaggagcagagctgttcatgaggcgcgcctccctgaagaaggaggagctgg
tggtgcacccagccaactcccctatcgccaacaagaatccagataatcccaagaaaaccacaaccctgtcctacgacgtgtataag
gataagaggttttctgaggaccagtacgagctgcacatcccaatcgccatcaataagtgccccaagaacatcttcaagatcaataca
gaggtgcgcgtgctgctgaagcacgacgataacccctatgtgatcggcatcgccaggggcgagcgcaatctgctgtatatcgtggtg

gtggacggcaagggcaacatcgtggagcagtattccctgaacgagatcatcaacaacttcaacggcatcaggatcaagacagatta
ccactctctgctggacaagaaggagaaggagaggttcgaggcccgccagaactggacctccatcgagaatatcaaggagctgaa
ggccggctatatctctcaggtggtgcacaagatctgcgagctggtggagaagtacgatgccgtgatcgccctggaggacctgaactct
ggctttaagaatagccgcgtgaaggtggagaagcaggtgtatcagaagttcgagaagatgctgatcgataagctgaactacatggtg
gacaagaagtctaatccttgtgcaacaggcggcgccctgaagggctatcagatcaccaataagttcgagagctttaagtccatgtcta
cccagaacggcttcatcttttacatccctgcctggctgacatccaagatcgatccatctaccggctttgtgaacctgctgaaaaccaagt
ataccagcatcgccgattccaagaagttcatcagctcctttgacaggatcatgtacgtgcccgaggaggatctgttcgagtttgccctgg
actataagaacttctctcgcacagacgccgattacatcaagaagtggaagctgtactcctacggcaaccggatcagaatcttccgga
atcctaagaagaacaacgtgttcgactgggaggaggtgtgcctgaccagcgcctataaggagctgttcaacaagtacggcatcaatt
atcagcagggcgatatcagagccctgctgtgcgagcagtccgacaaggccttctactctagctttatggccctgatgagcctgatgctg
cagatgcggaacagcatcacaggccgcaccgacgtggattttctgatcagccctgtgaagaactccgacggcatcttctacgatagc
cggaactatgaggcccaggagaatgccatcctgccaaagaacgccgacgccaatggcgcctataacatcgccagaaaggtgctgt
gggccatcggccagttcaagaaggccgaggacgagaagctggataaggtgaagatcgccatctctaacaaggagtggctggagt
acgcccagaccagcgtgaagcac

dHyperLbCas12a:
Atgagcaagctggagaagtttacaaactgctactccctgtctaagaccctgaggttcaaggccatccctgtgggcaagacccaggag
aacatcgacaataagcggctgctggtggaggacgagaagagagccgaggattataagggcgtgaagaagctgctggatcgctact
atctgtcttttatcaacgacgtgctgcacagcatcaagctgaagaatctgaacaattacatcagcctgttccggaagaaaaccagaac
cgagaaggagaataaggagctggagaacctggagatcaatctgcggaaggagatcgccaaggccttcaagggcaacgagggct
acaagtccctgtttaagaaggatatcatcgagacaatcctgccagagttcctggacgataaggacgagatcgccctggtgaacagctt
caatggctttaccacagccttcaccggcttctttcgcaacagagagaatatgttttccgaggaggccaagagcacatccatcgccttca
ggtgtatcaacgagaatctgacccgctacatctctaatatggacatcttcgagaaggtggacgccatctttgataagcacgaggtgcag
gagatcaaggagaagatcctgaacagcgactatgatgtggaggatttctttgagggcgagttctttaactttgtgctgacacaggaggg
catccgcgtgtataacgccatcatcggcggcttcgtgaccgagagcggcgagaagatcaagggcctgaacgagtacatcaacctgt
ataatcagaaaaccaagcagaagctgcctaagtttaagccactgtataagcaggtgctgagcgatcgggagtctctgagcttctacg
gccggggctatacatccgatgaggaggtgctggaggtgtttagaaacaccctgaacaagaacagcgagatcttcagctccatcaag
aagctggagaagctgttcaagaattttgacgagtactctagcgccggcatctttgtgaagaacggccccgccatcagcacaatctcca
agcgcatcttcggcgagtggaacgtgatccgggacaagtggaatgccgagtatgacgatatccacctgaagaagaaggccgtggt
gaccgagaagtacgaggacgatcggagaaagtccttcaagaagatcggctccttttctctggagcagctgcaggagtacgccgacg
ccgatctgtctgtggtggagaagctgaaggagatcatcatccagaaggtggatgagatctacaaggtgtatggctcctctgagaagct
gttcgacgccgattttgtgctggagaagagcctgaagaagaacgacgccgtggtggccatcatgaaggacctgctggattctgtgaa
gagcttcgagaattacatcaaggccttctttggcgagggcaaggagacaaacagggacgagtccttctatggcgattttgtgctggcct
acgacatcctgctgaaggtggaccacatctacgatgccatccgcaattatgtgacccagaagccctactctaaggataagttcaagct
gtattttcagaaccctcagttcatgggcggctgggacaaggataaggagacagactatcgggccaccatcctgagatacggctccaa
gtactatctggccatcatggataagaagtacgccaagtgcctgcagaagatcgacaaggacgatgtgaacggcaattacgagaag
atcaactataagctgctgcccggccctaataagatgctgccaaaggtgttcttttctaagaagtggatggcctactataaccccagcga
ggacatccagaagatctacaagaatggcacattcaagaagggcgatatgtttaacctgaatgactgtcacaagctgatcgacttcttta
aggatagcatctcccggtatccaaagtggtccaatgcctacgatttcaacttttctgagacagagaagtataaggacatcgccggctttt
acagagaggtggaggagcagggctataaggtgagcttcgagtctgccagcaagaaggaggtggataagctggtggaggagggc
aagctgtatatgttccagatctataacaaggacttttccgataagtctcacggcacacccaatctgcacaccatgtacttcaagctgctgtt
tgacgagaacaatcacggacagatcaggctgagcggaggagcagagctgttcatgaggcgcgcctccctgaagaaggaggagc
tggtggtgcacccagccaactcccctatcgccaacaagaatccagataatcccaagaaaaccacaaccctgtcctacgacgtgtata
aggataagaggttttctgaggaccagtacgagctgcacatcccaatcgccatcaataagtgccccaagaacatcttcaagatcaata

cagaggtgcgcgtgctgctgaagcacgacgataacccctatgtgatcggcatcgccaggggcgagcgcaatctgctgtatatcgtgg
tggtggacggcaagggcaacatcgtggagcagtattccctgaacgagatcatcaacaacttcaacggcatcaggatcaagacagat
taccactctctgctggacaagaaggagaaggagaggttcgaggcccgccagaactggacctccatcgagaatatcaaggagctga
aggccggctatatctctcaggtggtgcacaagatctgcgagctggtggagaagtacgatgccgtgatcgccctggaggacctgaact
ctggctttaagaatagccgcgtgaaggtggagaagcaggtgtatcagaagttcgagaagatgctgatcgataagctgaactacatgg
tggacaagaagtctaatccttgtgcaacaggcggcgccctgaagggctatcagatcaccaataagttcgagagctttaagtccatgtct
acccagaacggcttcatcttttacatccctgcctggctgacatccaagatcgatccatctaccggctttgtgaacctgctgaaaaccaag
tataccagcatcgccgattccaagaagttcatcagctcctttgacaggatcatgtacgtgcccgaggaggatctgttcgagtttgccctg
gactataagaacttctctcgcacagacgccgattacatcaagaagtggaagctgtactcctacggcaaccggatcagaatcttccgg
aatcctaagaagaacaacgtgttcgactgggaggaggtgtgcctgaccagcgcctataaggagctgttcaacaagtacggcatcaa
ttatcagcagggcgatatcagagccctgctgtgcgagcagtccgacaaggccttctactctagctttatggccctgatgagcctgatgct
gcagatgcggaacagcatcacaggccgcaccgacgtggattttctgatcagccctgtgaagaactccgacggcatcttctacgatag
ccggaactatgaggcccaggagaatgccatcctgccaaagaacgccgacgccaatggcgcctataacatcgccagaaaggtgct
gtgggccatcggccagttcaagaaggccgaggacgagaagctggataaggtgaagatcgccatctctaacaaggagtggctgga
gtacgcccagaccagcgtgaagcac

HA-2xMYC_NLS:
ggatcctatccctatgacgtgcccgattatgccagcctgggcagcggctcccctgctgccaaacgcgttaaactagacgaagatcctg
ccgcgaagcgcgtgaaactggac

4xSID:
ggctccggaatgaacattcagatgttgctcgaggctgctgattacctggagcgccgcgaacgggaagcggaacacggttacgctagt
atgctgcctgggagcggcatgaatatccagatgctgctcgaggcggccgattatctcgaacgcagagagagagaagccgaacatg
gctacgcctctatgctgcctggctccgggatgaatatacagatgctcctggaagctgccgactatctggagcgaagagaaagggagg
ctgagcacgggtatgcaagtatgctccctggctccggtatgaatatccagatgctgctggaggccgctgactatcttgaacgccgaga
aagggaagccgagcatggatatgcatccatgctgccttctaggtctgcctacccatacgatgttccagattacgcttcgccgaagaaa
aagcgcaaggtc

VPR:
Gacgcattggacgattttgatctggatatgctgggaagtgacgccctcgatgattttgaccttgacatgcttggttcggatgcccttgatga
ctttgacctcgacatgctcggcagtgacgcccttgatgatttcgacctggacatgctgattaactctagaagttccggatctccgaaaaa
gaaacgcaaagttggtagccagtacctgcccgacaccgacgaccggcaccggatcgaggaaaagcggaagcggacctacgag
acattcaagagcatcatgaagaagtccccccttcagcggccccaccgaccctagacctccacctagaagaatcgccgtgcccagca
gatccagcgccagcgtgccaaaacctgccccccagccttaccccttcaccagcagcctgagcaccatcaactacgacgagttccct
accatggtgttccccagcggccagatctctcaggcctctgctctggctccagcccctcctcaggtgctgcctcaggctcctgctcctgcac
cagctccagccatggtgtctgcactggctcaggcaccagcacccgtgcctgtgctggctcctggacctccacaggctgtggctccacc
agcccctaaacctacacaggccggcgagggcacactgtctgaagctctgctgcagctgcagttcgacgacgaggatctgggagcc
ctgctgggaaacagcaccgatcctgccgtgttcaccgacctggccagcgtggacaacagcgagttccagcagctgctgaaccagg
gcatccctgtggcccctcacaccaccgagcccatgctgatggaataccccgaggccatcacccggctcgtgacaggcgctcagag
gcctcctgatccagctcctgcccctctgggagcaccaggcctgcctaatggactgctgtctggcgacgaggacttcagctctatcgccg
atatggatttctcagccttgctgggctctggcagcggcagccgggattccagggaagggatgtttttgccgaagcctgaggccggctcc
gctattagtgacgtgtttgagggccgcgaggtgtgccagccaaaacgaatccggccatttcatcctccaggaagtccatggccaacc
gcccactccccgccagcctcgcaccaacaccaaccggtccagtacatgagccagtcgggtcactgaccccggcaccagtccctca
gccactggatccagcgcccgcagtgactcccgaggccagtcacctgttggaggatcccgatgaagagacgagccaggctgtcaaa

gcccttcgggagatggccgatactgtgattccccagaaggaagaggctgcaatctgtggccaaatggacctttcccatccgcccccaaggggccatctggatgagctgacaaccacacttgagtccatgaccgaggatctgaacctggactcacccctgaccccggaattgaacgagattctggataccttcctgaacgacgagtgcctcttgcatgccatgcatatcagcacaggactgtccatcttcgacacatctctgttt

P300:
Attttcaaaccagaagaactacgacaggcactgatgccaactttggaggcactttaccgtcaggatccagaatcccttccctttcgtcaacctgtggaccctcagcttttaggaatccctgattactttgatattgtgaagagccccatggatctttctaccattaagaggaagttagacactggacagtatcaggagccctggcagtatgtcgatgatatttggcttatgttcaataatgcctggttatataaccggaaaacatcacgggtatacaaatactgctccaagctctctgaggtctttgaacaagaaattgacccagtgatgcaaagccttggatactgttgtggcagaaagttggagttctctccacagacactgtgttgctacggcaaacagttgtgcacaatacctcgtgatgccacttattacagttaccagaacaggtatcatttctgtgagaagtgtttcaatgagatccaaggggagagcgtttctttgggggatgacccttcccagcctcaaactacaataaataaagaacaattttccaagagaaaaaatgacacactggatcctgaactgtttgttgaatgtacagagtgcggaagaaagatgcatcagatctgtgtccttcaccatgagatcatctggcctgctggattcgtctgtgatggctgtttaaagaaaagtgcacgaactaggaaagaaaataagttttctgctaaaaggttgccatctaccagacttggcacctttctagagaatcgtgtgaatgactttctgaggcgacagaatcaccctgagtcaggagaggtcactgttagagtagttcatgcttctgacaaaaccgtggaagtaaaaccaggcatgaaagcaaggtttgtggacagtggagagatggcagaatcctttccataccgaaccaaagccctctttgcctttgaagaaattgatggtgttgacctgtgcttctttggcatgcatgttcaagagtatggctctgactgccctccacccaaccagaggagagtatacatatcttacctcgatagtgttcatttcttccgtcctaaatgcttgaggactgcagtctatcatgaaatcctaattggatatttagaatatgtcaagaaattaggttacacaacagggcatatttgggcatgtccaccaagtgagggagatgattatatcttccattgccatcctcctgaccagaagatacccaagcccaagcgactgcaggaatggtacaaaaaaatgcttgacaaggctgtatcagagcgtattgtccatgactacaaggatatttttaaacaagctactgaagatagattaacaagtgcaaaggaattgccttatttcgagggtgatttctggcccaatgttctggaagaaagcattaaggaactggaacaggaggaagaagagagaaaacgagaggaaaacaccagcaatgaaagcacagatgtgaccaagggagacagcaaaaatgctaaaaagaagaataataagaaaaccagcaaaaataagagcagcctgagtaggggcaacaagaagaaacccgggatgcccaatgtatctaacgacctctcacagaaactatatgccaccatggagaagcataaagaggtcttctttgtgatccgcctcattgctggccctgctgccaactccctgcctcccattgttgatcctgatcctctcatcccctgcgatctgatggatggtcgggatgcgtttctcacgctggcaagggacaagcacctggagttctcttcactccgaagagcccagtggtccaccatgtgcatgctggtggagctgcacacgcagagccaggac

KRAB:
Gatgctaagtcactgactgcctggtcccggacactggtgaccttcaaggatgtgtttgtggacttcaccagggaggagtggaagctgctggacactgctcagcagatcctgtacagaaatgtgatgctggagaactataagaacctggtttccttgggttatcagcttactaagccagatgtgatcctccggttggagaagggagaagagccctggctggtggagagagaaattcaccaagagacccatcctgattcagagactgcatttgaaatcaaatcatcagttccgaaaaagaaacgcaaagtt

3xKRAB:
Cggacccttgttactttcaaggatgtattcgtggatttcacacgcgaggagtggaaattgttggatactgcccagcagattgtataccgcaatgtaatgctcgaaaactacaaaaatctcgtgagtctcggatatcaactgaccaagcctgacgttatcctccgattggagaaaggcgaggaacctgggagcggaggaaggacgcttgtcacgtttaaggatgtttttgttgacttcactcgagaggagtggaagctgttggatactgcccaacaaattgtgtatcgcaacgtgatgctcgagaactacaagaaccttgtgtctttgggctatcagctgactaaacctgatgtaatactccgcctcgagaaaggagaggagcctggtagtgggggacggaccctggtaacattcaaggatgtattcgtggatttcacgcgggaagaatggaaactcctcgatactgcccaacagatagtgtatcgaaacgtaatgctcgaaaactacaaaaacttggtcagcctcggataccaacttacaaaacctgatgttatcctcagacttgagaagggcgaggaacccggtatccacggagtcccagcagccccaaagaagaagcggaaggtc

**Note:** For the below constructs, we find that there is strong mCherry signal and puromycin resistance when dCas12a and pre-crRNA constructs are transduced into cells, but not when they are transfected. However, others have had success when transfecting a similar construct (see Ref. 44)

mCherry-P2A-puroR-MALAT1-Lb_crRNA-Esp3I-cloning-site-KanR-Lb_crRNA-WPRE:
atggtgagcaagggcgaggaggataacatggccatcatcaaggagttcatgcgcttcaaggtgcacatggagggctccgtgaacg
gccacgagttcgagatcgagggcgagggcgagggccgcccctacgagggcacccagaccgccaagctgaaggtgaccaaggg
tggcccccctgcccttcgcctgggacatcctgtcccctcagttcatgtacggctccaaggcctacgtgaagcaccccgccgacatcccc
gactacttgaagctgtccttccccgagggcttcaagtgggagcgcgtgatgaacttcgaggacggcggcgtggtgaccgtgacccag
gactcctccctgcaggacggcgagttcatctacaaggtgaagctgcgcggcaccaacttcccctccgacggccccgtaatgcagaa
gaagaccatgggctgggaggcctcctccgagcggatgtaccccgaggacggcgccctgaagggcgagatcaagcagaggctga
agctgaaggacggcggccactacgacgctgaggtcaagaccacctacaaggccaagaagcccgtgcagctgcccggcgcctac
aacgtcaacatcaagttggacatcacctcccacaacgaggactacaccatcgtggaacagtacgaacgcgccgagggccgccac
tccaccggcggcatggacgagctgtacaagggatccggcgcaacaaacttctctctgctgaaacaagccggagatgtcgaagaga
atcctggaccgatgaccgagtacaagcccacggtgcgcctcgccacccgcgacgacgtccccagggccgtacgcaccctcgccg
ccgcgttcgccgactaccccgccacgcgccacaccgtcgatccggaccgccacatcgagcgggtcaccgagctgcaagaactctt
cctcacgcgcgtcgggctcgacatcggcaaggtgtgggtcgcggacgacggcgccgcggtggcggtctggaccacgccggagag
cgtcgaagcgggggcggtgttcgccgagatcggcccgcgcatggccgagttgagcggttcccggctggccgcgcagcaacagatg
gaaggcctcctggcgccgcaccggcccaaggagcccgcgtggttcctggccaccgtcggcgtgtcgcccgaccaccagggcaag
ggtctgggcagcgccgtcgtgctccccggagtggaggcggccgagcgcgccggggtgcccgccttcctggagacctccgcgcccc
gcaacctccccttctacgagcggctcggcttcaccgtcaccgccgacgtcgaggtgcccgaaggaccgcgcacctggtgcatgacc
cgcaagcccggtgcctaagaattcgattcgtcagtagggttgtaaaggtttttcttttcctgagaaaacaaccttttgttttctcaggttttgctt
tttggcctttccctagctttaaaaaaaaaaaagcaaaactcaccgaggcagttccataggatggcaagatcctggtattggtctgcgaa
atttctactaagtgtagatggagacgatagaagatcctttgatcttttctacggggtctgacgctcagtggaacgaaaactcacgttaagg
gattttggtcatgagattatcaaaaaggatcttcacctagatccttttaaattaaaaatgaagttttaaatcaatctaaagtatatatgagtaa
cctgaggctatggcagggcctgccgccccgacgttggctgcgagccctgggccttcacccgaacttgggggtggggtggggaaaa
ggaagaaacgcgggcgtattggccccaatggggtctcggtggggtatcgacagagtgccagccctgggaccgaaccccgcgtttat
gaacaaacgacccaacaccgtgcgttttattctgtctttttattgccgtcatagcgcgggttccttccggtattgtctccttccgtgtttcagtta
gcctccccctagggtgggcgaagaactccagcatgagatccccgcgctggaggatcatccagccggcgtcccggaaaacgattcc
gaagcccaacctttcatagaaggcggcggtggaatcgaaatctcgtgatggcaggttgggcgtcgcttggtcggtcatttcgaacccc
agagtcccgctcagaagaactcgtcaagaaggcgatagaaggcgatgcgctgcgaatcgggagcggcgataccgtaaagcacg
aggaagcggtcagcccattcgccgccaagctcttcagcaatatcacgggtagccaacgctatgtcctgatagcgatccgccacaccc
agccggccacagtcgatgaatccagaaaagcggccattttccaccatgatattcggcaagcaggcatcgccatgggtcacgacgag
atcctcgccgtcgggcatgctcgccttgagcctggcgaacagttcggctggcgcgagcccctgatgctcttcgtccagatcatcctgatc
gacaagaccggcttccatccgagtacgtgctcgctcgatgcgatgtttcgcttggtggtcgaatgggcaggtagccggatcaagcgtat
gcagccgccgcattgcatcagccatgatggatactttctcggcaggagcaaggtgagatgacaggagatcctgccccggcacttcgc
ccaatagcagccagtcccttcccgcttcagtgacaacgtcgagcacagctgcgcaaggaacgcccgtcgtggccagccacgatag
ccgcgctgcctcgtcttgcagttcattcagggcaccggacaggtcggtcttgacaaaaagaaccgggcgcccctgcgctgacagccg
gaacacggcggcatcagagcagccgattgtctgttgtgcccagtcatagccgaatagcctctccacccaagcggccggagaacctg
cgtgcaatccatcttgttcaatcatgcgaaacgatcctcatcctgtctcttgatcgatctttgcaaaagcctaggcctccaaaaaagcctc
ctcactacttctggaatagctcagaggccgaggcggcctcggcctctgcataaataaaaaaaattagtcagccatggggcggagaat
gggcggaactgggcggagttaggggcgggatgggcggagttaggggcgggactatggttgctgactaattgagatgcatgctttgca
tacttctgcctgctggggagcctggggactttccacacctggttgctgactaattgagatgcatgctttgcatacttctgcctgctggggag

cctggggactttccacaccctaactgacacacattccacagctggttctttccgcctcaggactcttcctttttcaatattattgaagcatttat
cagggttattgtctcatgagcggatacatatttgaatgtatttagaaaaataaacaaataggggttccgcgcacatttccccgaaaagtg
ccatatcgtctctatttctactaagtgtagataaagaattcgatatcaagctttctagataatcaacctctggattacaaaatttgtgaaagat
tgactggtattcttaactatgttgctccttttacgctatgtggatacgctgctttaatgcctttgtatcatgctattgcttcccgtatggctttcattttc
tcctccttgtataaatcctggttgctgtctctttatgaggagttgtggcccgttgtcaggcaacgtggcgtggtgtgcactgtgtttgctgacgc
aacccccactggttggggcattgccaccacctgtcagctcctttccgggactttcgctttccccctccctattgccacggcggaactcatc
gccgcctgccttgcccgctgctggacaggggctcggctgttgggcactgacaattccgtggtgttgtcggggaaatcatcgtcctttcctt
ggctgctcgcctgtgttgccacctggattctgcgcgggacgtccttctgctacgtcccttcggccctcaatccagcggaccttccttcccgc
ggcctgctgccggctctgcggcctcttccgcgtcttcgccttcgccctcagacgagtcggatctccctttgggccgcctccccgc

mCherry-P2A-puroR-MALAT1-Lb_crRNA-Esp3I-cloning-site-TruSeq_3’-WPRE:
atggtgagcaagggcgaggaggataacatggccatcatcaaggagttcatgcgcttcaaggtgcacatggagggctccgtgaacg
gccacgagttcgagatcgagggcgagggcgagggccgcccctacgagggcacccagaccgccaagctgaaggtgaccaaggg
tggcccccctgcccttcgcctgggacatcctgtcccctcagttcatgtacggctccaaggcctacgtgaagcaccccgccgacatcccc
gactacttgaagctgtccttccccgagggcttcaagtgggagcgcgtgatgaacttcgaggacggcggcgtggtgaccgtgacccag
gactcctccctgcaggacggcgagttcatctacaaggtgaagctgcgcggcaccaacttcccctccgacggccccgtaatgcagaa
gaagaccatgggctgggaggcctcctccgagcggatgtaccccgaggacggcgccctgaagggcgagatcaagcagaggctga
agctgaaggacggcggccactacgacgctgaggtcaagaccacctacaaggccaagaagcccgtgcagctgcccggcgcctac
aacgtcaacatcaagttggacatcacctcccacaacgaggactacaccatcgtggaacagtacgaacgcgccgagggccgccac
tccaccggcggcatggacgagctgtacaagggatccggcgcaacaaacttctctctgctgaaacaagccggagatgtcgaagaga
atcctggaccgatgaccgagtacaagcccacggtgcgcctcgccacccgcgacgacgtccccagggccgtacgcaccctcgccg
ccgcgttcgccgactaccccgccacgcgccacaccgtcgatccggaccgccacatcgagcgggtcaccgagctgcaagaactctt
cctcacgcgcgtcgggctcgacatcggcaaggtgtgggtcgcggacgacggcgccgcggtggcggtctggaccacgccggagag
cgtcgaagcgggggcggtgttcgccgagatcggcccgcgcatggccgagttgagcggttcccggctggccgcgcagcaacagatg
gaaggcctcctggcgccgcaccggcccaaggagcccgcgtggttcctggccaccgtcggcgtgtcgcccgaccaccagggcaag
ggtctgggcagcgccgtcgtgctccccggagtggaggcggccgagcgcgccggggtgcccgccttcctggagacctccgcgcccc
gcaacctccccttctacgagcggctcggcttcaccgtcaccgccgacgtcgaggtgcccgaaggaccgcgcacctggtgcatgacc
cgcaagcccggtgcctaagaattcgattcgtcagtagggttgtaaaggtttttcttttcctgagaaaacaaccttttgttttctcaggttttgctt
tttggcctttccctagctttaaaaaaaaaaaagcaaaactcaccgaggcagttccataggatggcaagatcctggtattggtctgcgaa
atttctactaagtgtagatggagacgatatatcgtctctTacgagatcggaagagcacacgtctgaactccagtcacaaagaattcgat
atcaagctttctagataatcaacctctggattacaaaatttgtgaaagattgactggtattcttaactatgttgctccttttacgctatgtggata
cgctgctttaatgcctttgtatcatgctattgcttcccgtatggctttcattttctcctccttgtataaatcctggttgctgtctctttatgaggagttgt
ggcccgttgtcaggcaacgtggcgtggtgtgcactgtgtttgctgacgcaacccccactggttggggcattgccaccacctgtcagctc
ctttccgggactttcgctttccccctccctattgccacggcggaactcatcgccgcctgccttgcccgctgctggacaggggctcggctgtt
gggcactgacaattccgtggtgttgtcggggaaatcatcgtcctttccttggctgctcgcctgtgttgccacctggattctgcgcgggacgt
ccttctgctacgtcccttcggccctcaatccagcggaccttccttcccgcggcctgctgccggctctgcggcctcttccgcgtcttcgccttc
gccctcagacgagtcggatctccctttgggccgcctccccgc
